# FACED 2.0 enables large-scale voltage and calcium imaging *in vivo*

**DOI:** 10.1101/2025.03.06.641784

**Authors:** Jian Zhong, Ryan G. Natan, Qinrong Zhang, Justin S. J. Wong, Christoph Miehl, Krishnashish Bose, Xiaoyu Lu, François St-Pierre, Su Guo, Brent Doiron, Kevin K. Tsia, Na Ji

## Abstract

Monitoring neuronal activity at large scale and high spatiotemporal resolution is crucial for understanding information processing within the brain. We optimized a kilohertz-frame-rate two-photon fluorescence microscope with all-optical megahertz line-scan rate to achieve ultrafast imaging across large areas and volumes at subcellular resolution. Applying this technique to voltage and calcium imaging *in vivo*, we demonstrated simultaneous recording of voltage activity over 200 neurons and calcium activity over 14,000 neurons.

## Main

Understanding the complex mechanisms of neural computation in the intact brain requires monitoring neuronal activity at subcellular or cellular spatial resolution over large areas/volumes with sufficient time resolution to capture physiological events of interest. For example, for calcium imaging, sampling rates of a few Hz are sufficient, whereas hundreds to kilohertz (kHz) sampling rates are required to capture signal associated with individual action potentials (APs).

Because of its ability to image deep inside the opaque mammalian brain and visualize neurons at sub-micron lateral resolution, point-scanning two-photon fluorescence microscopy^1^ (2PFM) is the most common approach for *in vivo* neuronal activity imaging. Here, fluorescence generation is confined to the focus of a high numerical aperture (NA) microscope objective, enabling optical sectioning in tissues. Images are generated by scanning the excitation focus across the sample and detecting emitted photons at each position. Therefore, the imaging speed of 2PFM is limited by how fast the excitation focus can be scanned. The mechanical inertia of galvanometric and resonant mirrors used in conventional 2PFM limits the line scan rate to 10’s of kilohertz (kHz), resulting in 10’s-100’s two-dimensional (2D) frames per second (fps) and even lower three-dimensional (3D) volume rates^2,3^. Although faster line scanning up to hundreds of kHz can be achieved by acousto-optic deflectors, these scanners have a more limited angular steering range and a smaller number of resolvable scanned points^4,5^. Recently, several methods utilized high-speed polygonal scanners and scan multiplier units for fast 2PFM imaging, but their line scan rates remain below 600 kHz^6–8^.

Previously, we reported a laser-scanning technique named free-space angular-chirp-enhanced delay (FACED)^9^ that achieved up to 4-megahertz (MHz) line scan rates and 3,000 fps for ultrafast 2PFM imaging *in vivo*, enabling 2D 1-kHz voltage imaging over up to a 50 µm × 250 µm field of view (FOV) with 6-s continuous data acquisition^10^. Here, we describe FACED 2.0 where improved hardware and software have enabled large-scale, continuous activity imaging in both 2D and 3D *in vivo*. Achieving a pixel rate of 1.0×10^8^ pixels per second while maintaining the spatial resolution and imaging depth of conventional 2PFM, we demonstrate high-throughput recording of cerebral blood flow, population voltage imaging of hundreds of neurons, 3D calcium imaging that densely sampled neurons in millimeter-scale volumes in the awake mouse brain, and whole-brain calcium imaging of zebrafish larvae in vivo.

Key optics in the FACED module are a cylindrical lens and a pair of highly reflective and quasi-parallel mirrors (reflectivity > 99.5% and misaligned angle α < 1 mrad; orange box, **Fig. 1a**; **Methods**). When placed between a laser and a microscope, the FACED module transforms a laser pulse into a set of time-delayed pulses propagating at slightly different angles. These pulses then form a 1D array of excitation foci that are spatially separated and temporally delayed (i.e., a line scan) at the objective focal plane. Without utilizing active scanning, the line scan rate of this all-optical and passive module equals the repetition rate of the laser, which can easily go beyond MHz^9,11^. A standard galvanometric scanning mirror (“Y galvo”) then scanned the pulse train along the orthogonal direction (magenta boxes, **Fig. 1a,b**) to obtain raster-scanning 2D images.

**Figure 1.**
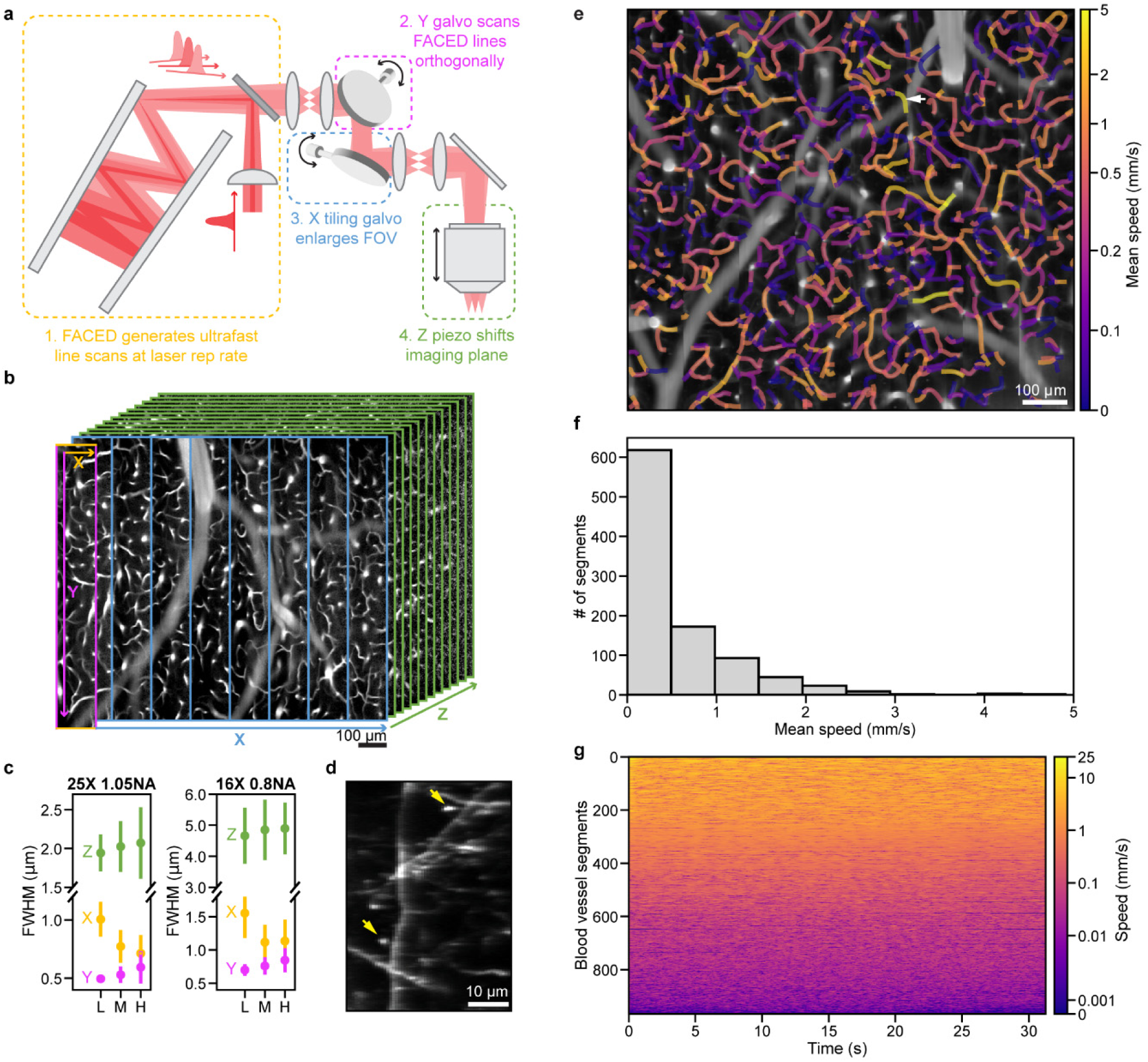
Schematics and performance of FACED 2.0. (**a**) Schematics of FACED 2.0. Colored boxes indicate scanning mechanisms of: 1, FACED module; 2, Y galvo scanning FACED lines perpendicularly to generate a FACED FOV; 3, X galvo tiling FACED FOV; 4, Z piezo scanning axially for volumetric imaging. (**b**) An 1,111 µm × 1,000 µm × 600 µm volume of mouse cortical vessels imaged at 624 ms/vol *in vivo*. Every 10^th^ images are shown. Colored lines and arrows correspond to scanning mechanisms in **a**. Voxel size: 1.4 µm × 2 µm × 5 µm. 1035 nm excitation at 240 mW post 0.8-NA objective. (**c**) Lateral (X: along FACED line; Y) and axial (Z) FWHMs measured from >500 200-nm-diameter beads for the first (L), middle (M), and last (H) third of FACED foci of 1.05-NA and 0.8-NA objectives, respectively. Error bars: standard deviation. (**d**) Maximum intensity projection of a 46 µm × 64 µm × 5 µm volume of dendrites at 36 µm depth in a Thy1-GFP line M mouse cortex. 1035 nm excitation at 150 mW post 1.05-NA objective. Arrows: putative synapses. (**e**) Time-averaged image of cortical blood vessels within a 1,111 µm × 1,000 µm FOV at 50 µm depth and 5.3 ms/frame *in vivo*. Pixel size: 1.4 µm × 2 µm. 1035 nm excitation at 215 mW post 0.8-NA objective. 967 capillary segments are color-coded by their mean blood flow speed. Arrow: capillary segment with a measured 25.9 mm/s maximal speed. (**f**) Histogram of mean blood speed of segments in **e**. (**g**) Temporal variations of flow speed in segments in **e**.

For FACED 2.0, we designed a FACED module that transformed each laser pulse from a 1-MHz excitation laser system of 920 nm or 1035 nm output into 100 pulses with a 1-ns inter-pulse interval in a 2PFM system constructed with off-the-shelf components (**Extended Data Fig. 1**; **Methods**). Scanning the Y galvo at 500 Hz with bidirectional data acquisition led to 1,000 fps over 140 µm × 1,035 µm FACED FOV (1.4 µm × 1.15 µm pixel sizes) for a 16× 0.8 NA objective and 80 µm × 585 µm FACED FOV (0.8 µm × 0.65 µm pixel sizes) for a 25× 1.05 NA objective. Scanning the Y galvo at 833 Hz enabled acquiring FOVs of comparable sizes at 0.6 ms/frame or 1,667 fps.

To enlarge the 2D FOV, FACED 2.0 utilized an additional X galvo to tile the kHz FACED FOVs laterally: for example, tiling 8 FACED FOVs laterally, we acquired a 1111 µm × 1000 µm FOV at 5.3 ms/frame with the 0.8 NA objective (blue boxes and arrow, **Fig. 1a,b**). To enable ultrafast volumetric imaging, we used a fast piezo stage to oscillate the objective axially (green boxes and arrow, **Fig. 1a,b**) – with the millisecond frame acquisition time of FACED 2.0, the continual Z movements during image acquisition did not generate in-frame motion artifacts (Supplementary Note 1). Upgrades in data acquisition hardware and software, including the implementation of field-programmable gate arrays (FPGA), enabled continuous streaming of data to disk at 0.256 GB/s (**Methods**).

Importantly, FACED 2.0 maintained subcellular resolution in XY and cellular resolution in Z (**Fig. 1c**), which are essential for resolving individual neurons in 3D and accurately measuring their activity. Even though field-dependent aberrations in the imaging setup caused modest variations in resolution (**Extended Data Figs. 2,3**), across the entire tiled FOV, 200-nm-diameter fluorescent beads had a lateral full width at half maximum (FWHM) of 0.83±0.19 µm in X (mean±SD) and 0.53±0.10 µm in Y when imaged by the 1.05 NA objective, and 1.27±0.38 µm in X and 0.76±0.14 µm in Y for the 0.8 NA objective. Axially, the FWHMs were 2.01±0.35 µm for the 1.05 NA objective and 4.79±0.91 µm for the 0.8 NA objective. Given that cortical neurons have cell bodies as small as 10 µm in size, with an axial FWHM less than 5 µm, FACED 2.0 had the resolution necessary for resolving single cells axially. Along the FACED line scans, the later foci had higher X resolution than the earlier foci (**Fig. 1c**; L versus H foci, p < 10^-47^ for both 16× 0.8 NA and 25× 1.05 NA objectives, Mann–Whitney U test), because their longer propagation distance led to a larger beam profile at the objective back pupil thus higher effective excitation NA. In a Thy1-GFP line M^12^ mouse cortex *in vivo*, the high lateral resolution enabled us to resolve dendritic spines with both 1.05 NA and 0.8 NA objectives (**Fig. 1d**, **Extended Data Fig. 4**). It also allowed the detection of neuronal processes and capillaries at 750 µm below dura (**Extended Data Figs. 5,6,7**).

We first applied FACED 2.0 to high-throughput measurement of cerebral blood flow in 2D. Using the 0.8 NA objective, we imaged a FOV of 1111 µm × 1000 µm with a pixel size of 1.4 µm × 2 µm at 5.3 ms/frame in the superficial cortex of an awake mouse (**Supplementary Video 1**). We retro-orbitally injected dextran-conjugated Rhodamine B to label the mouse blood plasma. Tracking the motion of unlabeled blood cells, we concurrently measured blood flow speed across 966 capillary segments (**Fig. 1e**) at a throughput (defined here as the FOV area imaged per millisecond) that was 18.7 times that of FACED 1.0^13^ and 5.2-fold higher than a recent study using another fast line-scanning technique^14^. Consistent with our previous result^13^, most segments had < 5 mm/s flow speed (**Fig. 1e,f**), but the high frame rate of FACED 2.0 enabled us to observe a maximal flow speed of 25.9 mm/s (white arrow, **Fig. 1e**). We observed minimal photobleaching during the 31.8-s recording, which allowed us to continuously monitor the temporally varying flow speed in these segments (**Fig. 1g**).

Next, we employed FACED 2.0 for population imaging of voltage activity from neurons expressing the genetically encoded voltage indicator JEDI-2P-kv^15^ in the awake mouse brain. Tiling two FACED FOVs, we used the 1.05 NA objective to image a 160 µm × 400 µm area with a pixel size of 0.8 µm × 0.8 µm at 1.25 ms/frame (Fig. 2).

**Figure 2.**
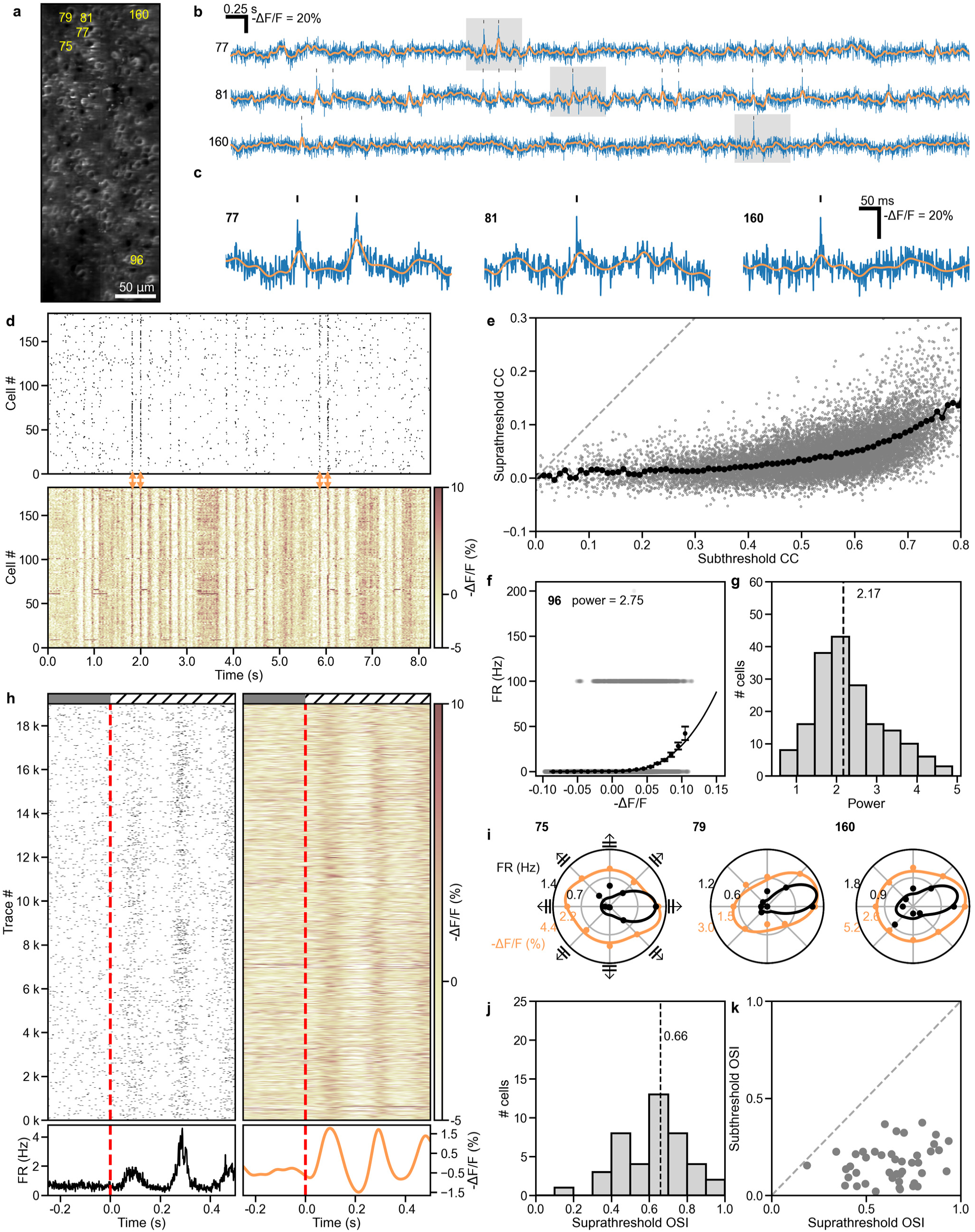
Imaging voltage activity from population of neurons *in vivo* at 1.25 ms/frame. (**a**) JEDI-2P-kv-expressing L2/3 neurons in a 160 µm × 400 µm FOV within a mouse visual cortex imaged at 132 µm depth and 1.25 ms/frame. Pixel size: 0.8 µm × 0.8 µm. 1035 nm excitation at 163 mW post 1.05-NA objective. Numbers: representative neurons. (**b**) Representative -ΔF/F voltage traces (in blue) of numbered neurons in **a**. Black ticks: suprathreshold spikes. Orange traces: subthreshold -ΔF/F. (**c**) Zoom-in view of voltage traces within the gray boxes in **b**. (**d**) Raster plots of (top) spikes and (bottom) subthreshold -ΔF/F traces of 182 neurons during a representative 8.25-s trial. (**e**) Suprathreshold versus subthreshold pairwise correlation coefficients (CC, gray dots) for 16,471 cell pairs from the 182 neurons in **a**. Black dots: binned averages with 0.01 bin size for subthreshold CC. Dashed gray line: reference line with slope = 1. (**f**) Plot of firing rate (FR) vs subthreshold -ΔF/F for Neuron 96, fit with a power-law function (black curve; power-law exponent = 2.75). Gray dots: FR vs subthreshold - ΔF/F. Black dots: binned average FR with 0.01 bin size in -ΔF/F. Error bar: s.e.m. (**g**) Histogram of fitted power-law exponent values of 182 neurons. Dashed line: median. (**h**) Spikes (left) and subthreshold responses (right) relative to stimulus onset (red dashed lines) extracted from 19,008 voltage traces of 88 neurons exhibiting visually evoked activity across 27 trials × 8 drifting grating orientations. Top: raster plots of individual traces; Bottom: Average firing rate (black curve) and average subthreshold -ΔF/F (orange curve). (**i**) Representative polar-plot tuning curves for supra- (black) and subthreshold (orange) voltage responses of Neuron 75, 79, 160, respectively. Dots: measured responses; curves: double-Gaussian fit. (**j**) Histogram of suprathreshold orientation-selectivity index (OSI) of 43 orientation selective (OS) neurons. Dashed line: median. (**k**) Scatter plot of subthreshold OSI vs. suprathreshold OSI for 43 OS neurons. Dashed line: reference line with slope = 1.

In one mouse, we simultaneously recorded voltage activity from 182 layer 2/3 visual cortical neurons (blue traces; representative traces from example neurons, **Fig. 2b**,**c**; representative traces from all neurons, **Extended Data Fig. 8**) in response to alternate 0.5-s blank screens and 0.5-s gratings (drifting in 8 directions from 0° to 315° at 45° intervals; 8.25 s per trial, 27 trials; **Methods**). During each 8.25-s trial, we observed on average 4% reduction in brightness due to photobleaching; Across the entire 222.75 s recording, the total photobleaching was around 15% (**Extended Data Fig. 9a,b**). We manually segmented each neuron’s soma membrane as the region of interest (ROI) and used VolPy^16^ to extract its voltage trace. We defined voltage transients as optically detected APs if they are detected as spikes by VolPy and also had their peak -ΔF/F larger than 2.5 times the standard deviations of the voltage trace within a 62-ms window centering on the spike (ticks, **Fig. 2b**; **Methods, Supplementary Note 2**). These spikes had peak -ΔF/F of ∼20%, consistent with the previously reported AP response of JEDI-2P-kv^15^. We extracted the subthreshold voltage responses by lowpass-filtering the voltage traces (orange traces, **Fig. 2b,c**).

Our ability to image voltage activity simultaneously from hundreds of neurons enabled experiments with throughput unachievable by electrophysiology. For example, intracellular whole-cell recording of APs and subthreshold membrane voltage is typically performed on one neuron at a time. Simultaneous whole-cell recordings from multiple neurons are possible but challenging to perform^17^, with up to four neurons^18–21^ simultaneously recorded *in vivo*. In comparison, voltage imaging by FACED 2.0 enabled us to simultaneously record spiking and subthreshold activity from these 182 neurons (representative 8.25-s trial, **Fig. 2d**).

One dominant feature of these data was the highly synchronized population-wide depolarization events in subthreshold activity in contrast to much less synchronized spiking (orange arrows, **Fig. 2d**). This characteristic was observed throughout all 27 trials for these 182 neurons (**Extended Data Fig. 10**). We calculated and compared the pairwise correlation coefficients for sub-versus suprathreshold activity (16,471 cell pairs from 182 neurons, **Fig. 2e**). Consistent with previous whole-cell recordings from brain slices^22^ and in vivo imaging^15^, we found that output (i.e., spiking) correlation increased with the increase in input (i.e., subthreshold) correlation, and that the input correlation was always substantially stronger than and bounded the correlation between the output spikes.

We frequently observed synchronized subthreshold 3-6 Hz oscillations lasting 1-2 s (**Fig. 2d**, **Extended Data Fig. 10**), in agreement with previous whole-cell^23,24^ and LFP^25^ recordings from visual cortex of the awake mouse. During these events, synchronized oscillatory firing was also observed from these neurons, consistent with extracellular recordings^26^ and likely resulting from thalamocortical interaction^24^.

Having access to both subthreshold activity and spiking also allowed us to investigate how individual neurons transform membrane potential V_m_ to spikes, by fitting a power-law function between the trial-averaged subthreshold input and firing rate^27,28^ (FR). Although ΔF/F from voltage imaging did not directly indicate V_m_ value, because the -ΔF/F response of JEDI-2P-kv is linearly proportional to V_m_ in the voltage range of -55 mV to -25 mV^15^, we could determine the exponent p in the power-law function FR=A·(-ΔF/F)^p^ for all 182 neurons, which ranged from 0.6 to 4.9 with a median value of 2.17 (example fit for ROI 96, **Fig. 2f**; Exponents for all 182 neurons, **Fig. 2g**). Previous electrophysiological recordings from the mouse brain produced exponent values within our measured range, albeit for much fewer numbers of neurons per mice and mostly acquired under anesthesia^29–32^.

From the spiking activity of the 182 neurons, we identified 88 neurons with visually evoked suprathreshold activity. Across 27 trials from these 88 neurons, we observed increased spiking and subthreshold activity following grating stimulus onsets as expected (**Fig. 2h**). We further identified 43 suprathreshold orientation-selective (OS) neurons within the imaging FOV. Consistent with prior electrophysiology recordings^29,32^, subthreshold depolarizations were evoked by a broader range of orientations than spiking (tuning curves of example neurons, **Fig. 2i**). We calculated the orientation selectivity index (OSI) using both supra- and subthreshold responses from the OS neurons. The distribution and median of the suprathreshold OSI (**Fig. 2j**) were similar to previous measurement from population calcium imaging of layer 2/3 neurons^33^. The sharper orientation tuning of the spiking compared with subthreshold responses was also reflected by their higher OSI values calculated from the OS population (**Fig. 2k**).

Tiling four FACED FOVs for each frame, we can double the imaging area to 320 µm × 400 µm at a pixel size of 0.8 µm × 0.8 µm and 2.6 ms/frame. Previous characterization on JEDI-2P dynamics indicates that imaging at 2.6 ms/frame should be able to capture most APs^15^. In one mouse, we recorded voltage activities from 225 layer 2/3 visual cortical neurons simultaneously (**Extended Data** Figs. 11-13), and observed low photobleaching (**Extended Data Fig. 9c,d**). Importantly, the population voltage dynamics observed at 2.6 ms/frame are highly similar to **Fig. 2** data acquired at 1.25 ms/frame, including activity correlation, input-output function, timing and magnitude of visually evoked responses, and population orientation tuning properties (out of 225 neurons in **Extended Data Fig. 11**, 155 neurons exhibited visually evoked suprathreshold activity, and 86 neurons displayed suprathreshold activity with orientation selectivity).

To test how the frame rate of imaging data affects the extracted voltage dynamics, we down-sampled the 800 Hz data in **Fig. 2** to 400 Hz and compared the activity characteristics from the down-sampled data with the original data (**Extended Data Fig. 14**). Consistent with the observations above, 400 Hz sampling can inform on neuronal activity with similar accuracy as 800 Hz in most application scenarios for JEDI-2P-kv and 800 Hz is only required when accurate recording of high-frequency firing is needed (**Supplementary Note 3**).

Finally, we demonstrated the ability of FACED 2.0 for high-throughput volumetric imaging, where its high 2D frame rate was used to rapidly record calcium activity at different depths by oscillating the objective axially. Importantly, FACED 2.0 does not sacrifice spatial resolution in exchange for imaging throughput, unlike other high-throughput volumetric imaging methods with reduced lateral and/or axial resolution (**Supplementary Table 1**). Using the 0.8 NA objective, we imaged a volume of 1111 µm × 1000 µm × 400 µm, at a voxel size of 1.4 µm × 2 µm × 10 µm and 260 ms/vol, within the visual cortex of an awake transgenic mouse with GCaMP6s^34^ expression in excitatory neurons (Slc17a7-IRES2-Cre^35^ × Ai162D^36^) in response to drifting grating stimuli (drifting in 8 directions from 0° to 315° at 45° intervals; 36 trials; **Fig. 3a**, **Supplementary Video 2**). The high lateral and axial resolutions of FACED 2.0 enabled cellular resolution imaging in 3D, while the voxel size allowed us to densely sample nearly all neurons within this volume.

**Figure 3.**
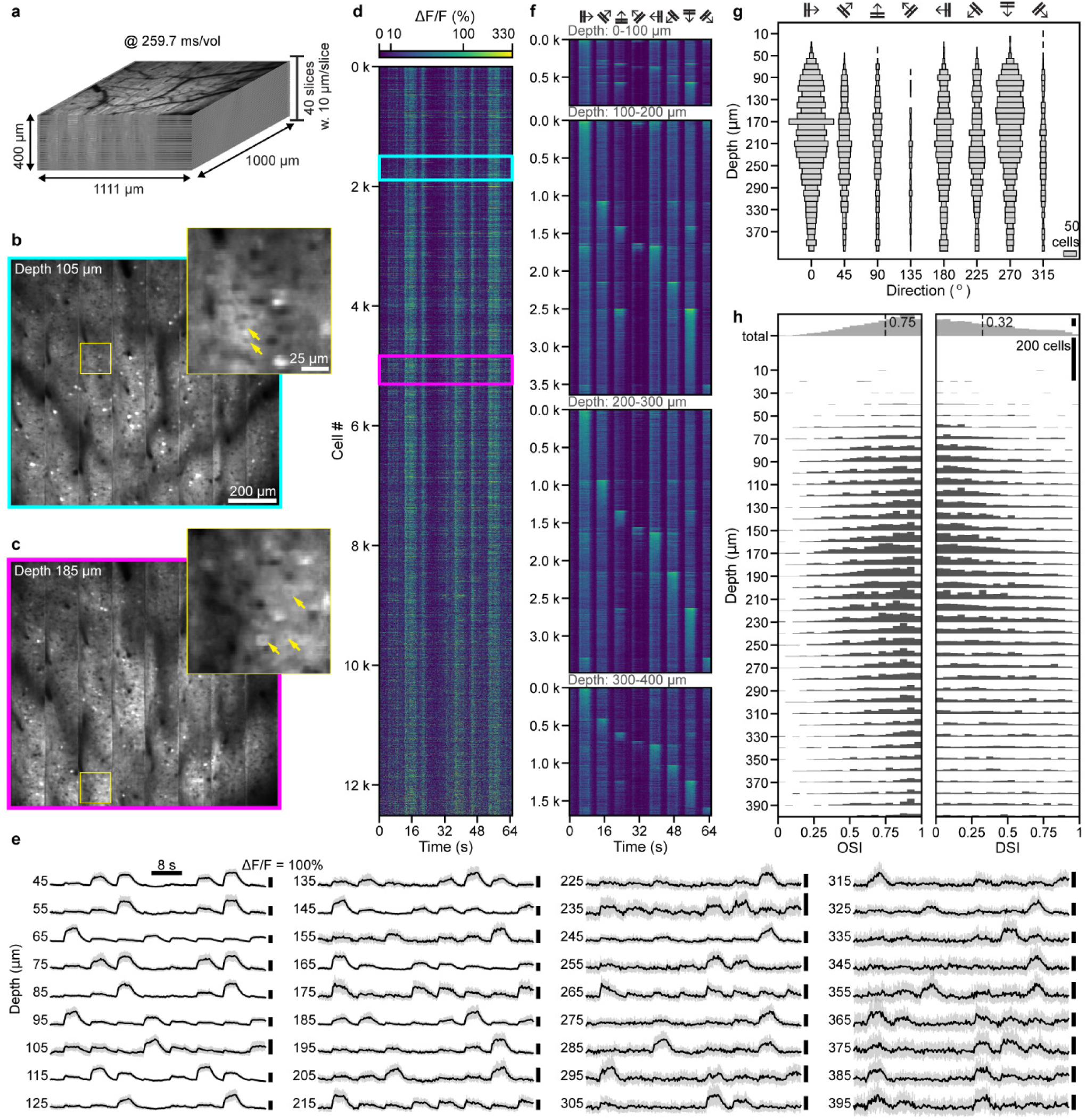
Large-scale in vivo volumetric imaging of calcium activity. (**a**) An 1,111 µm × 1,000 µm × 400 µm volume in the visual cortex of a Slc17a7-IRES2-Cre × Ai162D mouse imaged at 260 ms/vol. Voxel size: 1.4 µm × 2 µm × 10 µm. 920 nm excitation at 102 mW post 0.8-NA objective. (**b**,**c**) Example images at 105 µm and 185 µm depths. Insets: zoomed-in views of areas in yellow boxes. Arrows: neurons showing nucleus-excluded GCaMP6s expression. (**d**) Simultaneously recorded ΔF/F calcium traces from 12,511 neurons during a 64-s trial with drifting grating visual stimulation. Cyan and magenta boxes: calcium traces from neurons in b and c, respectively. (**e**) Example trial-averaged ΔF/F traces of neurons in **a** at 45 to 395 µm below dura. Gray shade: standard deviation. (**f**) 36-trial-averaged calcium traces from 9,811 neurons with visually evoked activity, grouped by depth 0-100 μm, 100-200 μm, 200-300 μm, and 300-400 μm. (**g**) Horizontal bars with their lengths indicating the numbers of neurons with their maximal ΔF/F at a specific grating direction for all 9,767 OS neurons across depths. (**h**) Histograms showing the OSI (left) and direction-selectivity index (DSI, right) distributions of OS neurons at each depth. Gray histogram (top): OSI and DSI distributions for all neurons. Dashed lines: median values.

Individual neurons were easily identified based on their morphology (**Fig. 3b,c**) and activity (**Supplementary Video 2**). We manually segmented 13,962 neurons from this dataset and identified 12,511 unique neurons based on their activity (ΔF/F traces for a single 64-s trial, **Fig. 3d**; trial-averaged traces for example neurons across the entire depth, **Fig. 3e**; entire 39-min recording, **Extended Data Fig. 15**). 9,811 neurons exhibited visually evoked activity (grouped by depths and ordered by their preferred directions, **Fig. 3e**), out of which we found 9,767 OS neurons. We identified the grating direction that evoked the maximal calcium response from each neuron with the 40 Z-planes and, consistent with previous reports^37–39^, observed a strong bias towards vertically and horizontally oriented gratings in layer 2/3 neurons (**Fig. 3f**). The OSI and direction selectivity index (DSI) distributions of OS neurons by depth (**Fig. 3g**) had similar median values (OSI: 0.75; DSI: 0.32) to those reported previously^33^. A comparison of OSI and DSI across depths (**Fig. 3g**) indicated that whereas OSI had mostly similar distributions across layer 2/3 and layer 4, neurons became more direction-selective (DS) at deeper depths, again consistent with previous reports^33,40^. Our ability to reach, from one imaged volume, similar conclusions on population feature selectivity to those acquired previously from many animals is a testament to the high throughput of FACED 2.0 in acquiring large-scale 3D calcium activity data.

Because FACED 2.0 maintains high excitation NA, it could image calcium activity deep in the mouse brain. In an awake wild-type mouse with its visual cortical neurons expressing GCaMP6s via viral transduction, we imaged a volume of 1111 µm × 1000 µm × 780 µm at a voxel size of 1.4 µm × 2 µm × 15 µm and 365.7 ms/vol (**Extended Data Fig. 16a**), allowing for the simultaneous recording of calcium activity during drifting grating stimulation from 14,005 unique neurons across the entire depth of the mouse visual cortex (**Extended Data Fig. 16b-g, Extended Data Fig. 17, Supplementary Video 3**). We identified 5,555 neurons with visually evoked activity (**Extended Data Fig. 16h**) down to ∼750 µm below dura (i.e., cortical layer 6^41^), where we observed OS and DS neurons (**Extended Data Fig. 16l**).

For animal models with smaller brains such as zebrafish larvae, FACED 2.0 is capable of performing simultaneous calcium imaging of multiple planes across the brain volume. Using the 1.05 NA objective, we imaged a 480 µm × 585 µm × 360 µm volume of a transgenic zebrafish larval brain (Tg[Elavl3:H2B-GCaMP6s]) with nucleus-targeted expression of GCaMP6s at a voxel size of 0.8 µm × 0.65 µm × 12 µm and 262.4 ms/vol (**Extended Data Fig. 18a, Supplementary Video 4**). We extracted activity traces from 4,718 spontaneously active neurons (**Extended Data Fig. 18b,c**). Switching to the 0.8 NA objective, we imaged a 555.2 µm × 800 µm × 360 µm volume with a voxel size of 1.4 µm × 1.6 µm × 15 µm at 96 ms/vol, over 2.6-fold faster than other methods^42–44^ for imaging zebrafish larval brains at similar volumes. Applying DeepCADRT^45^, a deep-learning-based denoising method to the volumetric images, we identified neuronal clusters with temporally distinct spontaneous activity across the brain volume (**Extended Data Fig. 19, Supplementary Video 5**).

In summary, FACED 2.0 achieved high-throughput imaging of neuronal activity over large scales. By scanning a high-NA focus at 1-MHz line-scan rate sequentially through the sample and detecting fluorescence with a non-imaging detector, it maintains the optical sectioning capability of conventional 2PFM and images deep within the opaque mouse brain. Importantly, FACED 2.0 achieves high-speed imaging without sacrificing spatial resolution, with its high excitation NA leading to subcellular resolution laterally and cellular resolution axially. As a result, compared with other high-throughput fluorescence imaging methods^7,8,10,14,46–53^, FACED 2.0 is unique in its combination of high throughput, high spatiotemporal resolution, and large imaging depths (**Supplementary Table 1**). Whereas either sparse labeling or sparse illumination is required for population voltage imaging utilizing single-photon excitation^54,55^, the intrinsic optical sectioning through two-photon absorption enables FACED 2.0 to be applicable to activity imaging in dense neuronal populations.

Most 2PFM systems utilize excitation lasers of ∼80 MHz repetition rate, where for each frame, fluorophores within a pixel are excited by multiple laser pulses with a ∼12.5-ns delay in between. In contrast, within one FACED frame, each pixel is only excited by one laser pulse. At 1,000 fps, subsequent excitation pulses arrive at the same sample position after 1-ms delays; hence FACED excitation provides ample time for fluorophores to return from their damage-prone dark states to their ground state. Such “dark state relaxation” has been shown to increase the photon yield in 2PFM by orders of magnitude^56^. By extracting more photons from individual fluorophores, FACED requires similar excitation power to conventional 2PFM but images at much higher frame rate while maintaining good signal-to-noise ratio (SNR).

Together, these characteristics enabled FACED 2.0 to overcome the optical constraints^57^ limiting the number of neurons that can be imaged simultaneously with more conventional 2PFM methods^5,58–61^, allowing for the first demonstration of population (defined here as >100 neurons) voltage imaging with 2P excitation^565,57–60^. With the voltage sensor JEDI-2P-kv, FACED 2.0 acquired fluorescence at high enough SNR to detect both subthreshold and suprathreshold activity from >200 neurons. From these data, we were able to measure the population correlation in both subthreshold and suprathreshold activity, characteristics that have been of longstanding interest^62,63^. We were also able to measure the power-law exponents of input-output transformation, important properties for developing and testing neural circuit models^27,28,64,65^, from hundreds of neurons in the awake mouse brain simultaneously.

JEDI-2P-kv has higher brightness and sensitivity than ASAP3^15^, which has enabled FACED 2.0 to simultaneously record voltage activity from more neurons than FACED 1.0 with ASAP3. With brighter and/or more sensitive activity indicators, fewer pixels per neuron would be needed to reach SNRs required for activity measurement, allowing larger pixel sizes thus further increasing area or volume and the number of neurons that can be imaged simultaneously. Therefore, the continuing advancements in fluorescence indicators are poised to further enhance the capabilities of FACED 2.0.

## Data availability

Raw structure images of Thy1-GFP mice can be found at https://figshare.com/s/fd97efe78295098ac37e. Raw blood flow and activity image data (> 1 TB) are available upon reasonable request to the correspondence author.

## Contributions

N. J. conceived of and supervised the project; J.Z. and N.J. designed the FACED module; J.Z. built the FACED module and microscope control system; J.Z. and J.W. built the data acquisition system; J.Z., R.G.N., Q.Z., and K.B. prepared samples; X.L. and F.S.P. provided reagents; J.Z. collected and analyzed data; C.M. and B.D. contributed to data analysis; K.K.T., and S.G. provided mentoring support; J.Z. and N.J. wrote the manuscript with input from all authors.

## Acknowledgements

We thank S. Olsen for discussion on electrophysiology data; J. Saulys and J. Zhu for help with the optical amplifier system; Y. Yang for help with sample preparation; G. Meng for discussion on blood flow speed measurements; J. Fan for help with visual stimulation setup; W. Chen for discussion on LabVIEW programming; J. Zhu for measuring pixel value to photon count conversion; and the reviewers for comments that led to materials in Supplementary Notes. This work was supported by NIH BRAIN Initiative grants UF1NS107696 (N.J.), U01NS118300 (N.J.), U19NS107613 (N.J.), R01NS109553 (N.J.), U01NS133971 (F.S.-P.), RF1NS128901 (F.S.-P); NIH grant R01EB032854 (F.S.-P), R01NS120219 (S.G.); a Klingenstein-Simons Fellowship Award in Neuroscience (F.S.-P.); the McNair Medical Foundation (F.S.-P.); Welch Foundation grants Q-2016-20190330 and Q-2016-20220331 (F.S.-P.); and Weill Neurohub (N.J.).

## METHODS

### Animals use

All animal experiments were conducted according to the National Institutes of Health guidelines for animal research. Procedures and protocols on mice and zebrafish were approved by the Institutional Animal Care and Use Committee at University of California, Berkeley.

### FACED 2.0

A simplified schematic of FACED 2.0 is presented in **Extended Data Fig. 1**. We used either the 1035 nm output of a fiber laser (Monaco 1035-40-40; Coherent) or the 920 nm output of an optical parametric amplifier (Opera-F; Coherent), both at 1 MHz repetition rate, for two-photon excitation. Before the FACED module, the 1035 nm beam was expanded to 5 mm in diameter (BE02M-B; Thorlabs) and the 920 nm beam to 4 mm in diameter (ACN254-040-B and AC254-050-B-ML; Thorlabs), followed by a further 3× expansion (BE03M-B; Thorlabs).

In the FACED module, the beam was focused in 1D (effective input numerical aperture 0.02, Δθ = 1.27°) by a cylindrical lens (LJ1267RM-B; Thorlabs) and then directed into a pair of nearly parallel mirrors (reflectivity > 99.9% and group delay dispersion < 40 fs² per reflection from 900 nm to 1050 nm, fused silica substrate, 250 mm long and 30 mm wide; Layertec). The mirror pair formed a small angle α = 0.0125° and had a separation distance of 150 mm. The focus of the cylindrical lens was located at the first mirror that the laser pulse encountered, after which the pulse reflected between the two mirrors, with the incidence angle reduced by α with each reflection. Because of 1D focusing by the cylindrical lens, different portions of the pulse had different initial incidence angles, thus underwent different numbers of reflections before their incidence angles became zero, when the light rays were retro-reflected. The light rays undergoing the same number of reflections between the mirrors formed a single output beamlet. Therefore, after retro-reflecting through the FACED mirrors, a single input pulse was split into a train of retro-reflected output pulses with distinct directions and time delays. A polarizing beam splitter (CCM1-PBS253, Thorlabs), in combination with a half-wave plate (AHWP10M-980, Thorlabs) and a quarter-wave plate (AQWP10M-980, Thorlabs), was employed to separate the input and output light of the FACED module. For FACED 2.0, we configured the optics to generate 100 output beamlets, with their propagation distances within the FACED module varying from ∼6 m to ∼36 m. To achieve precise angle alignment for the FACED mirrors, piezo actuators (PIA13; Thorlabs) were utilized to adjust the vertical and horizontal angles of the mirrors. The long propagation distance of light between the mirrors made the output sensitive to external perturbations such as laser pointing instability, temperature fluctuation, and mechanical vibration/drift. Depending on the stability of environment, realignment of the FACED mirrors using the piezo actuators needed to be performed every 5 to 30 minutes.

The spatially and temporally separated beamlets from the FACED module were then directed into a home-built 2PFM. A pair of achromatic doublets (AC508-500-B and AC508-250-B; Thorlabs) conjugated the focus of the cylindrical lens to the midpoint of a pair of closely and orthogonally arranged X and Y galvo mirrors (5-mm clear aperture, 6215H; Cambridge Technology). Subsequently, a scan lens pair (SL50-2P2 and TTL200MP; Thorlabs) conjugated the midpoint of the galvos to the back pupil plane of an objective lens that was mounted on a fast-moving piezo stage (SLC-1740-D-S; SmarAct). The total power throughput of the FACED module and the microscope is 9.3%. The objective, either 25× 1.05 NA (XLPLN25XWMP2; Olympus) or 16× 0.80 NA (N16XLWD-PF; Nikon), focused the excitation light and collected the two-photon excited fluorescence, which was reflected by a dichroic mirror (FF665-Di02-25×36; Semrock), focused by two lenses (AC508-080-A and LA1951-A; Thorlabs), filtered through an emission filter (F05-525/50-25 or FF01-680SP; Semrock), and finally detected by a photomultiplier tube (PMT) without a current protection circuit (H7442P-40-Y001, Hamamatsu). The output signal from the PMT was sampled at 10 Gigasamples per second (GSPS) using a high-speed digitizer (ADQ7-DC, Teledyne SP Devices).

FACED 2.0 module was designed to generate a line scan with 100 foci, spanning 80 µm and 140 µm for the 25× 1.05 NA and 16× 0.8 NA objectives, respectively. With the two mirrors separated by 150 mm, the time delay between adjacent foci was 1 ns. Additional analysis indicates that the choice of 1 ns delay does not cause detectable contamination in activity imaging (**Supplementary Note 4**, **Supplementary Fig. 7**). At 1-MHz laser repetition rate, the module operated at a line-scan rate of 1 MHz. By scanning the Y galvo orthogonally to the FACED line-scan direction at 833 Hz or 500 Hz and collecting data bidirectionally, we achieved an imaging speed of 0.6 ms/frame or 1 ms/frame. Here, each frame had 100 pixels along the FACED/X scan axis and up to 900 pixels along the Y axis. To extend the field of view (FOV) along X, we stepped the X galvo to tile multiple FACED FOVs. The maximal achievable FOV for our current system was 640 µm × 585 µm with a 25× 1.05 NA objective and 1,111 µm × 1,035 µm with the 16× 0.8 NA objective, limited by the maximal scanning angle of the galvo scanners and the focal length of the objective. If desired, volumetric imaging was accomplished by axially scanning the objective with the fast piezo stage, while keeping the excitation power constant. The imaging parameters and estimation of number of photons are summarized in **Supplementary Tables 2-4**. **Supplementary Note 5** provides additional discussion on how FACED 2.0 performs compared with FACED 1.0.

The 1D signal waveform recorded at 10 GSPS by the digitizer was further processed by an onboard FPGA which averaged every 10 sampling points as the fluorescence signal excited by the preceding FACED focus. The resulting data were first stored in the digitizer’s onboard memory and then transmitted to a data acquisition (DAQ) computer through a PCIe 3.0 ×8 interface at 0.256 GB/s. A custom multi-thread data acquisition program optimized data transfer and storage scheme, enabling continuous data streaming from the digitizer to the computer. Our DAQ system allows uninterrupted data collection, limited in its duration only by disk storage size.

### Point spread function (PSF) measurements

PSF measurements (**Extended Data** Figs. 2,3) were performed using 0.2-µm-diameter red fluorescent beads (Invitrogen™ F8786) positioned in the focal plane of the microscope objective. Using the galvo mirrors and the objective piezo stage, we moved the FACED focal array relative to the fluorescent beads in 3D and recorded the fluorescence signal at each position. For the 25× 1.05 NA objective, step sizes of 0.1 µm in X, 0.05 µm in Y, and 0.2 µm in Z were utilized. For the 16× 0.8 NA objective, step sizes of 0.15 µm in X, 0.05 µm in Y, and 0.5 µm in Z were used. We then measured the full widths at half maximum (FWHM) of the PSF profiles excited by each FACED focus along the X, Y, and Z axes. The same process was repeated for beads located along the horizontal, vertical, and the two diagonal directions of the entire FOV.

### Mice preparations

All experimental mice were purchased from Jackson Laboratories, including wildtype (C57BL/6J, stock no. 000664), Slc17a7-IRES2-Cre-D (stock no. 037512) × Ai162D (stock no. 031562) transgenic, and Thy1-GFP line M transgenic mice (stock no. 007788), and bred in house. All mice were >2-month old at the time of cranial window installation and housed under a reverse light cycle.

Virus injection and cranial window implantation procedures have been described previously^1^. Briefly, mice were anesthetized with isoflurane (1–2% by volume in oxygen) and given the analgesic buprenorphine (subcutaneously, 0.3 mg/kg of body weight). Animals were head-fixed in a stereotaxic apparatus (Model 1900; David Kopf Instruments). A 3.5-mm-diameter craniotomy was created over intact dura of the left visual cortex. For wildtype mice, virus injection was performed to express genetically encoded calcium and voltage sensors using a glass pipette (Drummond Scientific Company) beveled at a 35° angle with a 15–20-μm opening and back-filled with mineral oil. A fitted plunger controlled by a hydraulic manipulator (MO10; Narishige) was inserted into the pipette for loading and slow injection of the virus into 6-12 sites within the left visual cortex. At each injection site, 30 nl AAV2/1.syn.GCaMP6s solution (1.8 × 10^13^ GC/ml) was injected at 250 µm and 500 µm below the pia for calcium imaging and 50 nl pAAV-hSyn-JEDI2P-Kv2.1 solution (1.17 × 10^11^ GC/ml) was injected at 250 µm below pia for voltage imaging. A glass window made of a single coverslip (No. 1.5; Fisher Scientific) was then embedded in the craniotomy and sealed in place using dental acrylic. Subsequently, a titanium head-post was affixed to the skull using cyanoacrylate glue and dental acrylic. *In vivo* imaging was conducted after >2 weeks of recovery and/or virus expression, as well as habituation to head fixation. All imaging experiments were carried out on head-fixed awake mice.

### Zebrafish preparation for *in vivo* imaging

Nacre zebrafish larvae expressing Tg[Elavl3:H2B-GCaMP6s] were imaged on 6 dpf and 7 dpf. Each larva was mounted dorsal side up on a 35-mm polystyrene petridish using 2% low melting agarose. Throughout the imaging experiment, the larvae were maintained in E3 medium (5 mM NaCl, 0.17 mM KCl, 0.33 mM CaCl2, 0.33 mM MgSO4) at room temperature.

### Dye injection for blood vessel imaging in the mouse brain

Prior to imaging, animals were briefly anesthetized with isoflurane and retro-orbitally injected with 50 µl of 5% (w/v) 70k MW dextran-conjugated Rhodamine B fluorescent dye (D1841; ThermoFisher).

### Visual stimulation in head-fixed awake mice

Visual stimuli (Psychophysics Toolbox^2,3^) were presented on a liquid-crystal display (22-inch diagonal and 1,920 × 1,080 pixels; Dell P2219H), which was positioned ∼15 cm from the mouse’s right eye and oriented at ∼40° to the long body axis of the mice. Blue drifting sinusoidal gratings (100% contrast, 0.06 cycles per degree, 2Hz temporal frequency) were generated with Psychophysics Toolbox. For voltage imaging, each stimulus consisted of a 0.5-s blank screen followed by a 0.5-s drifting grating. For calcium imaging of the GCaMP6s transgenic mice, each stimulus included a 4-s blank screen followed by a 4-s drifting grating. For calcium imaging of wild-type mice with viral transduction of GCaMP6s, each stimulus included a 3-s blank screen, a 3-s drifting grating, and then a 2-s blank screen. Image sequences were acquired throughout each trial that spanned 8 stimuli of different gratings with drifting directions ordered from 0° to 315° at 45° intervals, where 0° corresponded to a vertical grating moving posteriorly and 90° to a horizontal grating moving upward. Visual stimulation computer generated a trigger signal to the microscope to start data acquisition at the beginning of each trial. A total of 16 to 51 trials were repeated for each experiment, determined by the available disk space or the SNR of single-trial data. Even though 36 (**Fig. 3**) and 51 trials (**Extended Data Fig. 16**) were collected for calcium imaging, 10 trials were sufficient for characterizing visually evoked activity, as shown for neurons randomly selected across the imaging depth (**Extended Data Fig. 20**).

### Analysis of cerebral blood flow in the mouse brain

Image sequences first underwent motion registration using CaImAn^4^. Line regions of interest (ROIs) with relatively uniform brightness and representing continuous blood vessel segments were manually selected. For each ROI, the image sequence was resliced along its length to create a kymograph. To reduce brightness variation along the length of the blood vessel segment, pixel values in the raw kymograph was normalized/divided by the temporally averaged signal profile of the corresponding ROI. A Radon-transform-based method^5^ was used to extract the temporally varying flow speed from the kymographs. On average in 9% of the imaging duration, artifactual results characterized by unphysiologically rapid and large oscillations of flow speed were observed. We adopted the following procedure to identify and mitigate these artifacts. First, we identified data points that fell outside the window defined by the mean±3×SD (standard deviation) of a 5-timepoint rolling window and replaced them with the mean value. Then we applied a Kalman filter^6^, which estimates states of a dynamic system by combining system modeling and actual measurements, to further reduce noise in the flow speed measurement. Here, we modeled the blood flow speed as:

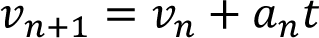

Here *t* is the temporal interval, ν_*n*_ and *a_n_* were the blood flow speed and acceleration at the *n*th time point, respectively, and ν_*n*+1_ was the blood flow speed at (*n*+1)th time point. The transition matrix *F*, observation matrix *H*, and initial state *X*_0_ of the Kalman filter were then written as follows:

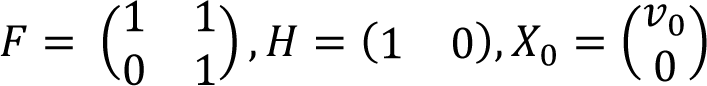

Here, ν_0_ was the blood flow speed at the very beginning of the imaging session. Other parameters of the Kalman filter were then automatically estimated using the expectation-maximization algorithm^7^ and data processing were completed using pykalman^8^.

### Analysis of voltage imaging datasets

Image sequences first underwent motion registration using CaImAn^4^. ROIs were manually segmented to encompass soma membrane. Image sequences and ROI masks were inputted into SVolPy^9^, which extracted the voltage traces ΔF/F (“dFF” output from Volpy) and detected spikes. From spikes identified by VolPy, we further rejected spikes with its peak -ΔF/F value lower than the mean+2.5×SD within a 20-frame rolling window in the corresponding voltage trace. Subthreshold ΔF/F was computed by applying a 20-Hz 5^th^-order Butterworth lowpass filter to the voltage trace ΔF/F.

We analyzed suprathreshold correlation versus subthreshold correlation, following procedures reported previously^10^. To mitigate the effects of spike frequency adaptation^10^, the initial 100 ms of each trial was excluded. Subsequently, 40-ms rolling windows were applied to both the detected spikes and the subthreshold response to calculate the firing rate and corresponding subthreshold -ΔF/F, with the firing rate computed by dividing the number of spikes by the window length and the subthreshold -ΔF/F determined by averaging the values within the window. The shift-corrected covariance of the responses between each pair of cells was then calculated as^10^:

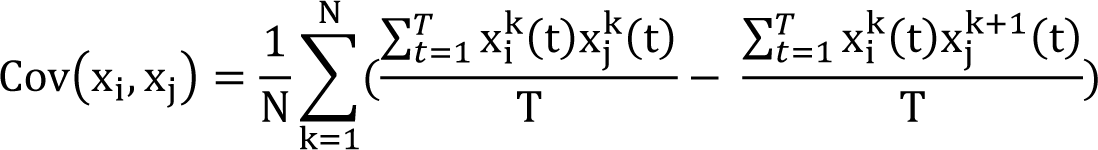

Here, N was the total number of trials (i.e., 27 in **Fig. 2**), T was the duration of each trial (i.e., 8.25 s from the 1^st^ to the last frame of each trial in **Fig. 2**), x_i_^k^(t) represented either the firing rate or the subthreshold -ΔF/F of neuron i at time t in the k^th^ trial. The sums over k = 1…N and t represented summation over N trials and time, respectively. We used rolling notation for trial numbers (i.e., the (N+1)^th^ trial equals the first trial with x_i_^N+1^ = x_i_^1^). Lastly, the correlation coefficient between neuron i and neuron j was computed as:

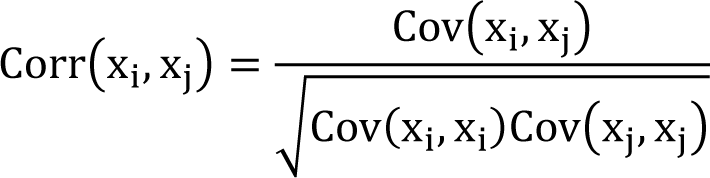

In **Fig. 2e**, the black symbols/line were calculated by averaging the suprathreshold correlation coefficients within subthreshold correlation coefficient bins of 0.01 in width.

For the analysis of neuronal input/output functions, 10-ms rolling windows were employed to calculate the firing rate^10^ (FR) and the corresponding subthreshold response of each neuron, with the firing rate determined by dividing the number of spikes by the window length and the corresponding subthreshold response computed by averaging the subthreshold -ΔF/F within the window. Subsequently, the firing-rate data points within bins of subthreshold -ΔF/F (bin width: 0.01) were averaged to generate the input-output curve (black symbols and curve, **Fig. 2f**), which was then fitted using a power-law expression to determine the amplitude A and power-law exponent p:

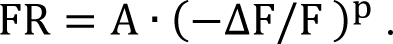

For the analysis of visually evoked activity, we defined the pre-stimulus period as the 250 ms before the onset of grating stimulus. We defined the response window to span the 750 ms after the onset of grating stimulus. We calculated the firing rates during pre-stimulus period and response window by dividing the numbers of spikes with their duration (i.e., 250 ms for pre-stimulus period, 750 ms for response window). Neurons exhibiting a significant difference in their firing rates between the pre-stimulus periods and response windows for at least one grating drifting direction (Holm-Bonferroni multiple comparison-corrected T test, P < 0.05) were defined as exhibiting visually evoked activity. Visually evoked neurons that demonstrated a significant difference in firing rates during the response windows across different grating stimuli (one-way ANOVA, P < 0.05) were defined as orientation-selective (OS) neurons.

In analyzing the suprathreshold tuning properties of OS neurons, we first calculated the mean spontaneous firing rate of each neuron by averaging its firing rates across all the pre-stimulus periods of all trials. We then calculated the neuron’s suprathreshold response towards each grating orientation by subtracting the mean spontaneous firing rate from the trial-averaged firing rate within the response window. The subthreshold response towards each grating orientation was determined by subtracting the mean subthreshold -ΔF/F in the pre-stimulus period for each orientation from the maximum trial-averaged subthreshold -ΔF/F in the corresponding response window. Tuning curve fitting and orientation selectivity index calculations followed the method in the **tuning analysis** section below.

### Analysis of volumetric calcium imaging datasets from the mouse visual cortex

The image sequences first underwent motion registration using CaImAn^4^. ROIs were manually segmented to encompass the somata of neurons in each Z image plane. Time-series images at each Z depth, along with the ROI masks, were processed using Suite2p^11^ to compute the fluorescence traces of each ROI (average of all pixels within, F_roi_(*t*)) and its associated neuropil. Within Suite2p, the neuropil masks were iteratively generated through a process of expanding each edge of the ROI mask by a margin of 5 pixels and then removing pixels that were associated with neighboring neurons. This iterative expansion continued until the resulting neuropil mask contained a minimum of 340 pixels. The average of all pixels within the neuropil mask was calculated as F_neu_(*t*).

We subtracted the neuropil transient ΔF_neu_(*t*) from F_roi_ to reduce neuropil contamination:

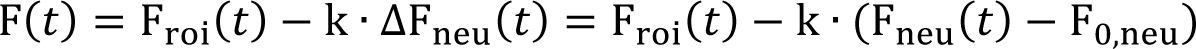

Here F_0,neu_ was the baseline fluorescence of the neuropil, which was the averaged neuropil signal within the 1-s blank screen periods before stimulus onset across all trials. We chose the neuropil subtraction coefficient k to be 0.7^12^. The calcium transients ΔF/F for each ROI was subsequently computed with ΔF/F = (F(*t*) − F_0_)/F_0_, where F_0_ was the averaged F signal within the 1-s blank screen periods before stimulus onset across all trials.

Because a neuron may show up in neighboring Z planes, we used a clustering algorithm to identify ROIs that corresponded to the same neuron. Initially, the algorithm traversed images at various depths, linking ROIs from adjacent Z depths with overlapping XY pixel coordinates into a ROI chain. Such a ROI chain encompassed all ROIs that were potentially affiliated with the same neuron. In instances where an ROI had multiple overlapping ROIs within the same neighboring Z plane, the algorithm selected the overlapping ROI whose ΔF/F had the highest Pearson correlation coefficient with that of the current ROI. Following the formation of ROI chains, the algorithm clusters the ROIs within each chain, ensuring that the ROIs within every cluster have 3D distances <25 µm and calcium-trace Pearson correlation coefficients >0.5 with one another. ROIs contained within the same ROI cluster were then identified as belonging to the same neuron, and the ROI with the highest SNR was selected to represent this neuron.

In the analysis of the visually evoked suprathreshold activity, the pre-stimulus period was defined as the 1 s before the onset of grating stimulus, while the response window was selected to extend from the onset of the grating stimulus to the start of the initial blank-screen period of the subsequent stimulus. Neurons exhibiting a significant difference between the time-averaged ΔF/F during the pre-stimulus periods and response windows for at least one grating orientation (Bonferroni multiple comparison-corrected T test, P < 0.05) were defined as exhibiting visually evoked activity. Visually evoked neurons with significantly different time-averaged ΔF/F during the response windows across different orientations (one-way ANOVA, P < 0.05) were defined as OS. The visually evoked response of a neuron towards a specific grating stimulus was calculated as the time-averaged (over the response window) and trial-averaged ΔF/F value. Tuning curve fitting and calculations for orientation and direction selectivity indices followed the **tuning analysis** section below.

For spontaneously active neurons, baseline fluorescence was recalculated using the mode (i.e., most frequent value) of the fluorescence intensity histograms of F_neu_(*t*) and F(*t*). The procedures for neuropil subtraction and ΔF/F calculations followed the procedures described above.

### Analysis of volumetric calcium imaging datasets from zebrafish larval brains

The image sequences first underwent motion registration using CaImAn^4^. ROIs were manually segmented to encompass nuclei of neurons in images captured at each Z plane. The fluorescence trace F_roi_(*t*) was computed from the pixel average within the ROI. We then low-pass-filtered F_roi_(*t*) (Gaussian filter with σ = 2 frames) and plotted its values into a fluorescence intensity histogram, which may exhibit multiple peaks. We located the peak with the lowest fluorescence intensity value and defined its value as the baseline fluorescence F_0_. We then calculated the calcium transients of zebrafish neurons as ΔF/F = (F_roi_(*t*) − F_0_)/F_0_.

For the dataset acquired with the 25× 1.05 NA objective, the motion-registered raw dataset was directly used. The clustering algorithm described in the analysis of the mouse visual cortex volumetric calcium imaging dataset was implemented to identify ROIs belonging to the same neuron. The clustering threshold for the zebrafish dataset was chosen such that all ROIs within the same cluster had 3D distances < 20 µm and calcium-trace Pearson correlation coefficients > 0.5. ROIs contained within the same ROI cluster were then identified as belonging to the same neuron, and the ROI with the largest SNR was selected to represent this neuron.

The dataset acquired with a 16× 0.8 NA objective first underwent denoising using DeepCAD-RT^13^. At each Z depth, baseline fluorescence F_0_ of each pixel was obtained following the baseline calculation process described above for zebrafish ROIs. ΔF(t) = F(*t*) − F_0_ was then calculated for each pixel at each time point, generating an image time series for ΔF. We then color-coded each ΔF(t) image based on its time t (using “gist_rainbow” color map in Matplotlib in Python), then sum-projected the entire time series to generate the temporal color-coded projection for each Z depth (**Extended Data Fig. 19**), where the color indicated the most active period for each pixel.

### Tuning analysis

For orientation tuning analysis, we fitted the response R and the orientation angle (θ in degrees) with the following bimodal Gaussian function^14^:

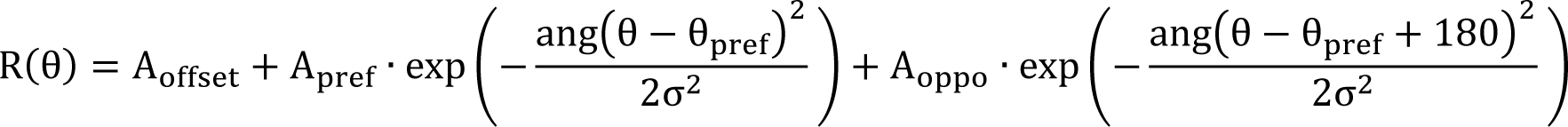

Here θ_pref_ was the preferred orientation, A_offset_ was a constant offset, and A_pref_ and A_oppo_ were the amplitudes at θ_pref_ and θ_pref_ − 180 degree, respectively. ang(x) = min (x, 360 − x, 360 + x) wrapped angular values onto the interval 0° to 180°. After fitting, the orientation-selectivity index (OSI) and the direction selectivity index (DSI) were calculated using the following formulas:

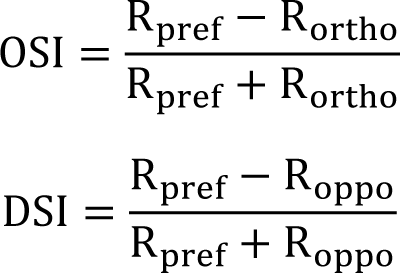

Where R_pref_, R_ortho_, and R_oppo_ are the responses calculated from the expression for R(θ) at θ_pref_, θ_pref_ − 90 and θ_pref_ − 180, respectively.

### Image presentation in figures and videos

Although activity analysis was carried out on motion-registered raw images as described above, images presented in figures and videos underwent normalization to account for the uneven excitation within the FACED line scan. This is because from the focus with the shortest to the focus with the longest propagation distance, the focal power decreases (because of power loss at each mirror reflection) while the effective excitation NA increases (because of increase in beam size with longer propagation) gradually. In addition, the Gaussian intensity profile of the input pulse also leads to higher energy foci in the middle of the line scan. As a result, for a uniform fluorescence sample, the 2P signal along the FACED line scan is brightest for the middle foci. When tiling FACED FOVs, the dimmer edges led to perception of dark vertical edges. To improve the visibility of structures excited by the early and late foci at the two edges of the FACED line scans, we normalized the pixel values across the FACED line scans as detailed below.

Each image had its rows along the FACED line-scan direction and its columns along the Y galvo scanning direction (for activity imaging data, their time-averaged image was used to determine the illumination profile). Illumination normalization entailed dividing each row of the image by a line-scan illumination profile, which was estimated via an optimization process. We calculated the percentile values for pixels within each column (at 1%, 2%, …, 100%). For each percentile, we then generated a candidate illumination profile using the corresponding pixel values. We then individually scaled each candidate profile by dividing it by its mean and set all values less than 0.1 in the profile to be 0.1 (the profile thus had values greater or equal to 0.1). We then used each scaled candidate profile to normalize the raw image and evaluated the strength of vertical edges in the normalized image using a row-wise sobel filter. The illumination profile that minimized the appearance of vertical edges in the normalized image was chosen as the optimal profile that was used for normalization of all images for this FOV.

To generate activity videos, an offset of 0.9 was added to the optimal illumination profile to prevent noise amplification caused by small values within the profile. After illumination normalization, we then plotted in the activity videos ΔF for each pixel, with ΔF = F(*t*) − min(F(*t*)), which improved the visibility of active neurons.

## Code availability

Data acquisition and analysis code can be found at https://github.com/JiLabUCBerkeley/FACED2PFM2p0-analysis.

