## Supplementary information for "FACED 2.0 enables large-scale voltage and calcium imaging *in vivo*"

### This PDF file includes the following:

### Extended Data Figures

Extended Data Figure 1. Optical layout of the FACED 2.0 system.

Extended Data Figure 2. PSF measurements across the entire imaging FOV of a 25× 1.05 NA objective.

Extended Data Figure 3. PSF measurements across the entire imaging FOV of a 16× 0.8 NA objective.

Extended Data Figure 4. Subcellular resolution of FACED 2.0 with a 16× 0.8 NA objective.

Extended Data Figure 5. FACED 2.0 with a 25× 1.05 NA objective visualizes neuronal processes down to 800 µm in vivo.

Extended Data Figure 6. FACED 2.0 with a 16× 0.8 NA objective visualizes neuronal processes down to 750 µm in vivo.

Extended Data Figure 7. FACED 2.0 with a 16× 0.8 NA objective visualizes capillaries down to 750 µm in vivo.

Extended Data Figure 8. Representative -ΔF/F voltage traces (in blue) of all 182 neurons in Fig. 2.

Extended Data Figure 9. Photobleaching of JEDI-2P-kv fluorescence during FACED 2PFM imaging.

Extended Data Figure 10. Raster plots of (top) spikes and (bottom) subthreshold -ΔF/F traces from 27 8.25-s trials of 182 neurons in Fig. 2.

Extended Data Figure 11. Imaging voltage activity from population of neurons in vivo at 2.6 ms/frame.

Extended Data Figure 12. Representative -ΔF/F voltage traces (in blue) of all 225 neurons in Extended Data Fig. 11.

Extended Data Figure 13. Raster plots of (top) spikes and (bottom) subthreshold -ΔF/F traces from 26 8.32-s trials of 225 neurons in Extended Data Fig. 11.

Extended Figure 14. Analysis comparison of an 800-Hz voltage imaging dataset and its 2× down-sampled (to 400 Hz) data.

Extended Data Figure 15. Simultaneously recorded ΔF/F calcium traces from 12,511 neurons during 36 64-s trials.

Extended Data Figure 16. Large-scale in vivo volumetric imaging of calcium activity through the depth of visual cortex.

Extended Data Figure 17. Simultaneously recorded ΔF/F calcium traces from 14,005 neurons in during 51 trials.

Extended Data Figure 18. 3.8 Hz whole-brain imaging of spontaneous calcium activity in a transgenic zebrafish larva.

Extended Data Figure 19. 10.4 Hz whole-brain imaging of spontaneous calcium activity in a transgenic zebrafish larva.

Extended Data Figure 20. Example 10-trial-averaged ΔF/F calcium traces from calcium imaging datasets.

### Supplementary Notes

Supplementary Note 1: Additional characterization of volumetric imaging via continual Z movement of microscope objective

Supplementary Note 2: Additional analysis on spike detection pipeline

Supplementary Note 3: Voltage imaging at 800 Hz versus 400 Hz

Supplementary Note 4: Discussion on potential contamination due to fluorescence lifetime in activity imaging

Supplementary Note 5: FACED 2.0 versus FACED 1.0

### Supplementary Tables

Supplementary Table 1: Comparison of 2PFM techniques

Supplementary Table 2: Parameters for 2D imaging with FACED 2.0

Supplementary Table 3: Parameters for 3D imaging with FACED 2.0

Supplementary Table 4: Brightness comparison for FACED 2.0 figures

Supplementary Table 5: Estimated numbers of photons in activity measurement datasets

Supplementary Table 6: Estimated baseline noise level for activity measurement datasets

### Other Supplementary Materials for this manuscript include the following:

Supplementary Video 1. Imaging cerebral blood flow over 1,111 µm × 1,000 µm at 5.3 ms/frame in the mouse cortex *in vivo* using the 16× 0.8 NA objective.

Supplementary Video 2. Volumetric calcium imaging over 1,111 µm × 1,000 µm × 400 µm at 259.7 ms/vol in the visual cortex of a Slc17a7-IRES2-Cre × Ai162D mouse using the 16× 0.8 NA objective.

Supplementary Video 3. Volumetric calcium imaging over 1,111 µm × 1,000 µm × 780 µm at 365.7 ms/vol in the visual cortex of a wild-type mouse transfected with GCaMP6s using the 16× 0.8 NA objective.

Supplementary Video 4. Volumetric calcium imaging over 480 µm × 585 µm × 360 µm at 262.4 ms/vol of a larval GCaMP6s transgenic zebrafish (Tg[Elavl3:H2B-GcaMP6s]) brain using the 25× 1.05 NA objective.

Supplementary Video 5. Volumetric calcium imaging over 555.6 µm × 800 µm × 360 µm at 96 ms/vol of a larval GcaMP6s transgenic zebrafish (Tg[Elavl3:H2B-GCaMP6s]) brain using the 16× 0.8 NA objective.

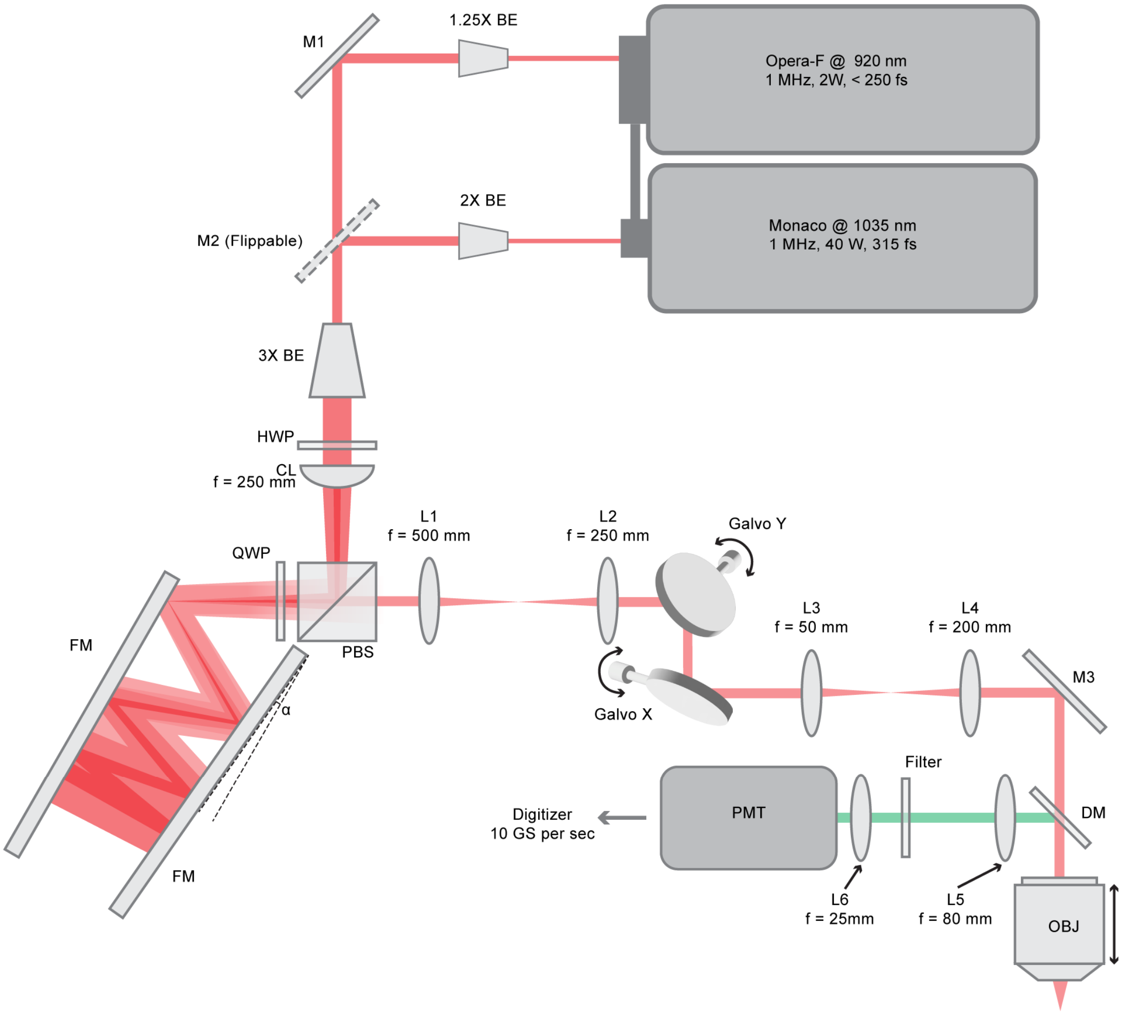

**Extended Data Figure 1. Optical layout of the FACED 2.0 system**. BE: beam expander; M: mirror; HWP: half-wave plate; CL: cylindrical lens; PBS: polarizing beam splitter; QWP: quarter-wave plate; FM: FACED mirror; α: angle between the FACED mirrors; L: lens; Galvo: galvanometer scanner; DM: dichroic mirror; OBJ: objective lens, PMT: photomultiplier tube.

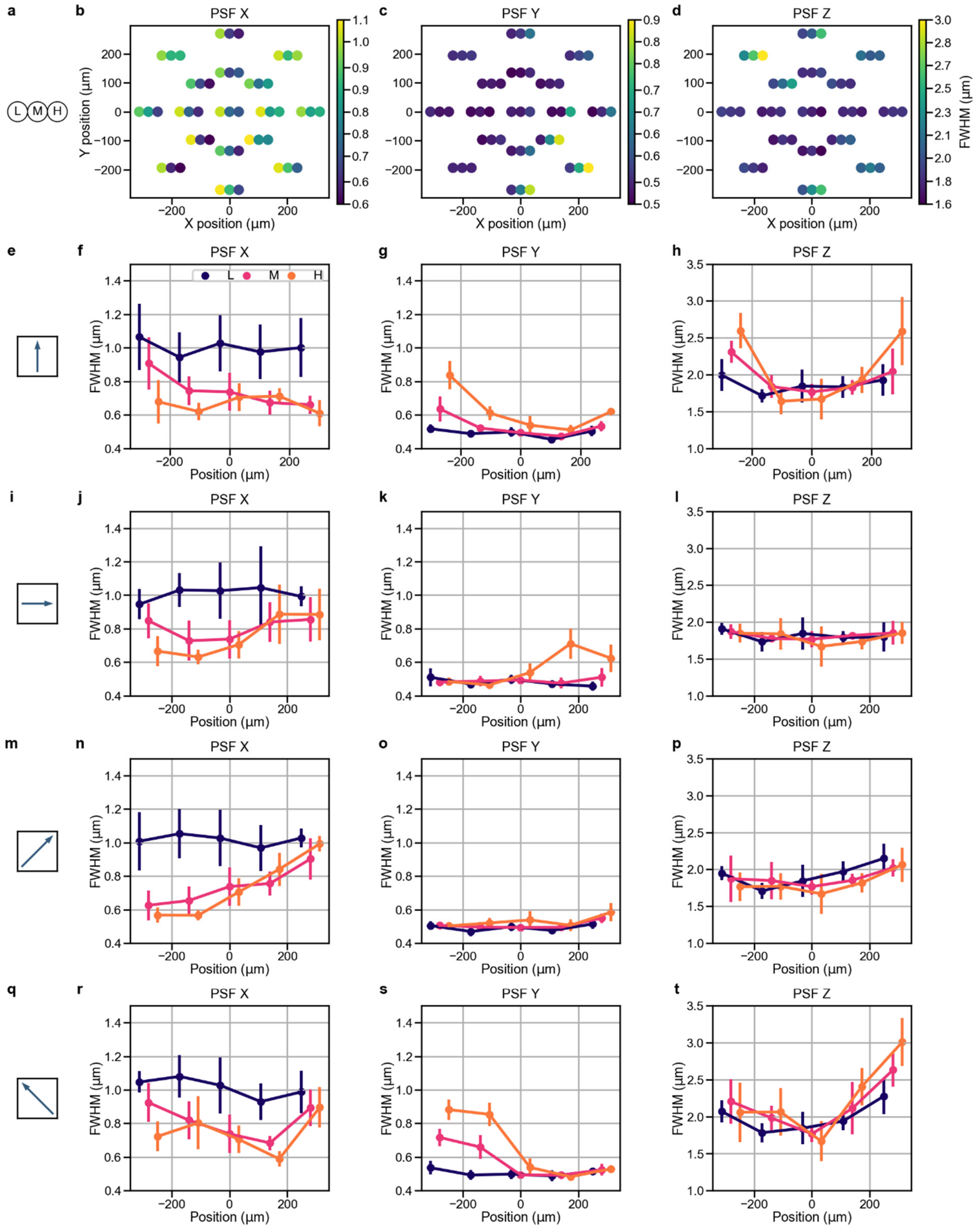

**Extended Data Figure 2. PSF measurements across the entire imaging FOV of a 25× 1.05 NA objective.** (**a**) Three consecutive dots with labels "L", "M", and "H" denote the average FWHMs of PSFs measured for the first (L), middle (M), and last (H) third of foci within a FACED line. (**b, c, d**) FWHMs along the X (FACED line) (**b**), Y (**c**), and Z (**d**) directions, respectively, measured at different FOV locations, using notation in **a**. (**e-h**) Variation of L, M, and H FWHMs (**e**) along the middle vertical axis of the FOV for (**f**) X, (**g**) Y, and (**h**) Z PSFs. (**i-l**) Variation of L, M, and H FWHMs (**i**) along the middle horizontal axis of the FOV for (**j**) X, (**k**) Y, and (**l**) Z PSFs. (**m-p**) Variation of L, M, and H FWHMs (**m**) along one diagonal axis of the FOV for (**n**) X, (**o**) Y, and (**p**) Z PSFs. (**q-t**) Variation of L, M, and H FWHMs (**q**) along the other diagonal axis of the FOV for (**r**) X, (**s**) Y, and (**t**) Z PSFs. Error bar: SD. PSFs were measured from 0.2-µm-diameter fluorescent beads. N > 6 beads were measured for each third of a FACED line. N > 500 beads were measured in total.

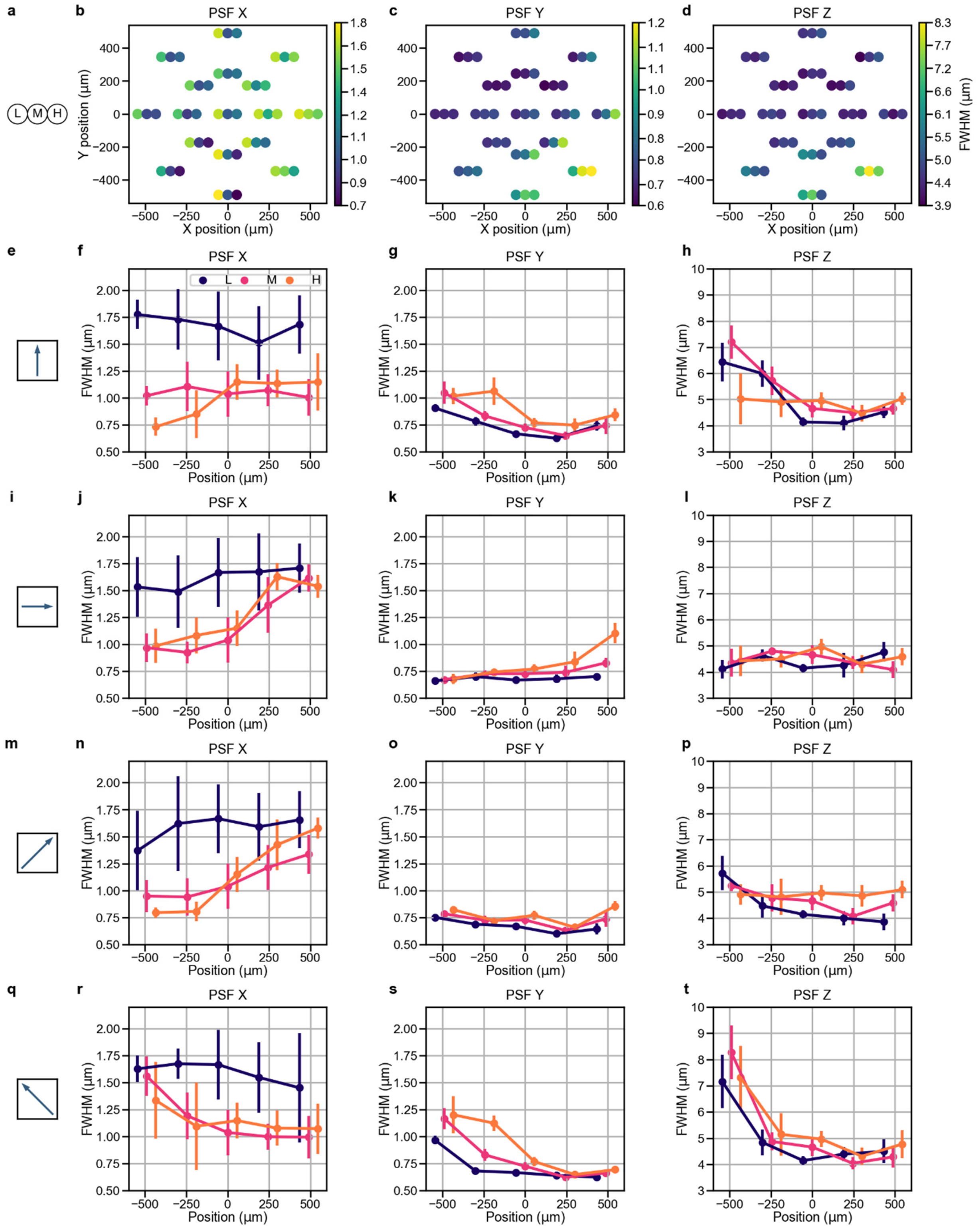

**Extended Data Figure 3. PSF measurements across the entire imaging FOV of a 16× 0.8 NA objective.** (**a**) Three consecutive dots with labels "L", "M", and "H" denote the average FWHMs of PSFs measured for the first (L), middle (M), and last (H) third of foci within a FACED line. (**b, c, d**) FWHMs along the X (FACED line) (**b**), Y (**c**), and Z (**d**) directions, respectively, measured at different FOV locations, using notation in **a**. (**e-h**) Variation of L, M, and H FWHMs (**e**) along the middle vertical axis of the FOV for (**f**) X, (**g**) Y, and (**h**) Z PSFs. (**i-l**) Variation of L, M, and H FWHMs (**i**) along the middle horizontal axis of the FOV for (**j**) X, (**k**) Y, and (**l**) Z PSFs. (**m-p**) Variation of L, M, and H FWHMs (**m**) along one diagonal axis of the FOV for (**n**) X, (**o**) Y, and (**p**) Z PSFs. (**q-t**) Variation of L, M, and H FWHMs (**q**) along the other diagonal axis of the FOV for (**r**) X, (**s**) Y, and (**t**) Z PSFs. Error bar: SD. PSFs were measured from 0.2-μm-diameter fluorescent beads. N > 6 beads were measured for each third of a FACED line. N > 500 beads were measured in total.

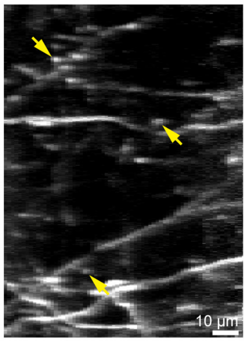

**Extended Data Figure 4. Subcellular resolution of FACED 2.0 with a 16× 0.8 NA objective.** Maximum intensity projection of a 92 µm × 128 µm × 10 µm volume of dendrites at 29.5 µm depth in a Thy1-GFP line M mouse cortex imaged with a 16× 0.8 NA objective. Arrows: putative synapses. 1035 nm excitation at 165 mW post objective.

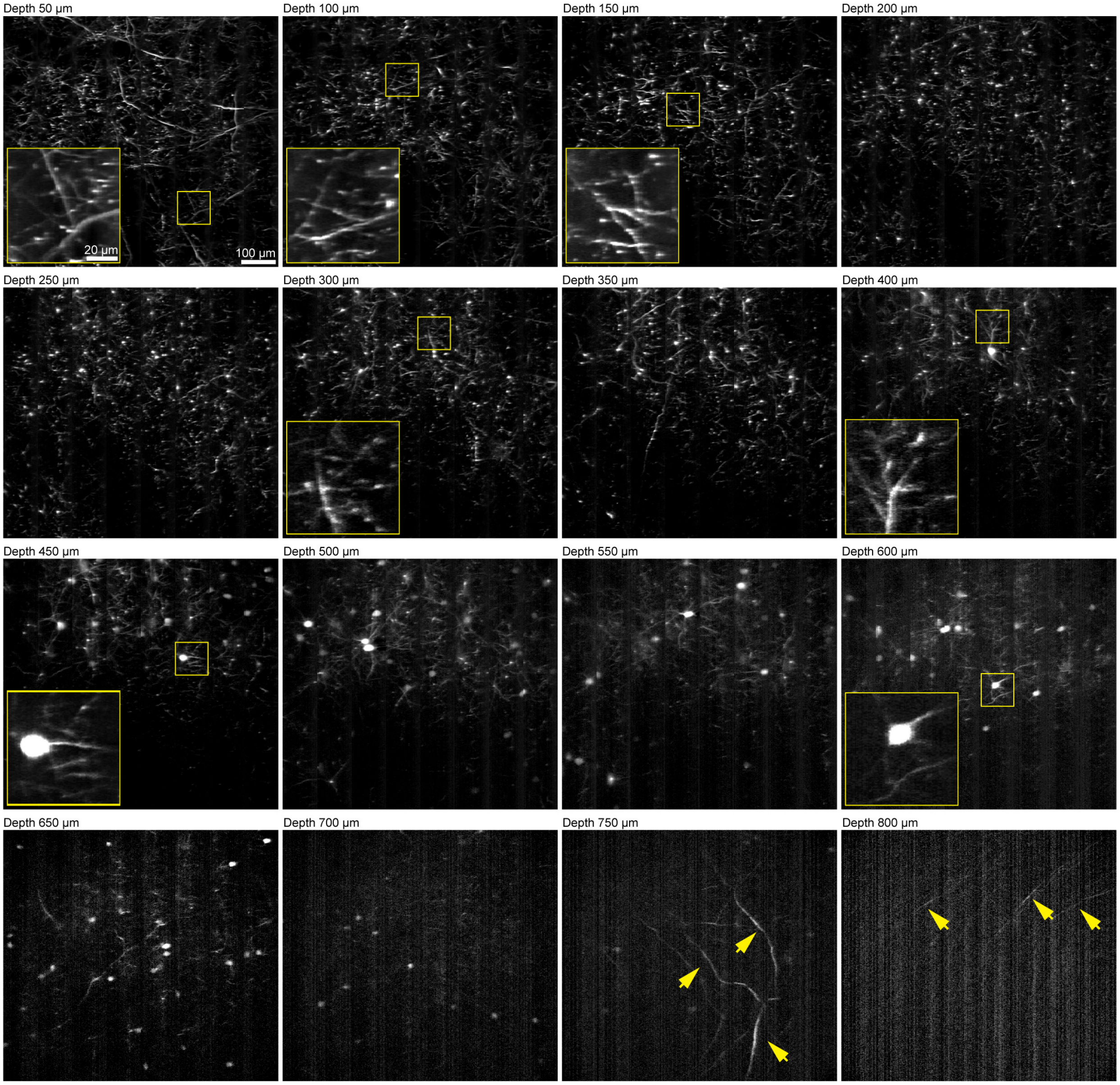

**Extended Data Figure 5. FACED 2.0 with a 25× 1.05 NA objective visualizes neuronal processes down to 800 µm *in vivo*.** 640 µm × 585 µm images acquired from different depths in a Thy1-GFP line M mouse cortex. Pixel size: 0.8 µm × 0.65 µm. Insets: zoomed-in views of areas in yellow boxes. Arrows: neuronal processes at 750 µm and 800 µm depths. 1035 nm excitation at 140 mW post objective.

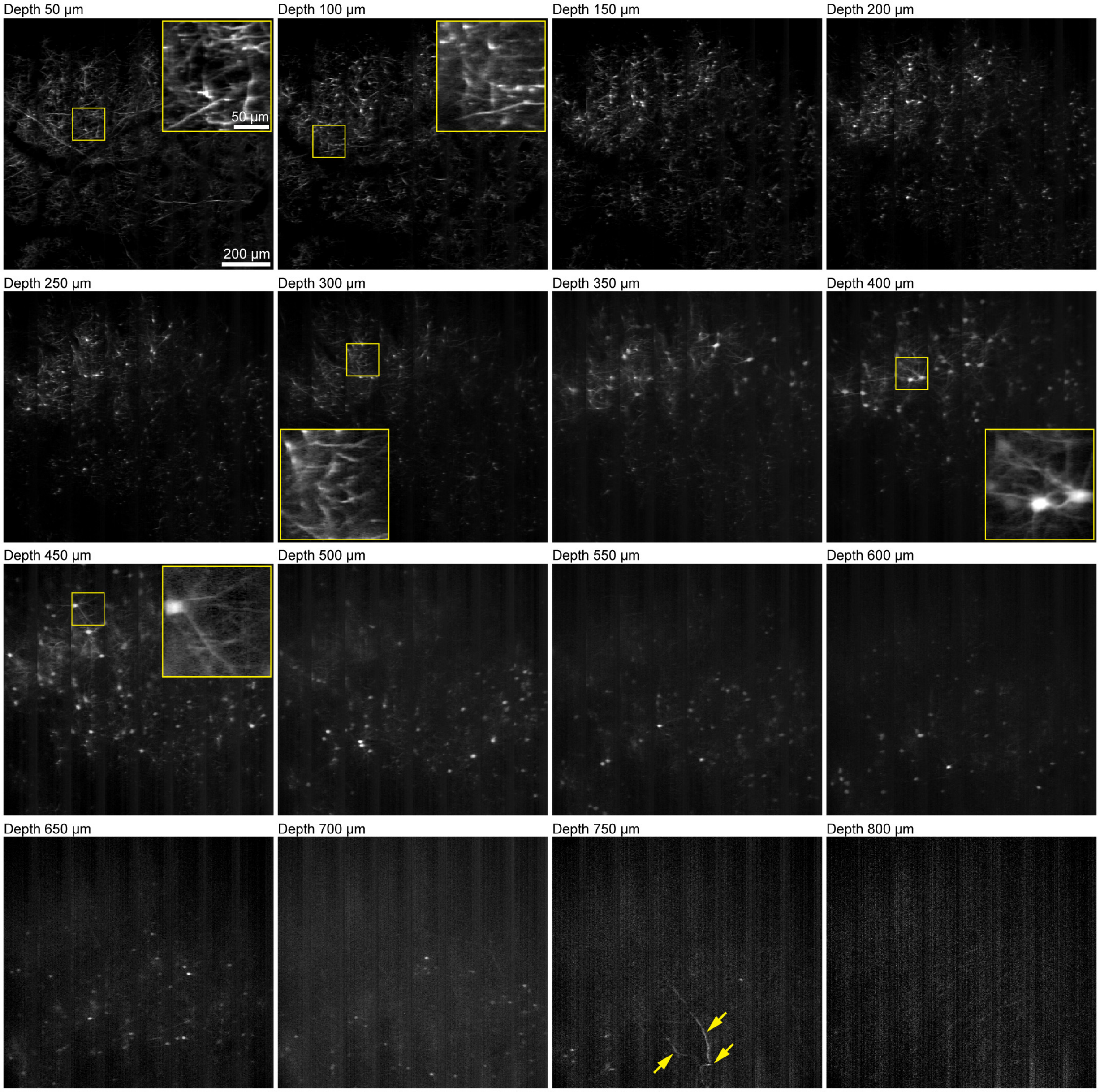

**Extended Data Figure 6. FACED 2.0 with a 16× 0.8 NA objective visualizes neuronal processes down to 750 μm *in vivo*.** 1,111 μm × 1,035 μm images acquired from different depths in a Thy1-GFP line M mouse cortex. Pixel size: 1.4 μm × 1.15 μm. Insets: zoomed-in views of areas in yellow boxes. Arrows: neuronal processes at 750 μm and 800 μm depths. 1035 nm excitation at 170 mW post objective.

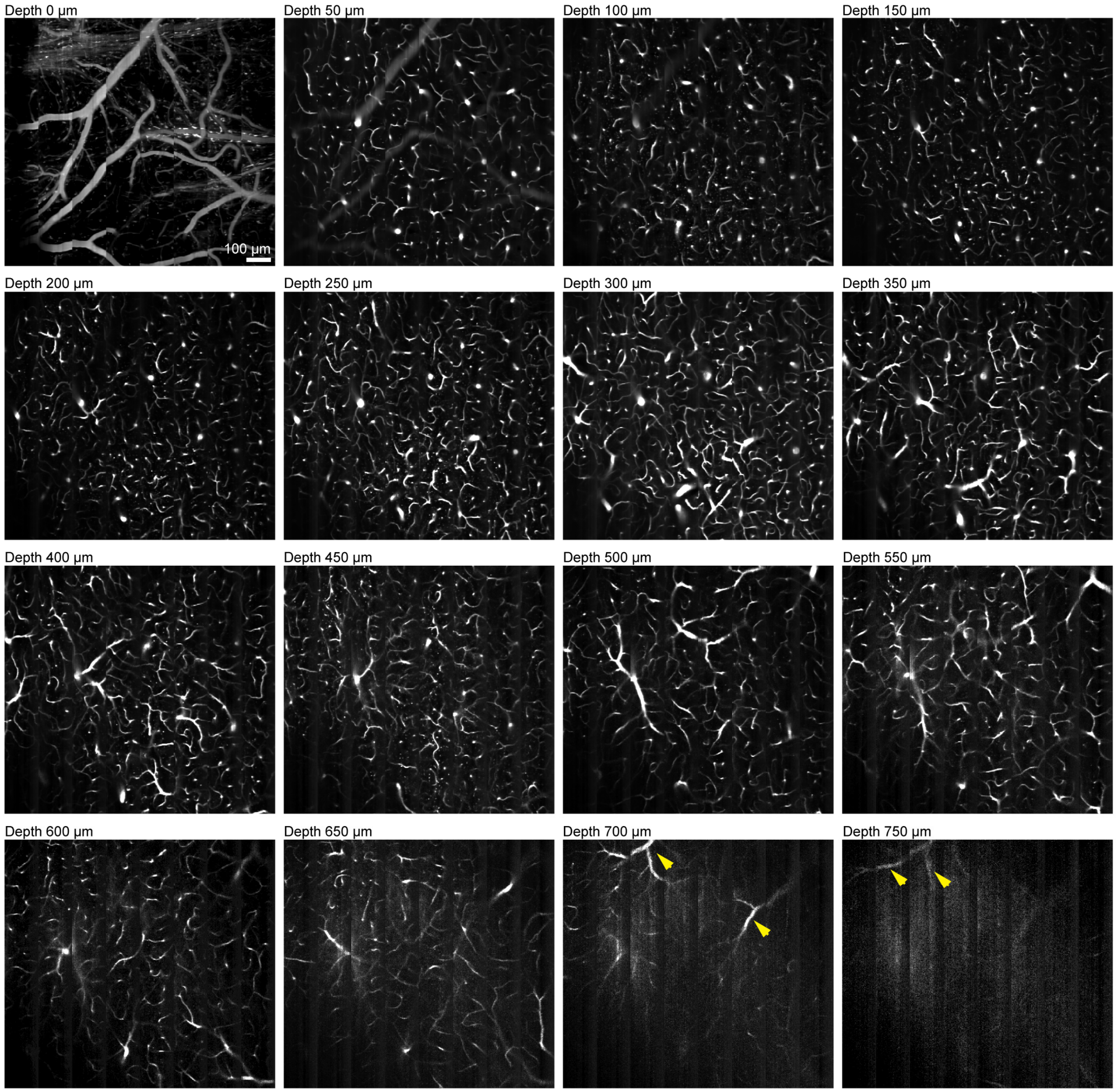

**Extended Data Figure 7. FACED 2.0 with a 16× 0.8 NA objective visualizes capillaries down to 750 μm *in vivo*.** 1,111 μm × 1,000 μm images acquired from different depths in a mouse cortex. Capillaries are labeled with dextran-conjugated Rhodamine B. Pixel size: 1.4 μm × 2 μm. Arrows: capillaries at 700 μm and 750 μm depths. 1035 nm excitation at 208 mW post objective.

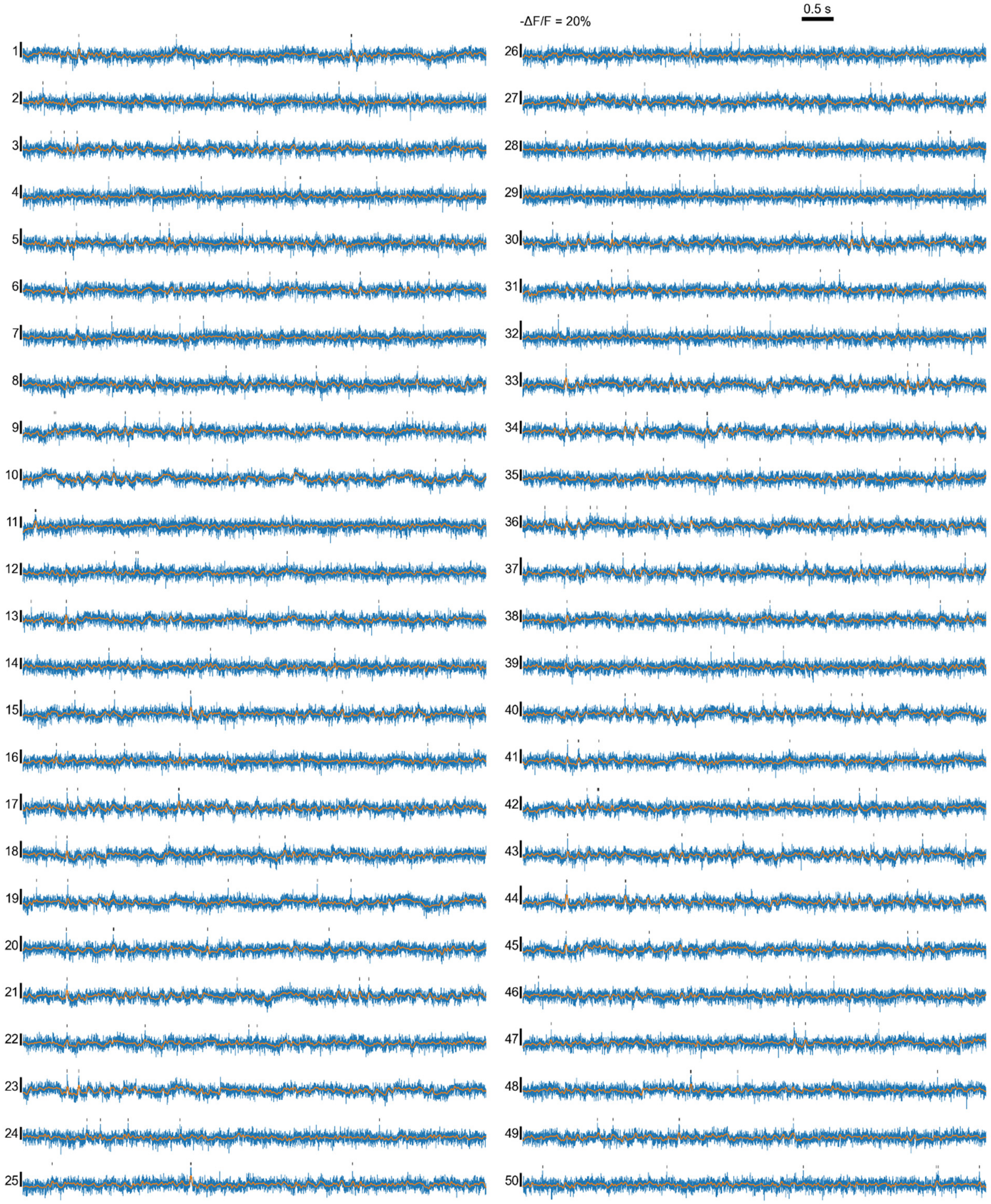

0.5 s
-ΔF/F = 20%

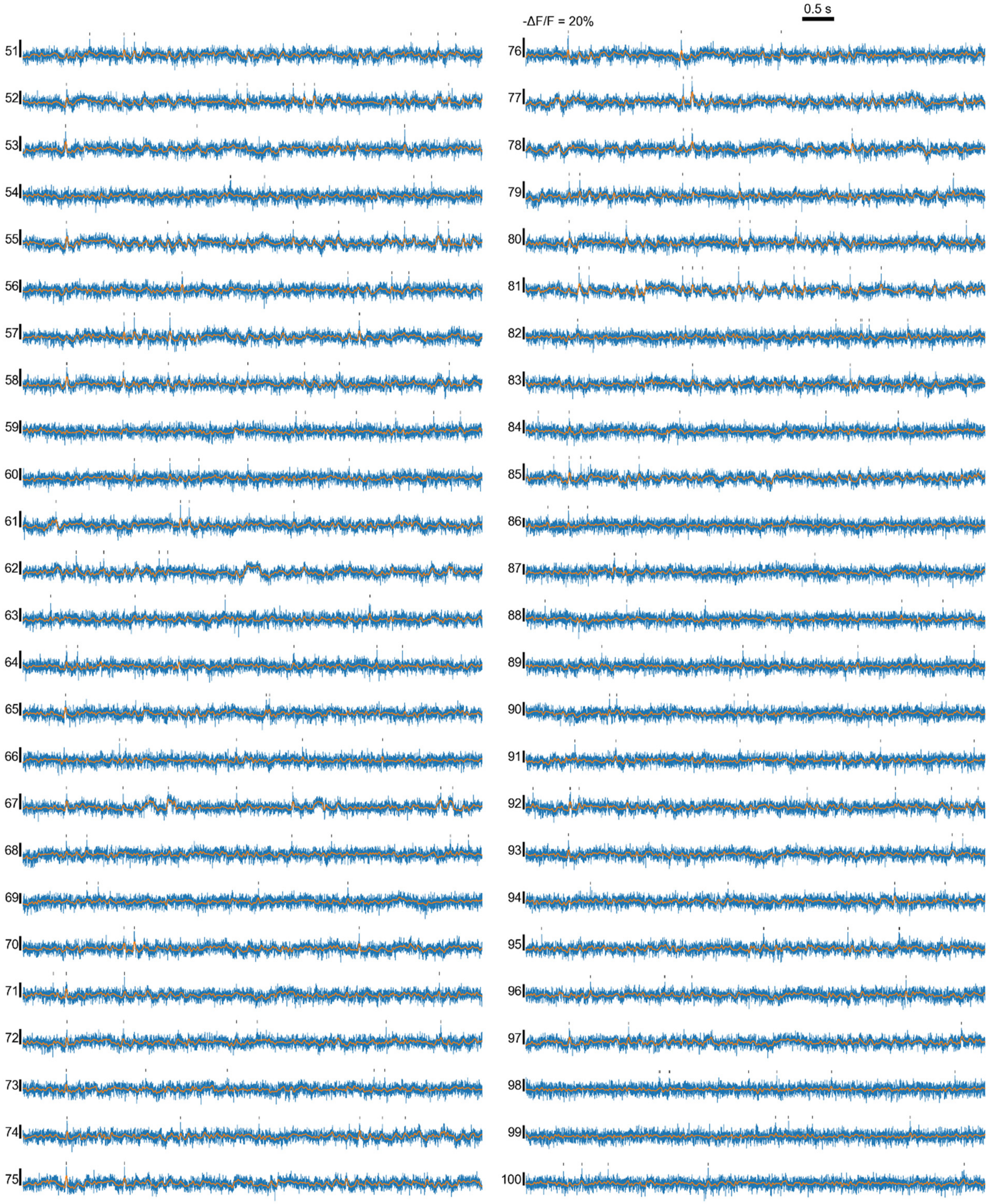

0.5 s
-ΔF/F = 20%

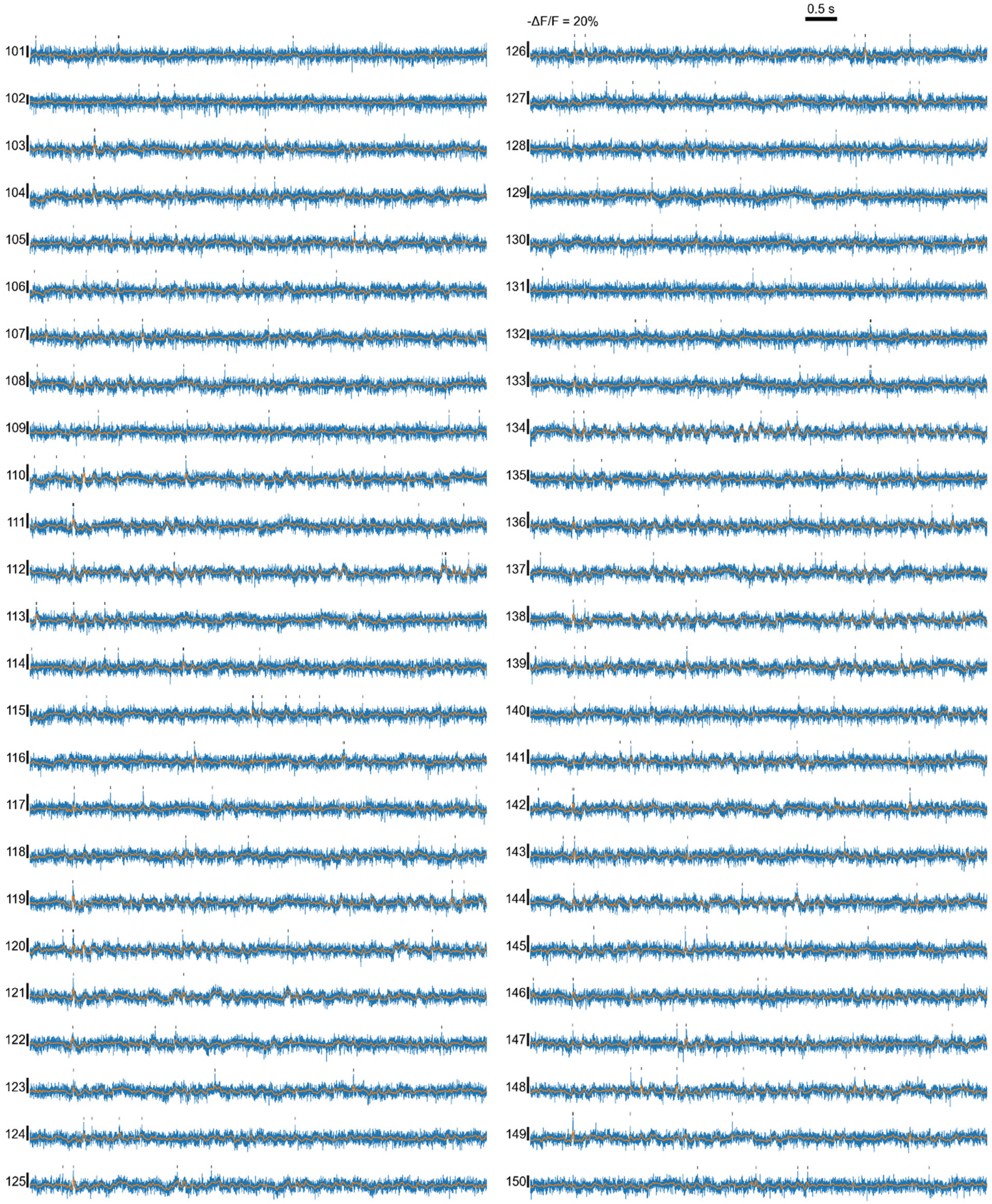

0.5 s
-ΔF/F = 20%

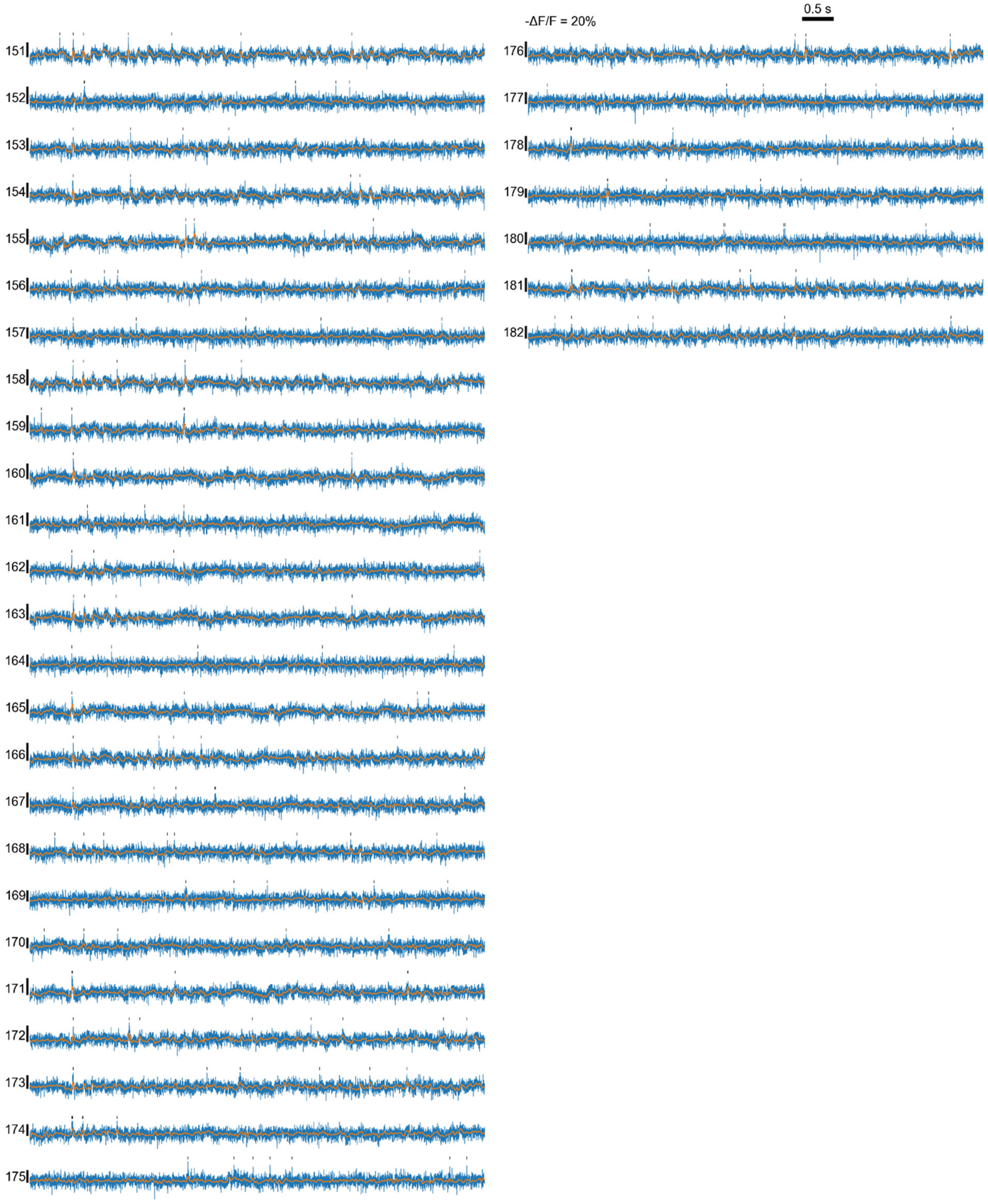

**Extended Data Figure 8. Representative -ΔF/F voltage traces (in blue) of all 182 neurons in Fig. 2.** Black ticks: suprathreshold spikes. Orange traces: extracted subthreshold -ΔF/F.

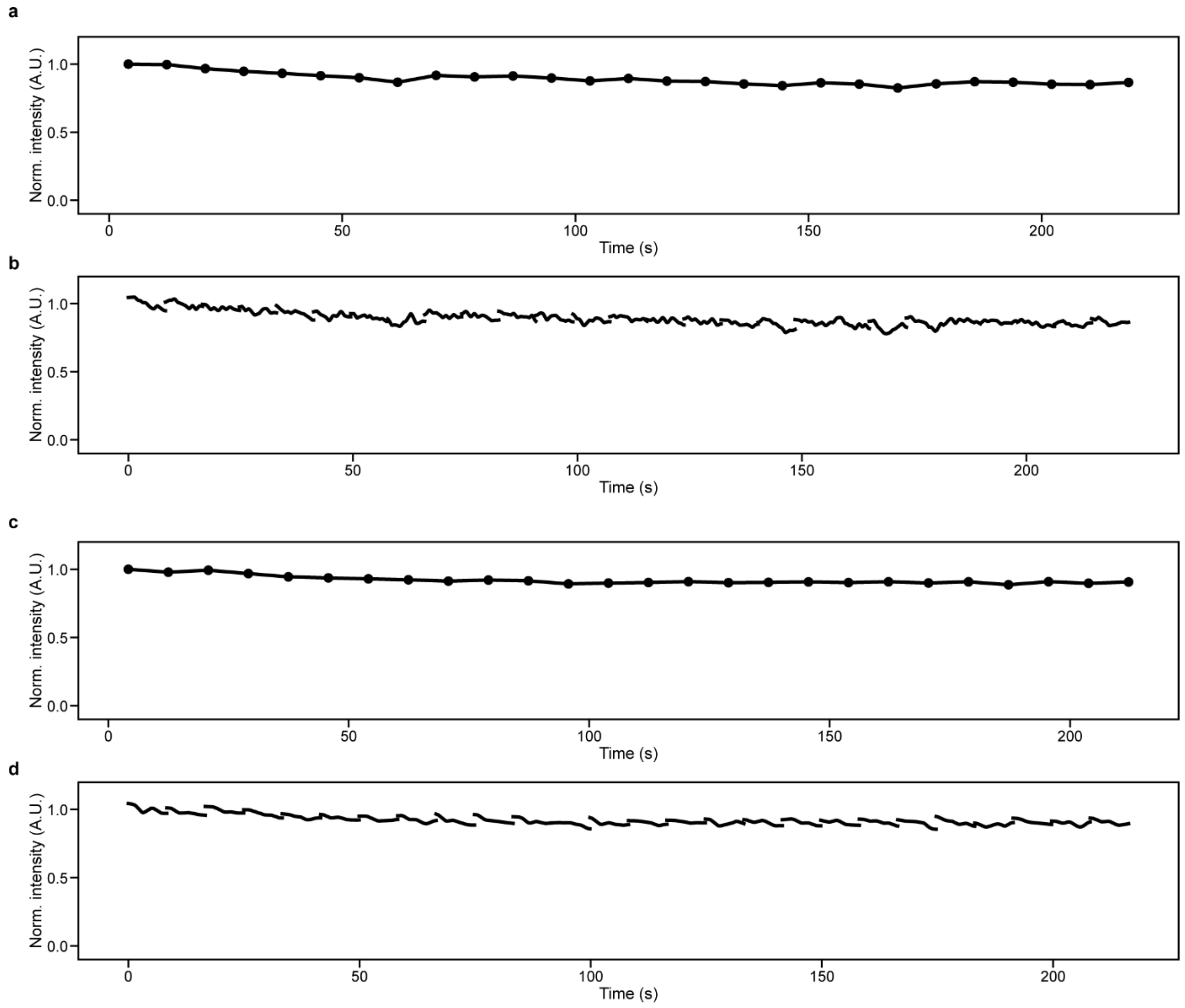

**Extended Data Figure 9. Photobleaching of JEDI-2P-kv fluorescence during FACED 2PFM imaging.** (**a**) Time-averaged fluorescence signal over each 8.25-s continuous imaging trial across 27 trials. Same data as **Fig. 2**, 160 µm × 400 µm FOV acquired at 1.25 ms/frame for a total duration of 222.75 s using 1035 nm excitation at 163 mW post-objective from a depth of 132 µm below dura. A 15% photobleaching observed over the entire 222.75-s acquisition duration. (**b**) Fluorescence signal over each 8.25-s continuous imaging trial across 27 trials. Same data as **a** and an average of 4% photobleaching observed within each 8.25-s-long trial. (**c**) Time-averaged fluorescence signal over each 8.32-s continuous imaging trial across 26 trials. Same data as **Extended Data Fig. 11**, 320 µm × 400 µm FOV acquired at 2.6 ms/frame for a total duration of 216.32 s using 1035 nm excitation at 164 mW post-objective from a depth of 140 µm below dura. A 10% photobleaching observed over the entire 216.32-s acquisition duration. (**d**) Fluorescence signal over each 8.32-s continuous imaging trial across 26 trials. Same data as **c** and an average of 5% photobleaching observed within an 8.32-s-long trial. Pixel size for all data: 0.8 µm × 0.8 µm.,

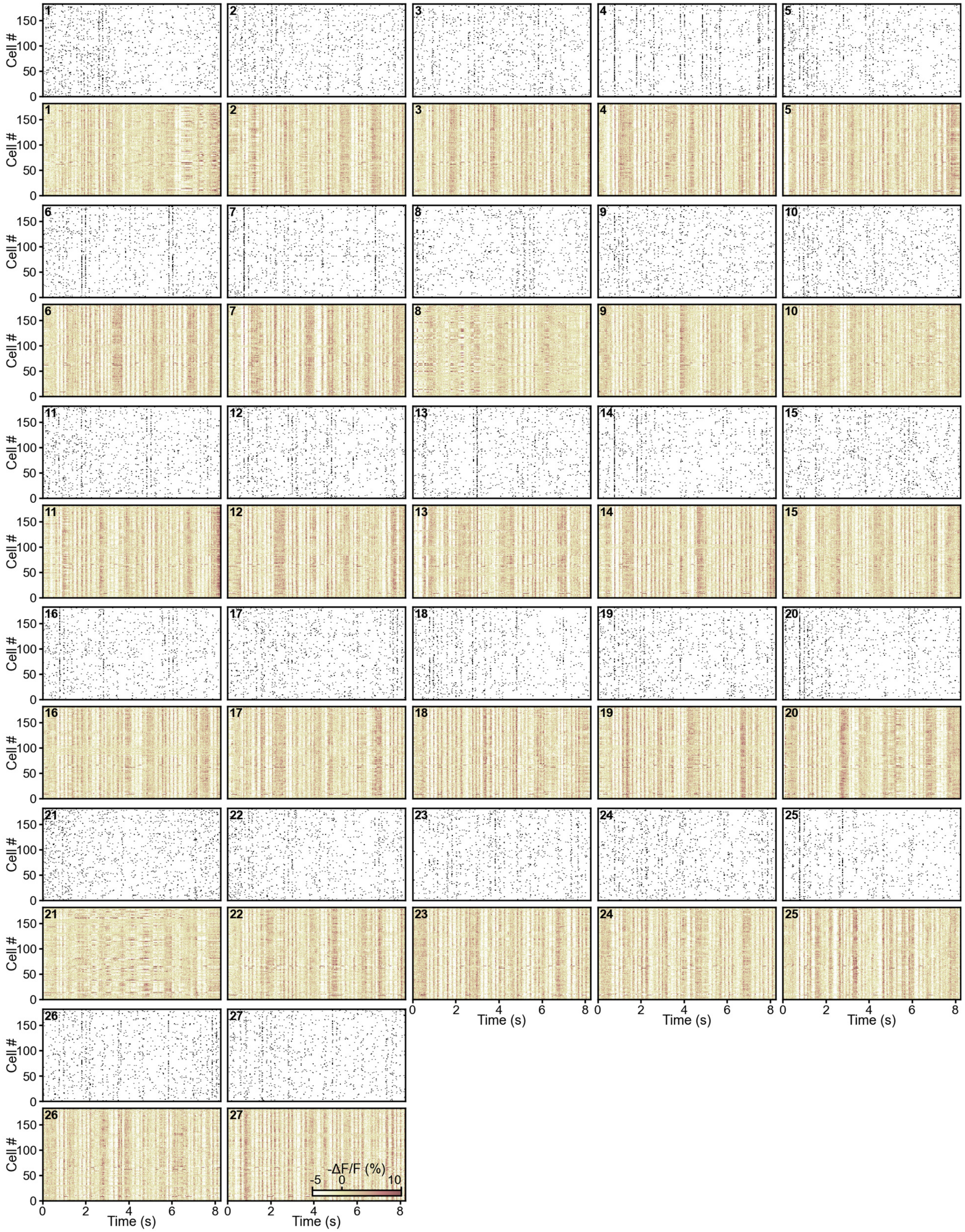

**Extended Data Figure 10. Raster plots of (top) spikes and (bottom) subthreshold -ΔF/F traces from 27 8.25-s trials of 182 neurons in Fig. 2.**

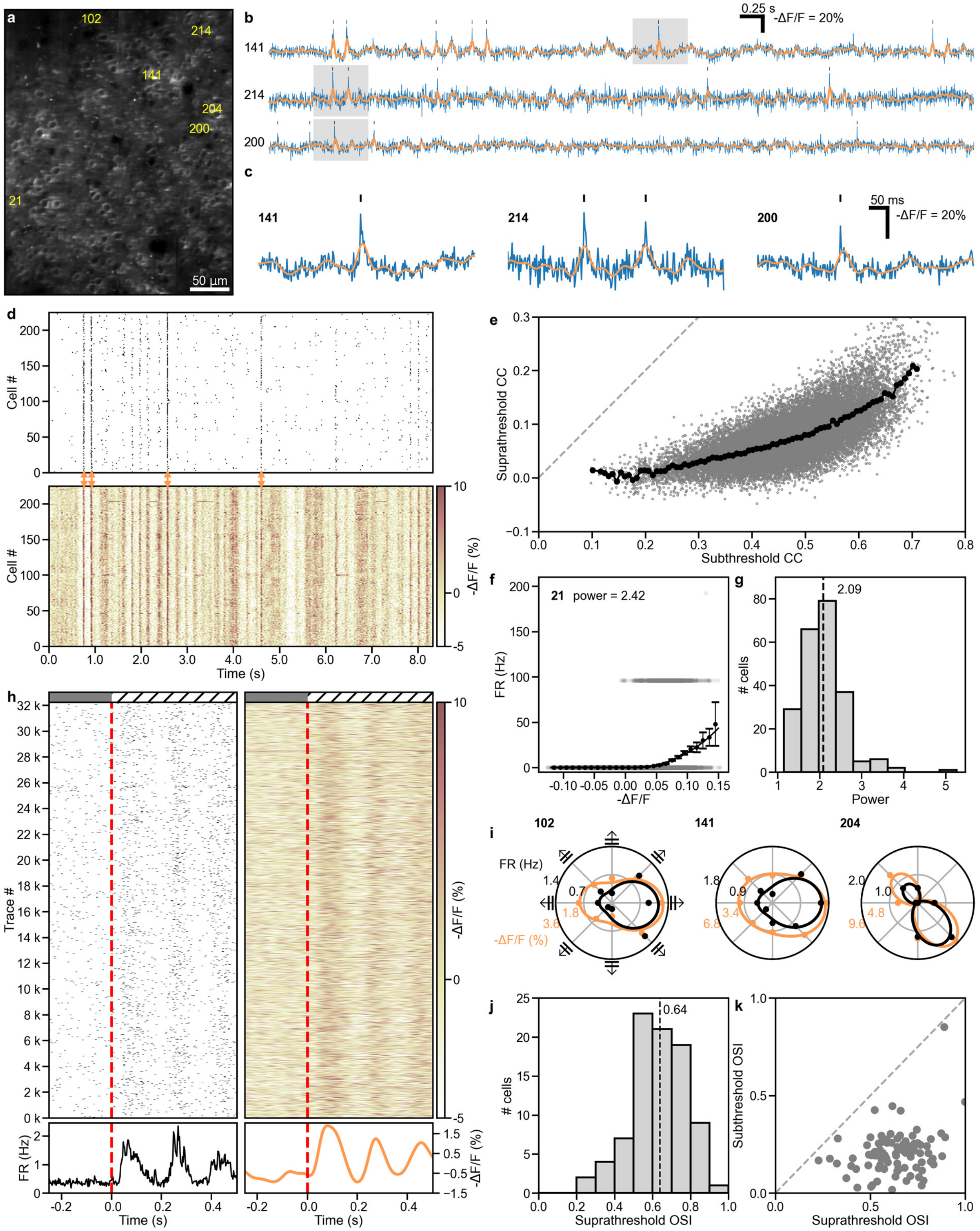

**Extended Data Figure 11. Imaging voltage activity from population of neurons in vivo at 2.6 ms/frame.** (**a**) JEDI-2P-kv-expressing L2/3 neurons in a 320 µm × 400 µm FOV within a mouse visual cortex imaged at 140 µm depth and 2.6 ms/frame. Pixel size: 0.8 µm × 0.8 µm. 1035 nm excitation at 164 mW post 1.05-NA objective. Numbers: representative neurons. (**b**) Representative -ΔF/F voltage traces (in blue) of numbered neurons in **a**. Black ticks: suprathreshold spikes. Orange traces: extracted subthreshold -ΔF/F. (**c**) Zoom-in view of voltage traces from the gray boxes in **b**. (**d**) Raster plots of (top) spikes and (bottom) subthreshold -ΔF/F traces of 225 neurons during a representative 8.32-s trial. (**e**) Suprathreshold versus subthreshold pairwise correlation coefficients (CC, gray dots) for 25,200 cell pairs from the 225 neurons in **a**. Black dots: binned averages with 0.01 bin size for subthreshold CC. Dashed gray line: reference line with slope = 1. (**f**) Plot of firing rate (FR) vs subthreshold -ΔF/F for Neuron 21, fit with a power-law function (black curve; power-law exponent = 2.42). Gray dots: FR vs subthreshold -ΔF/F. Black dots: binned average FR with 0.01 bin size in -ΔF/F. Error bar: s.e.m. (**g**) Histogram of fitted power-law exponent values of 225 neurons. Dashed line: median. (**h**) Suprathreshold (left) and subthreshold (right) responses relative to stimulus onset extracted from 32,240 voltage traces of 155 neurons exhibiting visually evoked activity across 26 trials × 8 drifting grating orientations. Top to bottom: raster plot, averaged firing rate (black curve, bottom left panel) and averaged subthreshold -ΔF/F (orange curve, bottom right panel). Red dashed line: onset of drifting grating stimuli. (**i**) Representative polar-plot tuning curves for supra- (black) and subthreshold (orange) voltage responses of Neuron 102, 141, 204, respectively. Dots: measured responses; curves: double-Gaussian fit. (**j**) Histogram of suprathreshold orientation-selectivity index (OSI) of 86 orientation selective (OS) neurons. Dashed line: median. (**k**) Scatter plot of subthreshold OSI vs. suprathreshold OSI for 86 OS neurons. Dashed line: reference line with slope = 1.

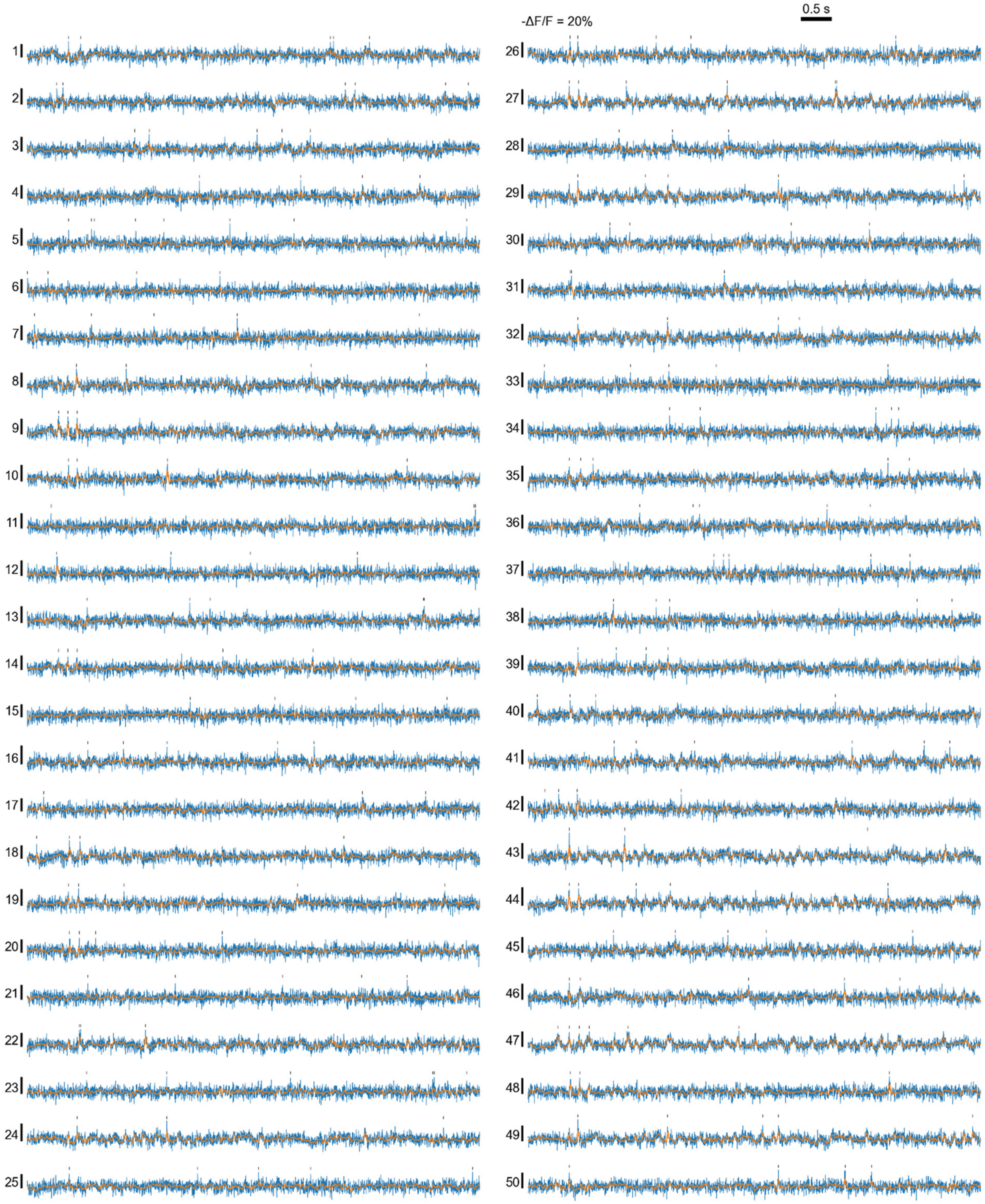

0.5 s
-ΔF/F = 20%

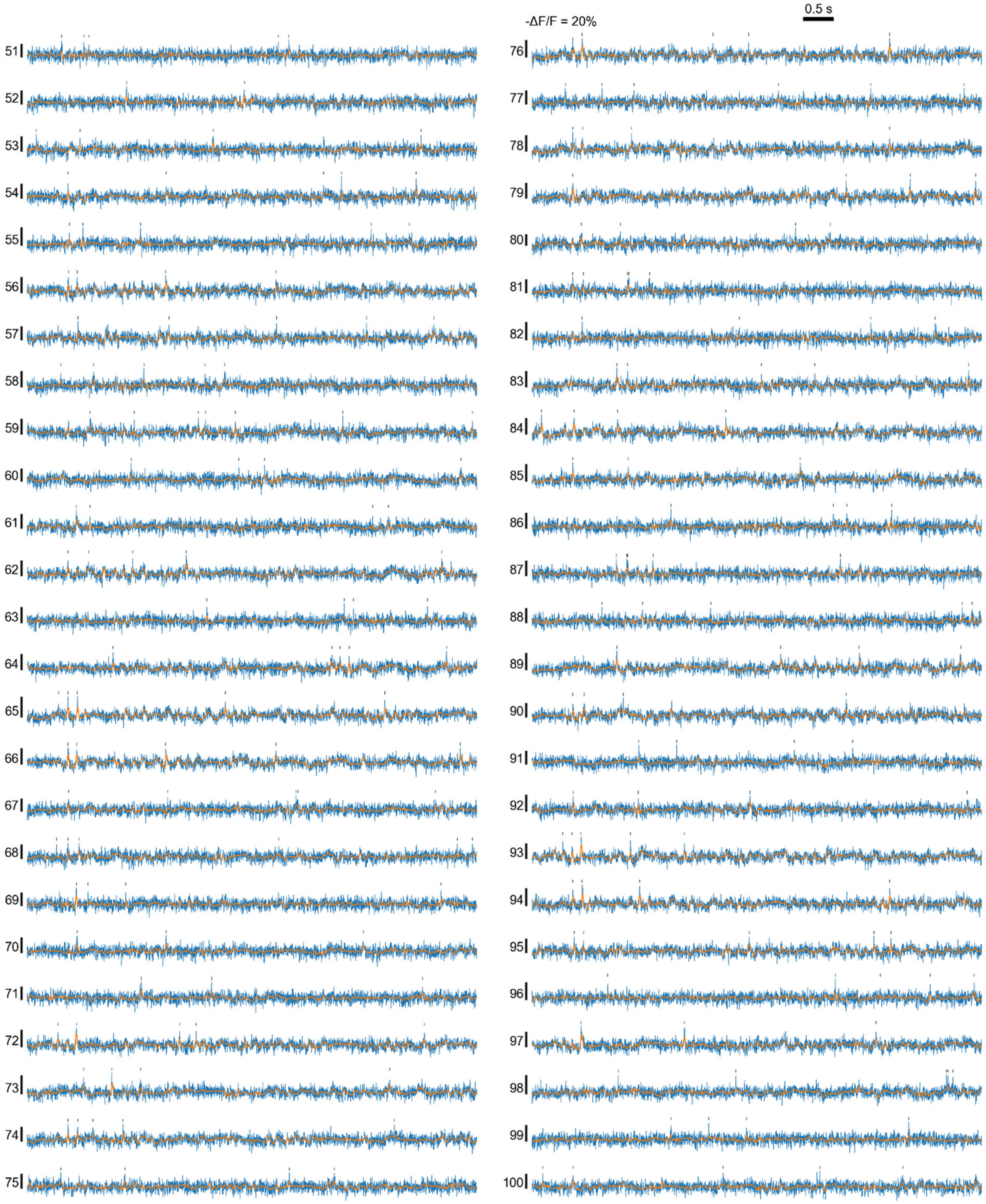

-ΔF/F = 20%
0.5 s

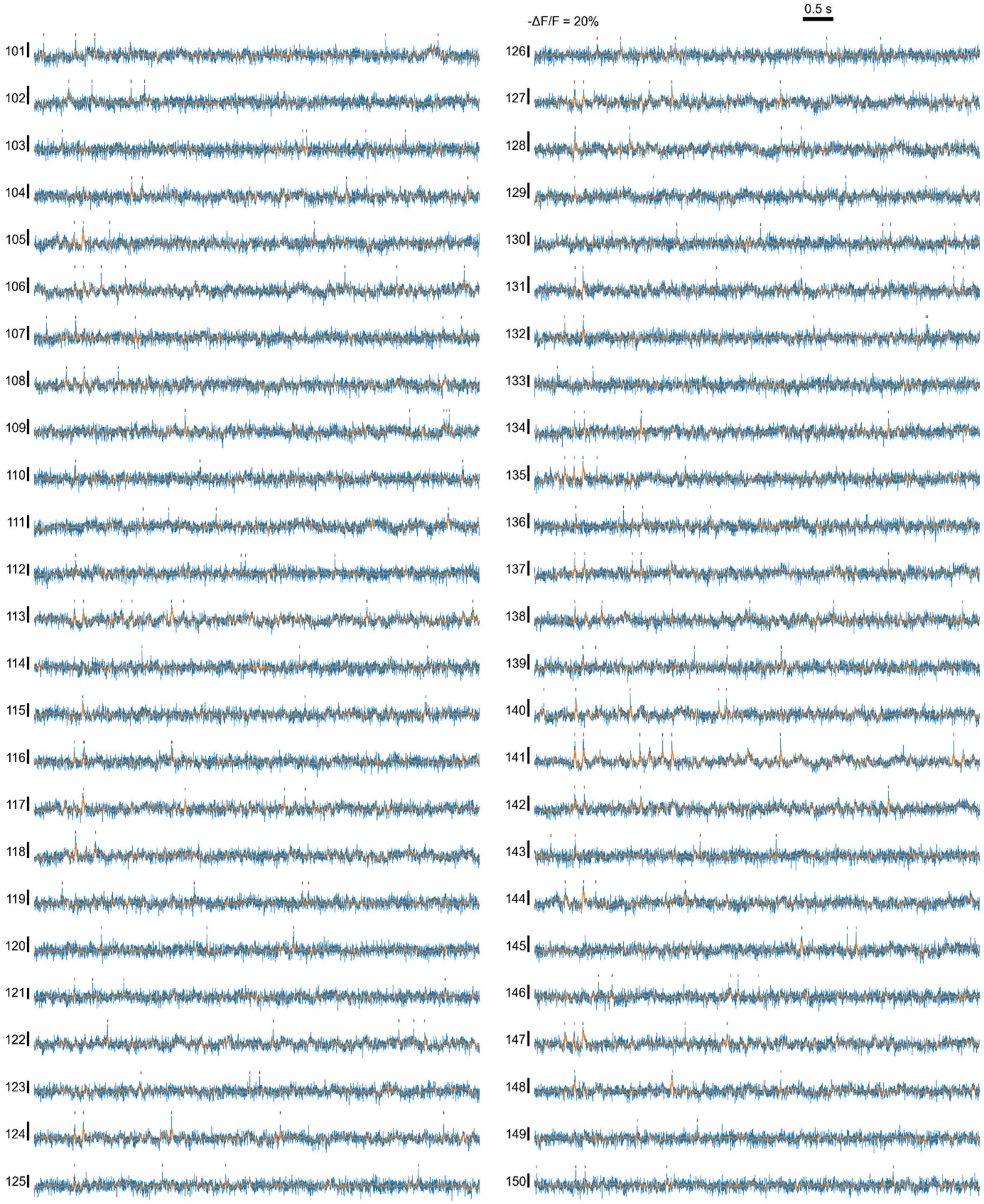

0.5 s
-ΔF/F = 20%

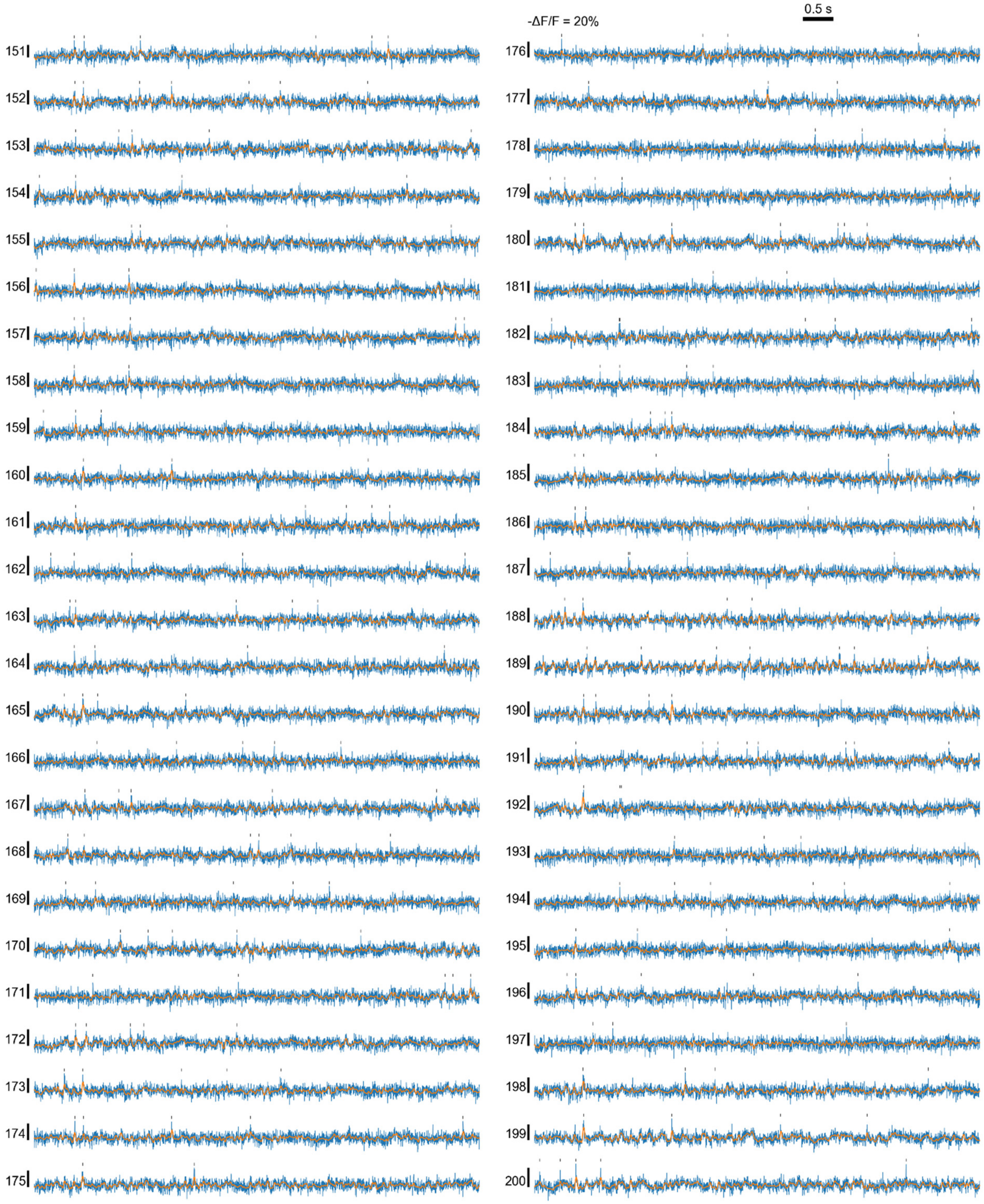
0.5 s
-ΔF/F = 20%

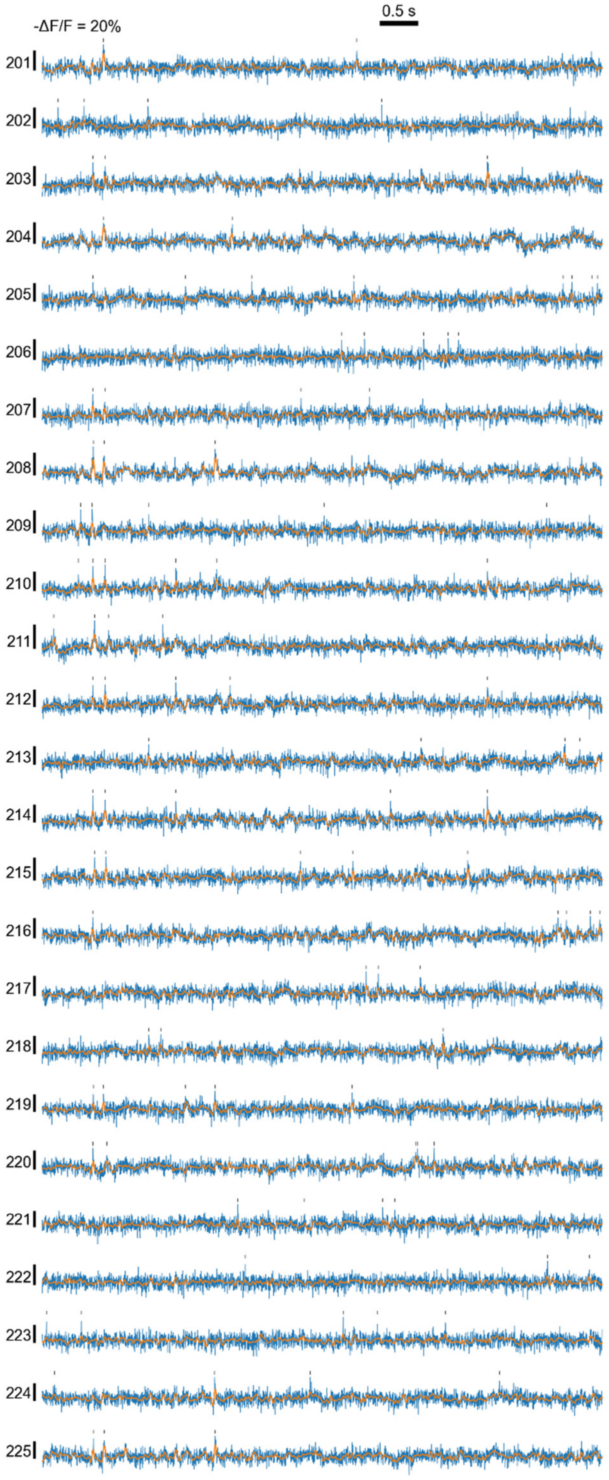

**Extended Data Figure 12. Representative -ΔF/F voltage traces (in blue) of all 225 neurons in Extended Data Fig. 11.** Black ticks: suprathreshold spikes. Orange traces: extracted subthreshold -ΔF/F.

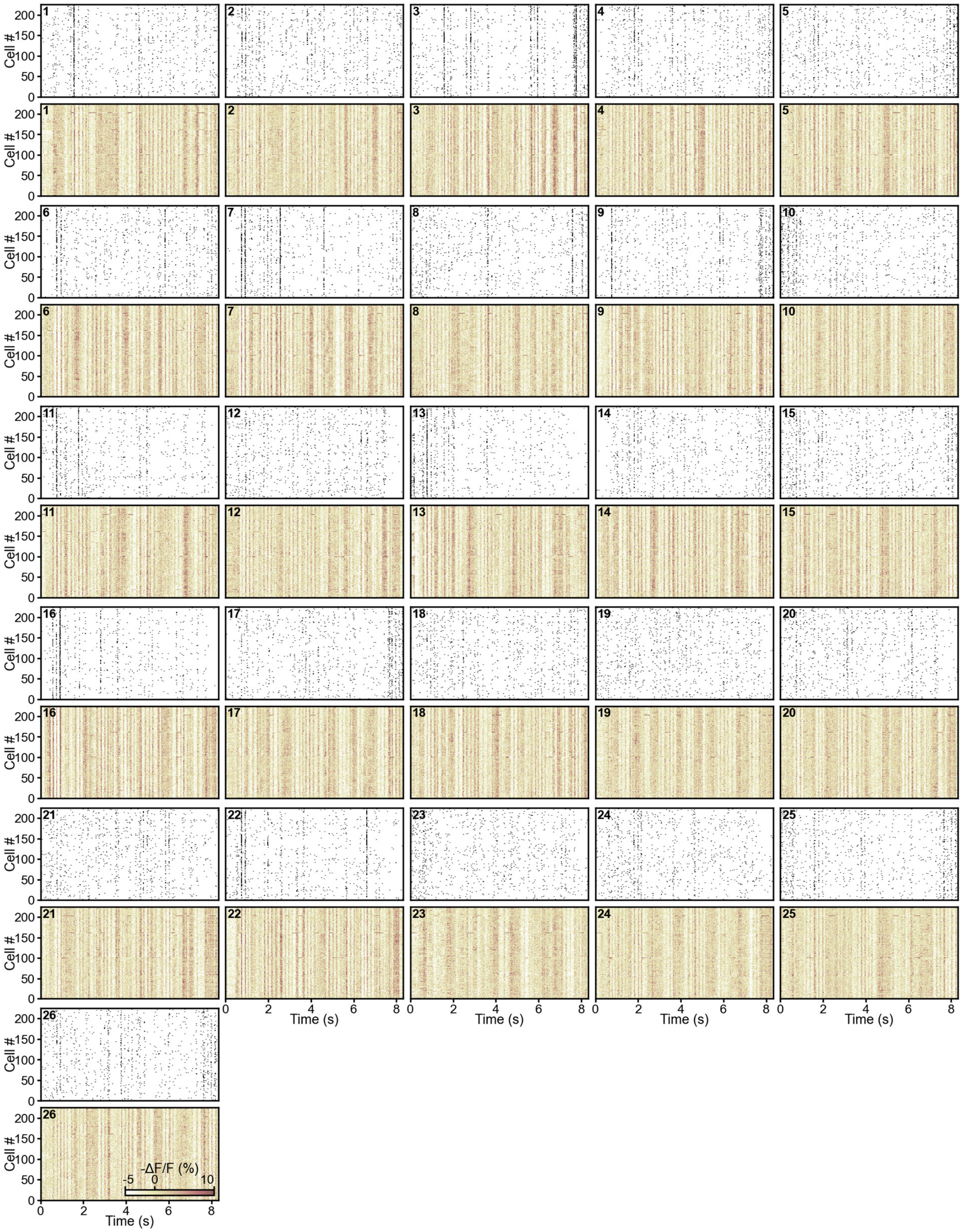

**Extended Data Figure 13. Raster plots of (top) spikes and (bottom) subthreshold -ΔF/F traces from 26 8.32-s trials of 225 neurons in Extended Data Fig. 11.**

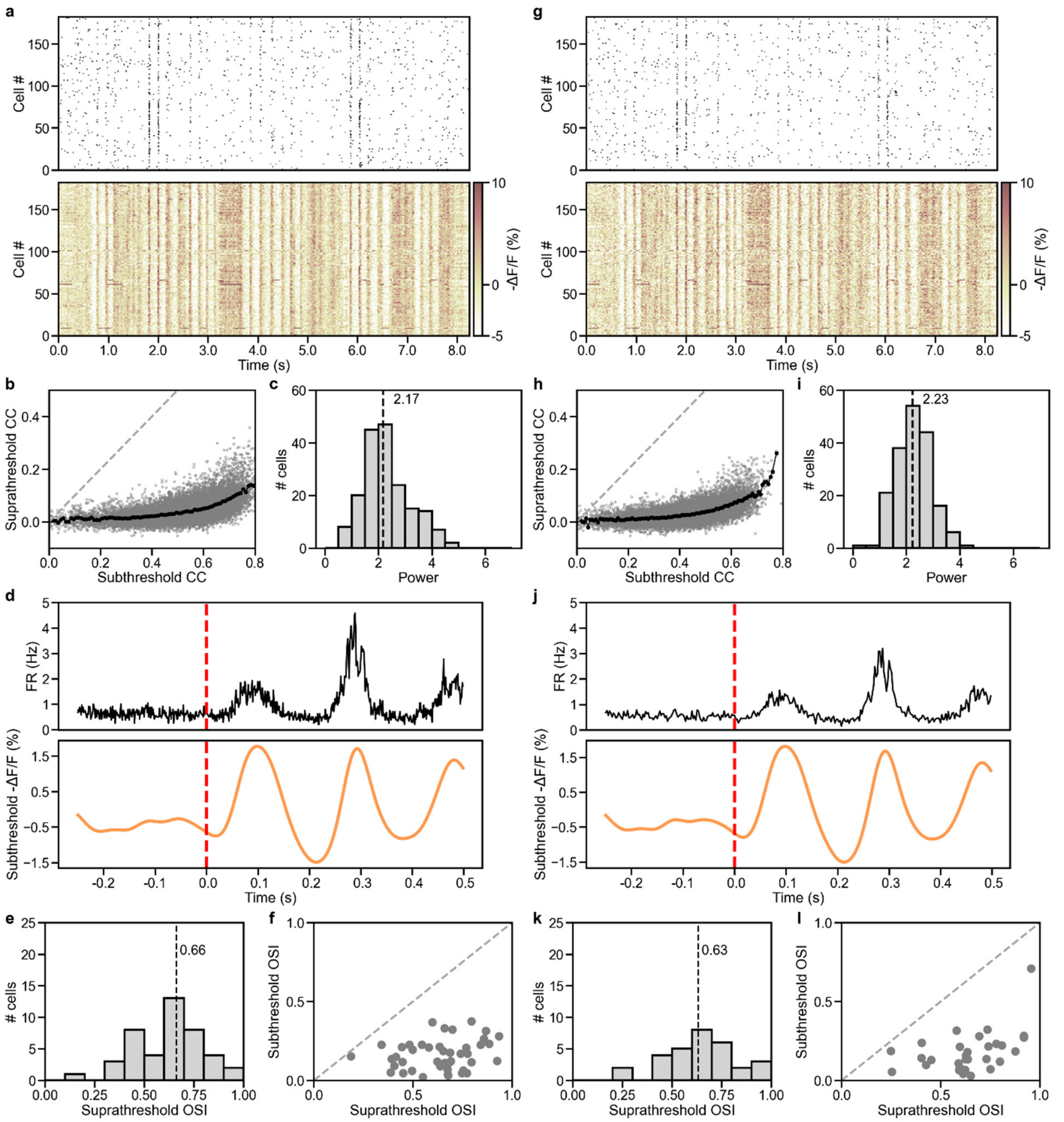

**Extended Figure 14. Analysis comparison of an 800-Hz voltage imaging dataset and its 2× down-sampled (to 400 Hz) data.** (**a-f**) Analyses of 800-Hz data (1.25 ms/frame; same as **Fig. 2**). (**g-l**) Analyses of down-sampled data (2.5 ms/frame). (**a, g**) Raster plots of (top) spikes and (bottom) subthreshold -ΔF/F traces of 182 neurons during a representative 8.25-s trial. (**b, h**) Suprathreshold versus subthreshold pairwise correlation coefficients (CC, gray dots) for 16,471 cell pairs from the 182 neurons in **a**. Black dots: binned averages with 0.01 bin size for subthreshold CC. Dashed gray line: reference line with slope = 1. (**c**, **i**) Histogram of fitted power-law exponent values of 182 neurons. Dashed line: median. (**d**, **j**) Average firing rate (black curve, top panel) and subthreshold -ΔF/F (orange curve, bottom panel) relative to the onset of drifting grating stimuli (red dashed line) extracted from 19,008 voltage traces of 88 neurons exhibiting visually evoked activity across 27 trials × 8 drifting grating orientations. (**e**, **k**) Histogram of suprathreshold orientation-selectivity index (OSI) of 43 orientation selective (OS) neurons in **e** and 30 OS neurons in **k**. Dashed line: median. (**f**, **l**) Scatter plot of subthreshold OSI vs. suprathreshold OSI for 43 OS neurons in **f** and 30 OS neurons in **l**. Dashed line: reference line with slope = 1.

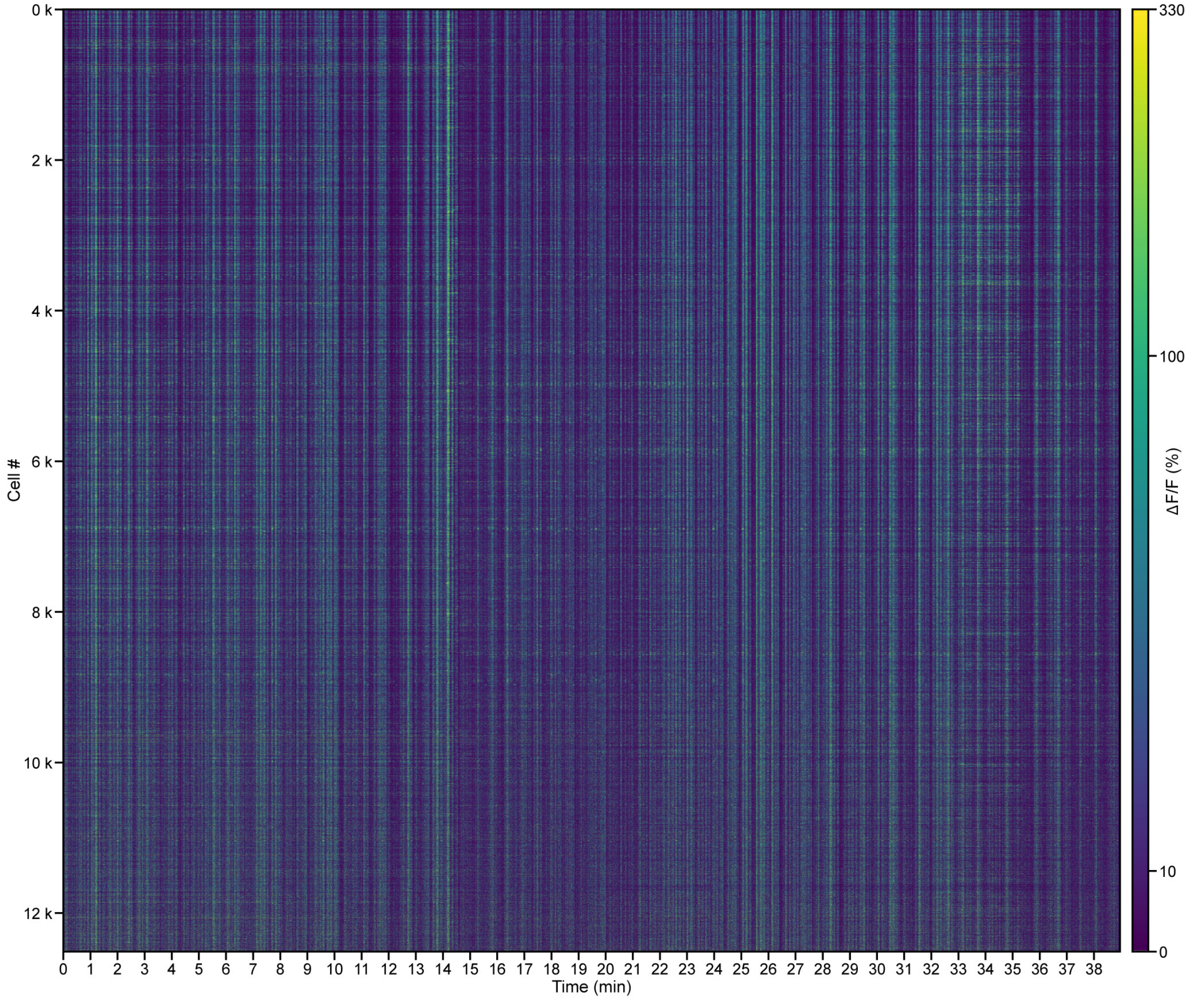

**Extended Data Figure 15. Simultaneously recorded ΔF/F calcium traces from 12,511 neurons during 36 64-s trials.** Same data as **Fig. 3**.

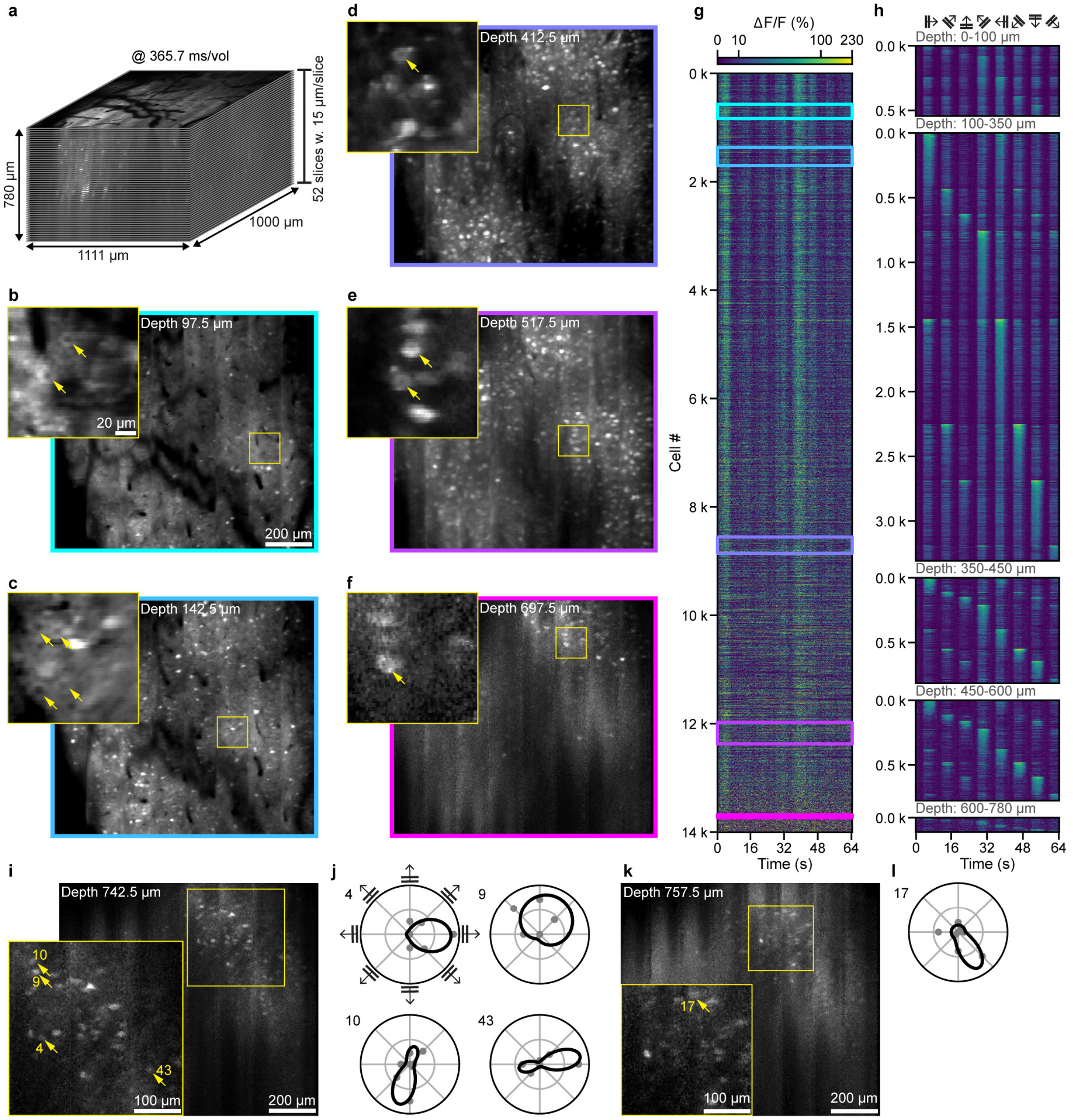

**Extended Data Figure 16. Large-scale *in vivo* volumetric imaging of calcium activity through the depth of visual cortex.** (**a**) An 1,111 μm × 1,000 μm × 780 μm volume in the visual cortex of a wildtype mouse with virally induced expression of GCaMP6s imaged at 366 ms/vol. Voxel size: 1.4 μm × 2 μm × 15 μm. 920 nm excitation at 140 mW post 16× NA 0.8 objective. (**b-f**) Example images at depths of 97.5 μm, 142.5 μm, 412.5 μm, 517.5 μm and 697.5 μm. Insets: zoomed-in views of areas in yellow boxes. Arrows: neurons showing nucleus-excluded GCaMP6s expression. (**g**) Simultaneously recorded ΔF/F calcium traces from 14,005 neurons during a 64-s trial. Colored boxes: calcium traces from neurons of depths with matched colors in **b-f**. (**h**) Trial-averaged calcium traces from 5,555 neurons with visually evoked activity, grouped by depth 0-100 μm, 100-350 μm, 350-450 μm, 450-600 μm, and 600-780 μm. (**i,k**) Images acquired at 742.5 μm and 757.5 μm depth. Insets: zoomed-in views of areas in yellow boxes. Arrows: OS neurons. (**j,l**) Polar-plot tuning curves of calcium responses of OS neurons in **i** and **k**, respectively. Dots: measured responses; curves: double-Gaussian fit.

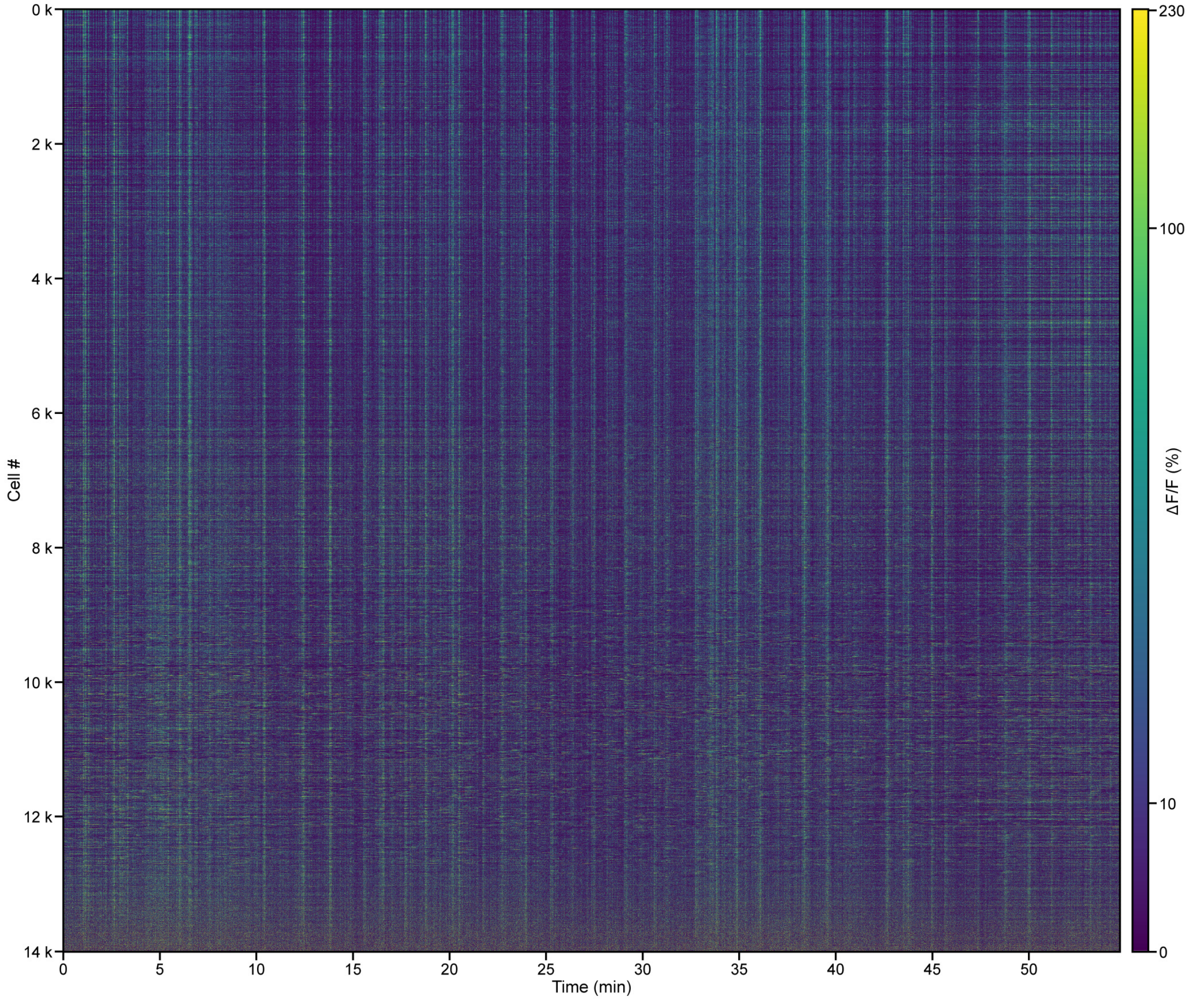

**Extended Data Figure 17. Simultaneously recorded ΔF/F calcium traces from 14,005 neurons in during 51 trials.** Same data as **Extended Data Fig. 16**.

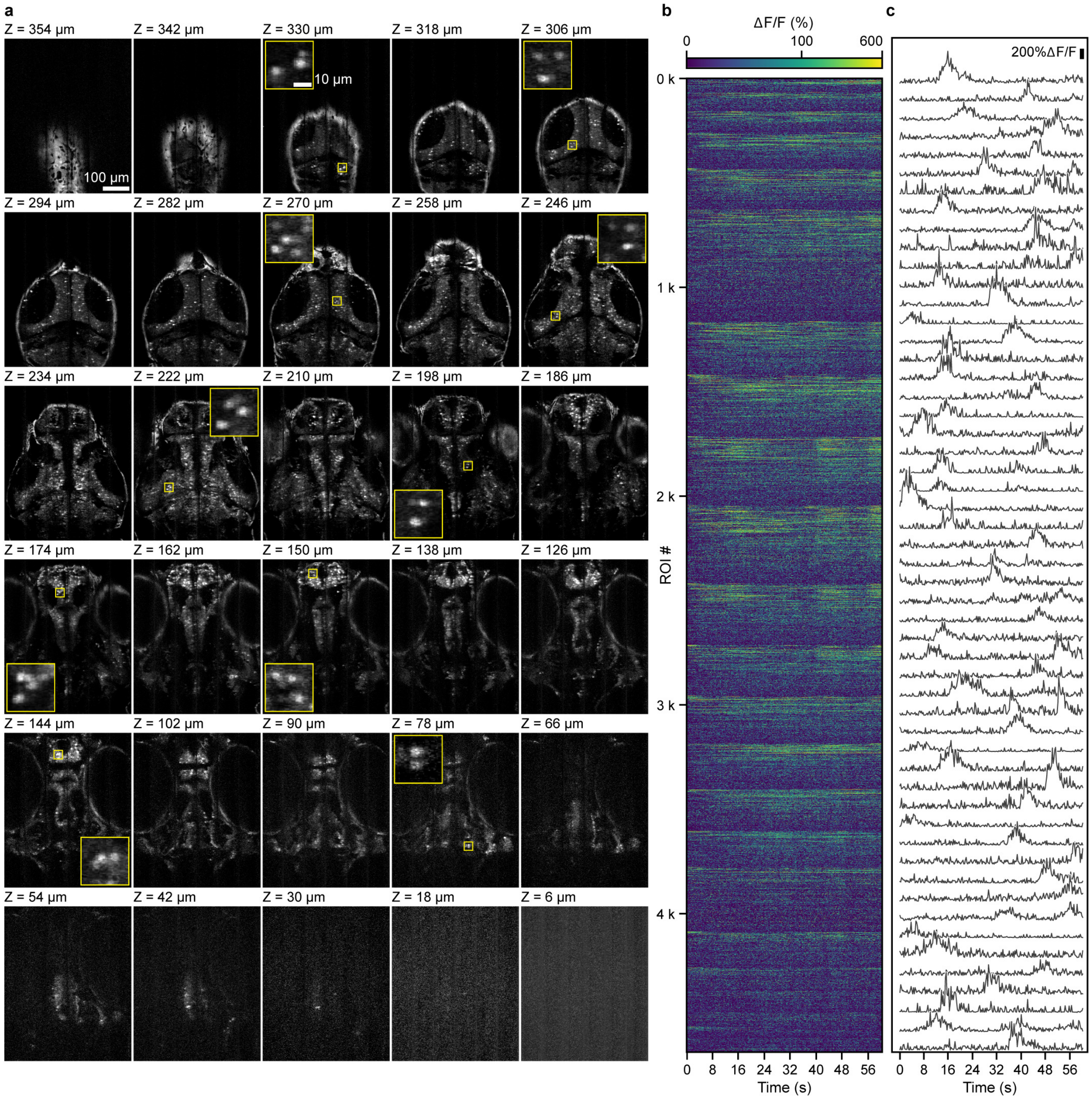

**Extended Data Figure 18. 3.8 Hz whole-brain imaging of spontaneous calcium activity in a transgenic zebrafish larva.** (**a**) XY images from a 480 µm × 585 µm × 360 µm volume of a larval Tg[Elavl3:H2B-GCaMP6s] zebrafish brain imaged at 262 ms/vol. Voxel size: 0.8 µm × 0.65 µm × 12 µm. 920 nm excitation at 78 mW post 25×1.05 NA objective. Insets: zoomed-in views of areas in yellow boxes. (**b**) Simultaneously recorded ΔF/F calcium traces from 4,718 spontaneously active neurons. (**c**) Representative calcium traces.

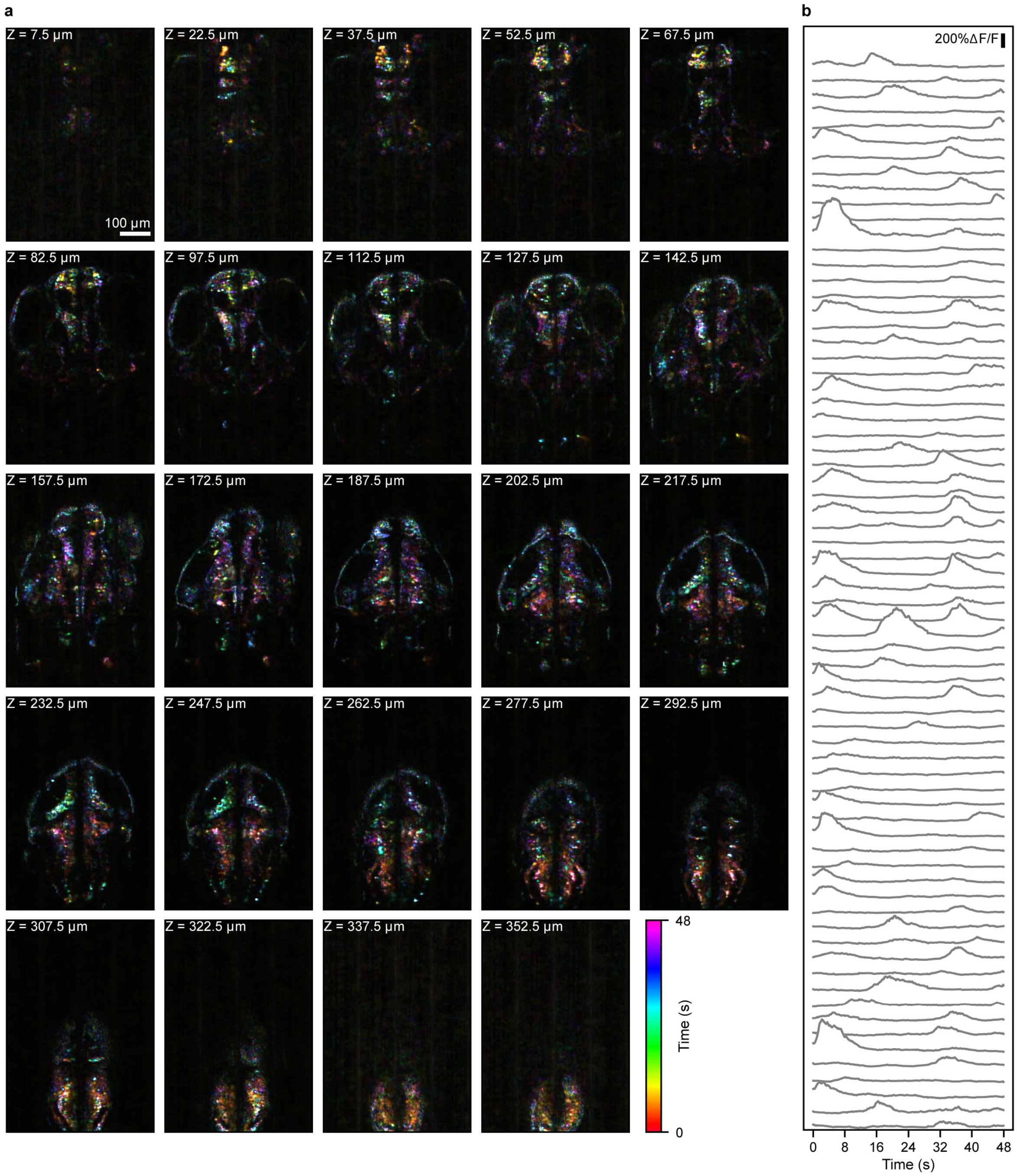

**Extended Data Figure 19. 10.4 Hz whole-brain imaging of spontaneous calcium activity in a transgenic zebrafish larva.** (**a**) XY images from a 555.6 µm × 800 µm × 360 µm volume of a larval Tg[Elavl3:H2B-GCaMP6s] zebrafish brain imaged at 96 ms/vol. A deep-learning-based denoising method (DeepCADRT) was applied to the volumetric imaging data. Color coding by temporal projection indicates the time period with high activity. Voxel size: 1.4 µm × 1.6 µm × 15 µm. 920 nm excitation at 121 mW post 16× 0.8 NA objective. (**b**) Representative calcium traces of neurons.

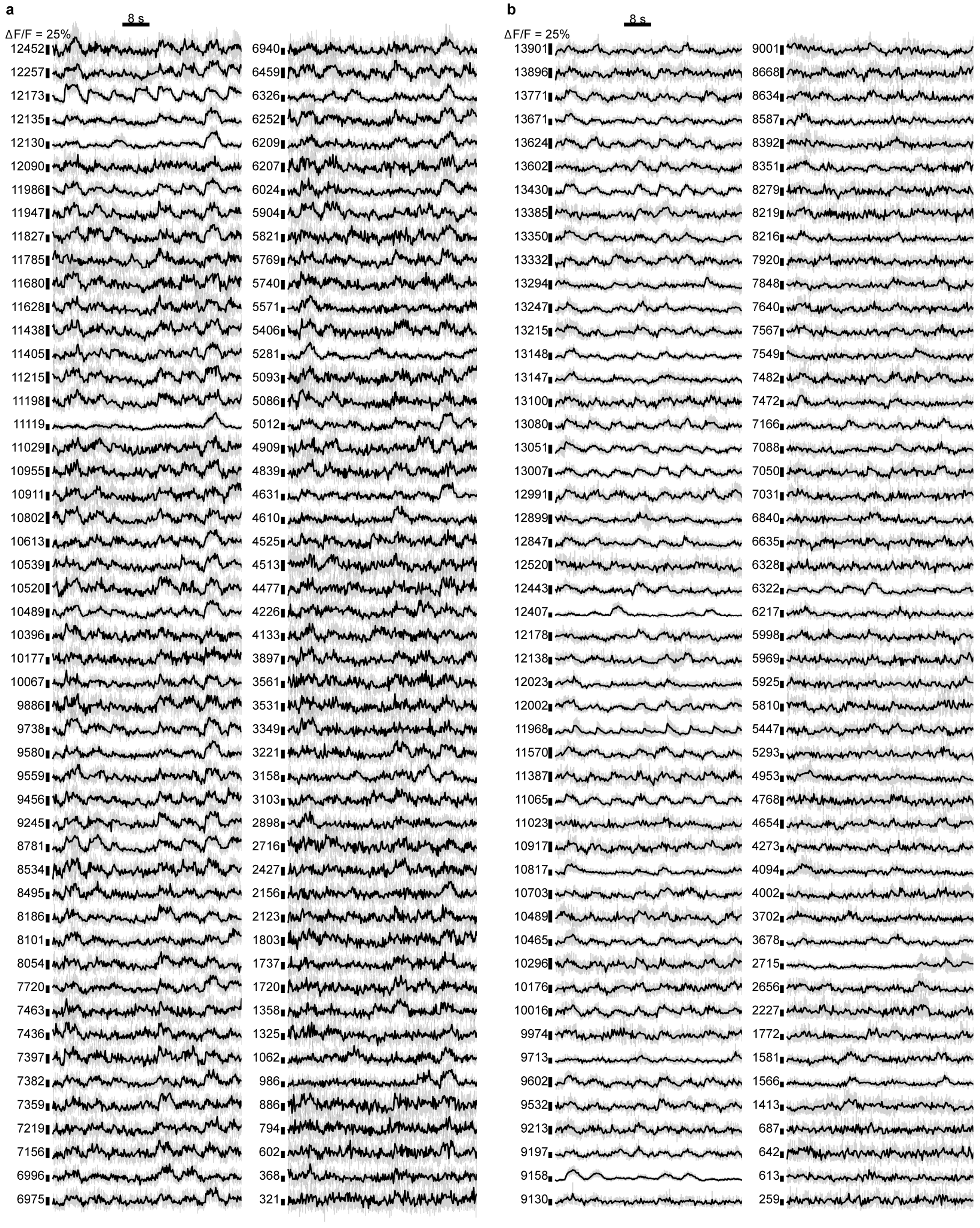

**Extended Data Figure 20. Example 10-trial-averaged ΔF/F calcium traces from calcium imaging datasets.** Example neurons were randomly selected from the visually evoked neurons across all depth ranges within the imaging volume of **Fig. 3** and **Extended Data Fig. 16**, which was achieved by sampling neurons from a discrete uniform distribution encompassing all visually evoked neurons (implemented by the Python randint() function). Numbers: Cell numbers for selected neurons. Black curves: 10-trial averaged ΔF/F calcium traces. Gray region: standard deviation.

**Supplementary Notes**

### 1. Additional characterization of volumetric imaging via continual Z movement of microscope objective

For volumetric imaging, FACED 2.0 uses a fast piezo stage to continuously oscillate the 16× 0.8-NA objective axially. To confirm that the continual Z movements during image acquisition did not generate in-frame motion artifacts, we compared 3D image stacks of static fluorescent fiber structures acquired at different rates (**Supplementary Figs. 1,2**).

For ground-truth measurements, we stepped and stopped the objective piezo at a Z position and acquired 100 XY images of 1,110 µm × 1,000 µm FOVs at 5.3 ms/frame; we repeated this at 40 Z positions with 10 µm Z-step size (**Supplementary Fig. 1**) or at 52 Z positions with 15 µm Z-step size (**Supplementary Fig. 2**) to cover the same volumes as in **Fig. 3** and **Extended Data Fig. 16**, respectively. The averages of the 100 XY images at each Z position are used as the ground-truth volumetric datasets (left panels, **Supplementary Figs. 1,2**).

We then imaged the same structures by continuously moving the objective in Z to achieve the same volumetric rates as in **Fig. 3** (259.7 ms/volume by scanning the objective piezo continuously at 1,887 µm /s) and **Extended Data Fig. 16** (365.7 ms/volume by scanning the objective piezo continuously at 2,830 µm/s). We acquired 100 volumes and compared the volume-averaged XY images at the same depths (right panels, **Supplementary Figs. 1,2**) with those from the ground-truth datasets.

Visual inspection indicates that the image stacks acquired by these two methods are highly similar. To quantify the similarity between the two stacks, we calculated the normalized cross-correlation coefficients ($\rho_{XY}$), which measures how well the images match, with higher values indicating greater similarity:

$$\rho_{XY} = \frac{\frac{1}{NML}\sum_{i=1}^{N}\sum_{j=1}^{M}\sum_{k=1}^{L}(X_{ijk}-\mu_X)(Y_{ijk}-\mu_Y)}{\sigma_X\sigma_Y}$$

Here, $X_{ijk}$ and $Y_{ijk}$ are the pixel values at position (i, j, k) in the volumetric (X) and ground-truth stacks (Y), respectively. *N*, *M*, and *L* represent the number of pixels along the X, Y, and Z axes. $\mu_X$ and $\mu_Y$ are the means, and $\sigma_X$ and $\sigma_Y$ are the standard deviations of pixel values within the entire image stacks:

$$\mu_X = \frac{1}{NML}\sum_{i=1}^{N}\sum_{j=1}^{M}\sum_{k=1}^{L}(X_{ijk}),\ \mu_Y = \frac{1}{NML}\sum_{i=1}^{N}\sum_{j=1}^{M}\sum_{k=1}^{L}(Y_{ijk})$$

$$\sigma_X = \sqrt{\frac{1}{NML}\sum_{i=1}^{N}\sum_{j=1}^{M}\sum_{k=1}^{L}(X_{ijk}-\mu_X)^2},\ \sigma_Y = \sqrt{\frac{1}{NML}\sum_{i=1}^{N}\sum_{j=1}^{M}\sum_{k=1}^{L}(Y_{ijk}-\mu_Y)^2}$$

We found $\rho_{XY} = 98\%$ for data in **Supplementary Fig. 1** and $\rho_{XY} = 95\%$ for data in **Supplementary Fig. 2**, demonstrating a high level of agreement between the stack acquired through continuous Z scanning and the ground-truth stack acquired through step-and-stop Z scanning.

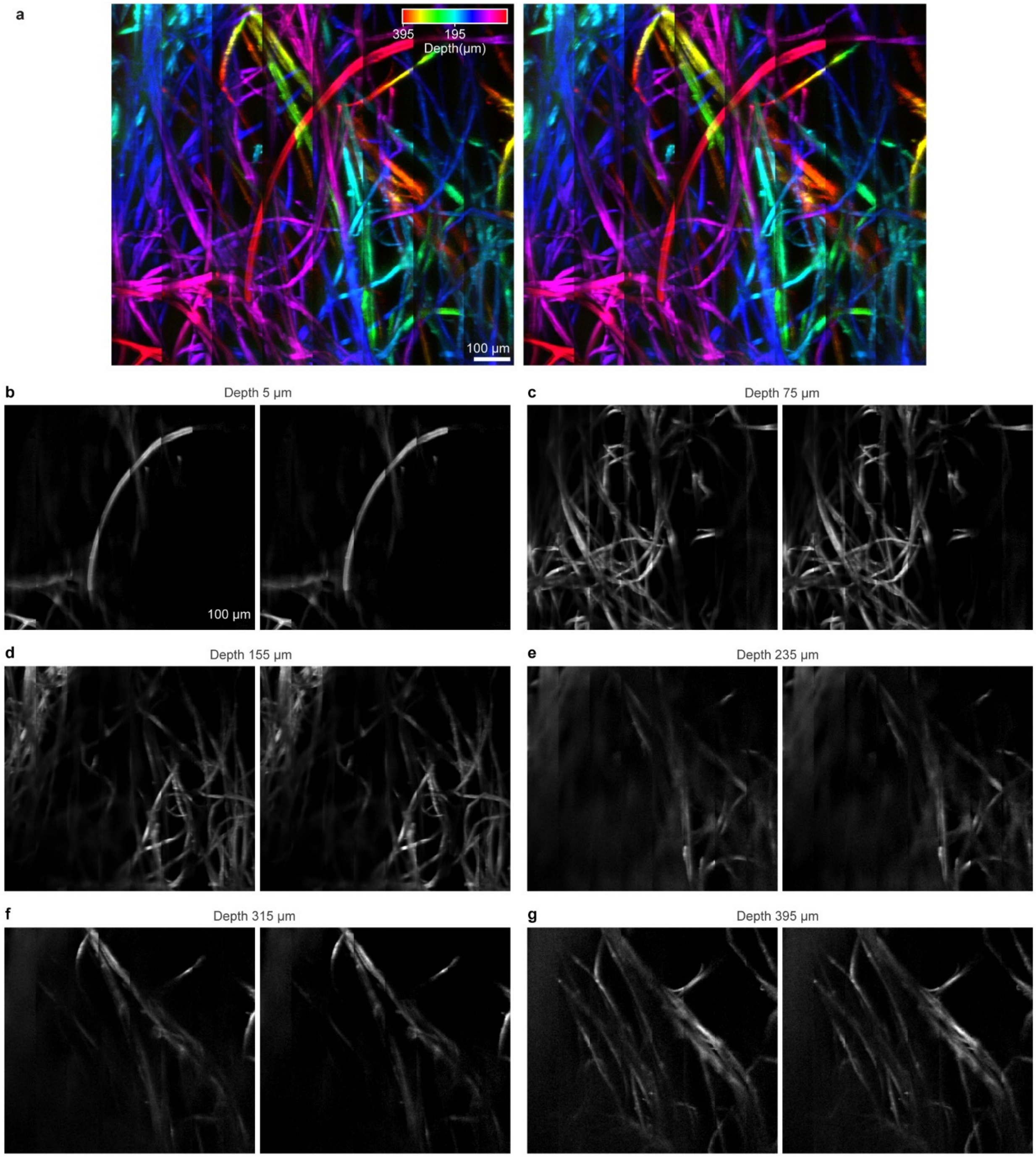

**Supplementary Figure 1. FACED volumetric imaging accurately captures 3D structures in a 1,111 µm × 1,000 µm × 400 µm volume at 259.7 ms/vol without in-frame motion artifacts.** Comparison of ground-truth (left, acquired by keeping the objective lens stationary at each depth) and volumetric (right, acquired by continuous Z scanning) images of fluorescent fibers. (**a**) Maximum intensity projection of 3D fiber structures (color-coded by depth) across the entire imaging volume for both ground-truth (left) and volumetric (right) datasets. (**b-g**) Representative ground-truth (left) and volumetric (right) images at depths of 5 µm, 75 µm, 155 µm, 235 µm, 315 µm, and 395 µm. Voxel size: 1.4 µm × 2 µm × 10 µm. 1035 nm excitation at 21 mW post 16× 0.8-NA objective.

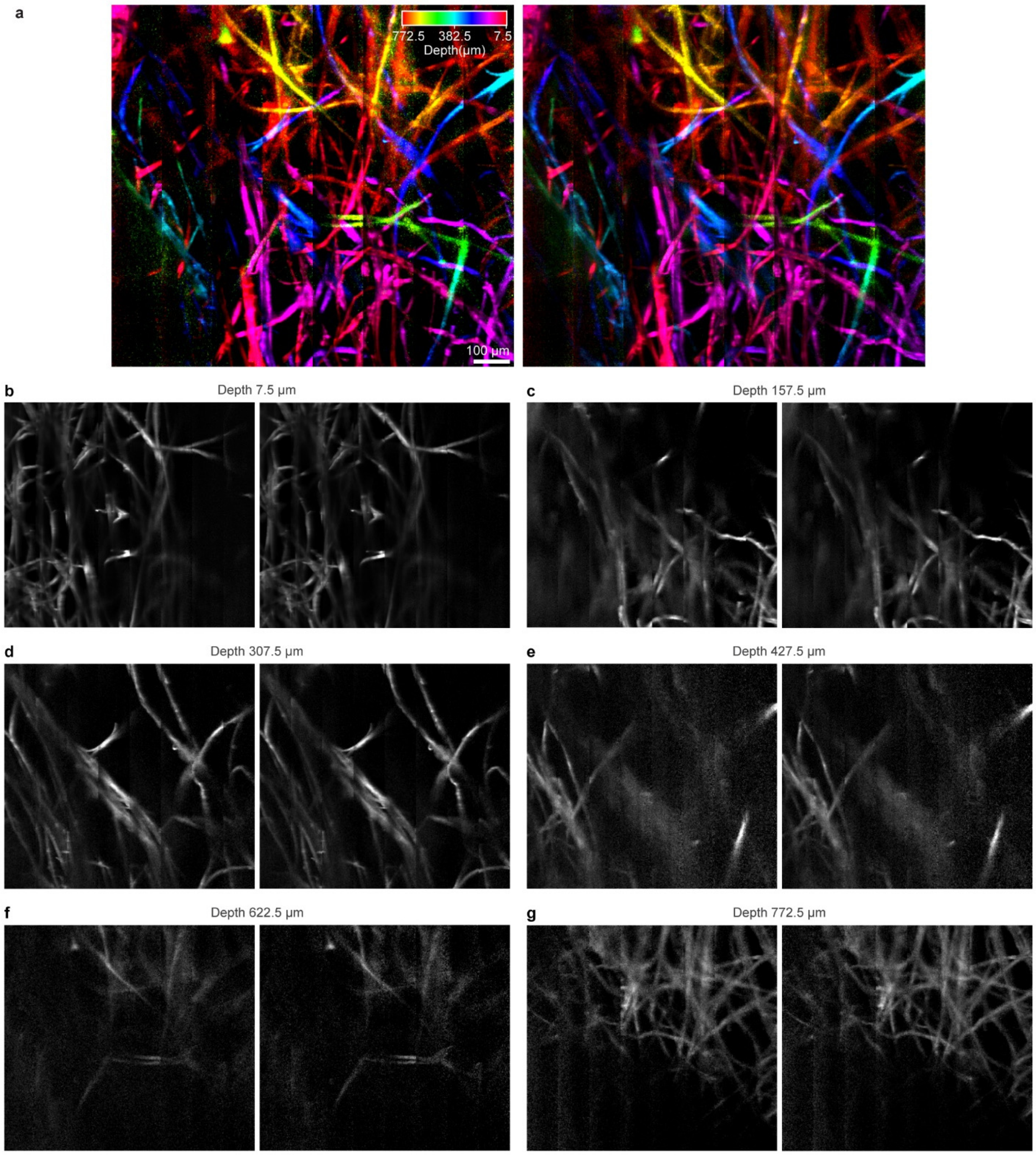

**Supplementary Figure 2. FACED volumetric imaging accurately captures 3D structures in a 1,111 μm × 1,000 μm × 780 μm volume at 365.7 ms/vol without in-frame motion artifacts.** Comparison of ground-truth (left, acquired by keeping the objective lens stationary at each depth.) and volumetric (right, acquired by continuous Z scanning) images of fluorescent fibers. (**a**) Maximum intensity projection of 3D fiber structures (color-coded by depth) across the entire imaging volume for both ground-truth (left) and volumetric (right) datasets. (**b-g**) Representative ground truth (left) and volumetric (right) images at depths of 7.5 μm, 157.5 μm, 307.5 μm, 427.5 μm, 622.5 μm, and 772.5 μm. Voxel size: 1.4 μm × 2 μm × 15 μm. 1035 nm excitation at 21 mW post 16× 0.8-NA objective.

### 2. Additional analysis on spike detection pipeline

#### 2.1 Comparison with a SNR-thresholding criterion

We carried out additional analysis on the two-step spike detection pipeline described in **Methods**: First VolPy[1], a template-matching based method, was used to detect spikes from voltage traces. VolPy's spike detection leverages signal information across multi frames (thus utilizing photons across multiple frames), allowing for more accurate spike detection than simple thresholding of SNR. Subsequently, we implemented a standard deviation (SD) based rejection step, which discards VolPy-detected spikes with peak -ΔF/F values lower than the mean+2.5×SD within a 20-frame rolling window. This rejection step minimizes false spikes.

Here, we calculated the signal-to-noise ratio (SNR), defined as $-\Delta F/\sqrt{F}$, for spikes detected in **Fig. 2** and **Extended Data Fig. 11** using the above spike detection criterion (**Supplementary Fig. 3**). We found that 96.2 % and 75.1% of neurons, respectively, had their spike SNR values greater than 7.5.

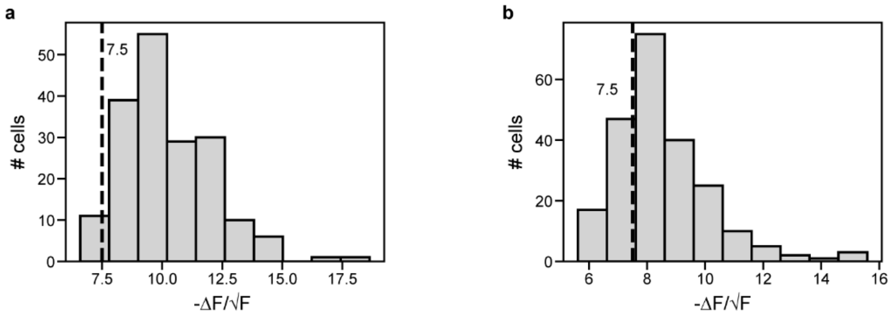

**Supplementary Fig. 3. Histograms of the average -ΔF/√F of detected spikes for 182 neurons in (a) Fig. 2 and (b) 225 neurons in Extended Data Fig. 11.** Dashed line: -ΔF/√F = 7.5.

We then reanalyzed the voltage traces from **Fig. 2** and **Extended Data Fig. 11** using a $-\Delta F/\sqrt{F}$ thresholding criterion for spike detection, which we previously employed[2]. Here a spike was detected when the SNR value was greater than 7.5. This criterion produced quantitatively similar population activity characteristics to those in **Fig. 2** and **Extended Data Fig. 11** (**Supplementary Figs. 4**,**5**).

---

[1] Cai, C. et al. VolPy: Automated and scalable analysis pipelines for voltage imaging datasets. PLoS Comput Biol 17, (2021).

[2] Wu, J. *et al.* Kilohertz two-photon fluorescence microscopy imaging of neural activity in vivo. *Nat Methods* **17**, 287–290 (2020).

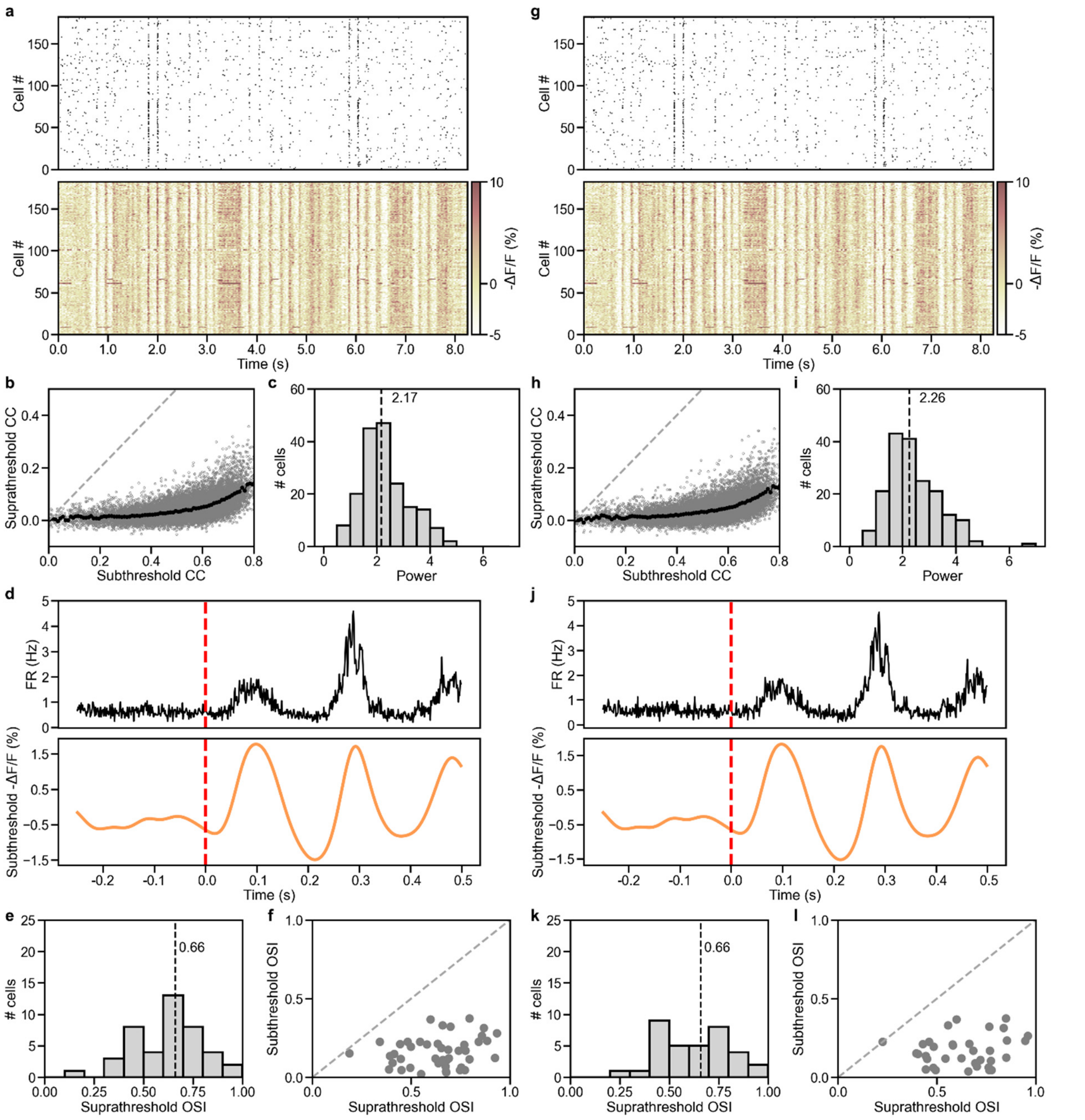

**Supplementary Fig. 4. Comparison of voltage imaging analysis results of the dataset in Fig. 2, using two spike detection methods: our method (left) and -ΔF/√F > 7.5 threshold-based detection (right).** (**a, g**) Raster plots of (top) spikes and (bottom) subthreshold -ΔF/F traces of 182 neurons during a representative 8.25-s trial. (**b, h**) Suprathreshold versus subthreshold pairwise correlation coefficients (CC, gray dots) for 16,471 cell pairs from the 182 neurons. Black dots: binned averages with 0.01 bin size for subthreshold CC. Dashed gray line: reference line with slope = 1. (**c**, **i**) Histogram of fitted power-law exponent values of 182 neurons. Dashed line: median. (**d**, **j**) Average firing rate (black curve, top panel) and subthreshold -ΔF/F (orange curve, bottom panel) relative to the onset of drifting grating stimuli (red dashed line). **d** is computed from 19,008 voltage traces from 88 neurons exhibiting visually evoked activity across 27 trials × 8 drifting grating orientations. **j** is computed from 16,416 voltage traces from 76 neurons exhibiting visually evoked activity across 27 trials × 8 drifting grating orientations. (**e**, **k**) Histogram of suprathreshold orientation-selectivity index (OSI) of 43 orientation selective (OS) neurons in **e** and 35 OS neurons in **k**. Dashed line: median. (**f**, **l**) Scatter plot of subthreshold OSI vs. suprathreshold OSI for 43 OS neurons in **f** and 35 OS neurons in **l**. Dashed line: reference line with slope = 1.

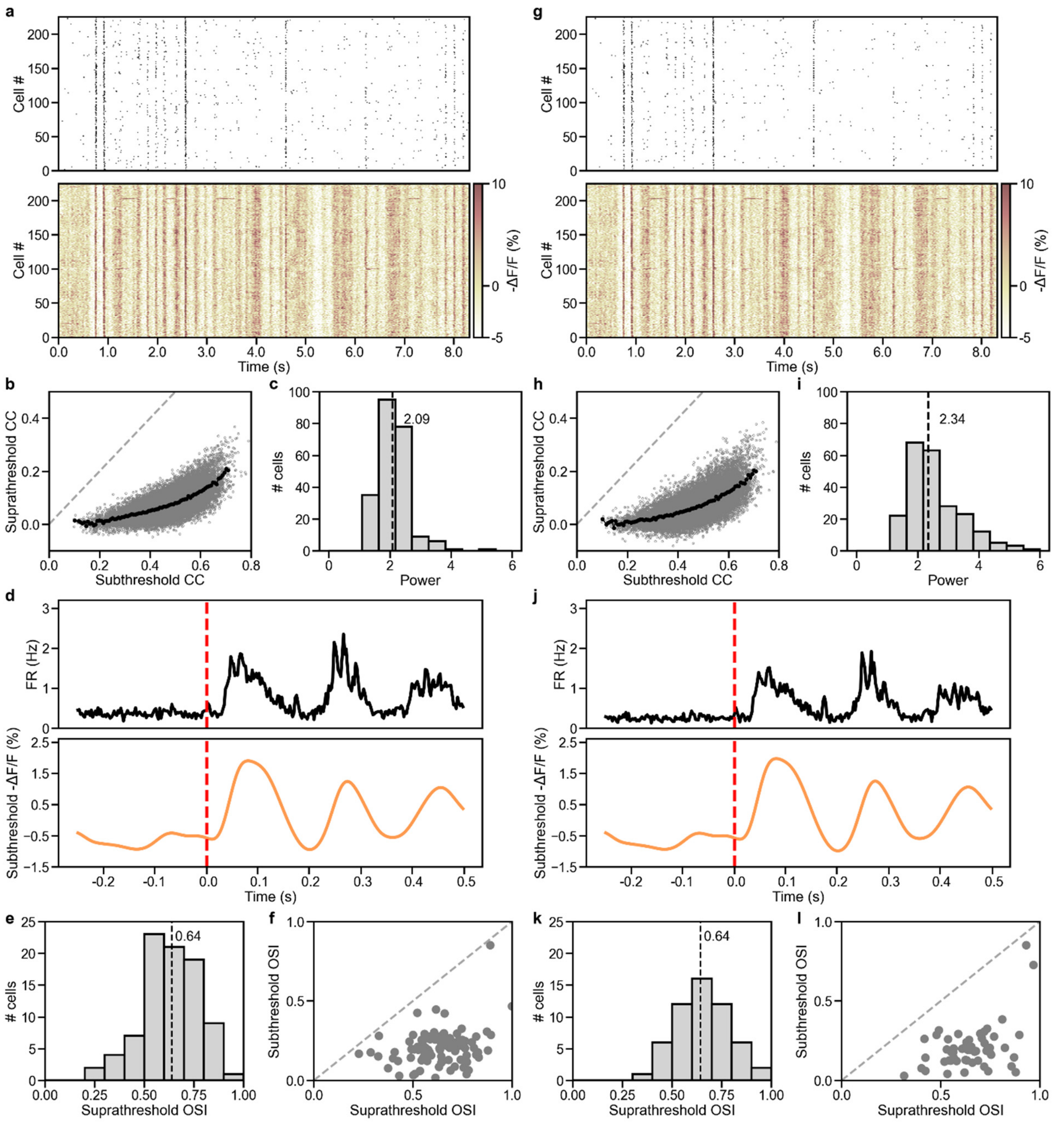

**Supplementary Figure 5. Comparison of voltage imaging analysis results from the dataset in Extended Data Fig. 11, using two spike detection methods: our method (left) and $-\Delta F/\sqrt{F} > 7.5$ threshold-based detection (right).** (**a, g**) Raster plots of (top) spikes and (bottom) subthreshold -ΔF/F traces of 182 neurons during a representative 8.32-s trial. (**b, h**) Suprathreshold versus subthreshold pairwise correlation coefficients (CC, gray dots) for 25,200 cell pairs from the 225 neurons. Black dots: binned averages with 0.01 bin size for subthreshold CC. Dashed gray line: reference line with slope = 1. (**c**, **i**) Histogram of fitted power-law exponent values of 225 neurons. Dashed line: median. (**d**, **j**) Averaged firing rate (black curve, top panel) and subthreshold -ΔF/F (orange curve, bottom panel) relative to the onset of drifting grating stimuli (red dashed line) computed from 32,240 voltage traces from 155 neurons exhibiting visually evoked activity across 26 trials × 8 drifting grating orientations. (**e**, **k**) Histogram of suprathreshold orientation-selectivity index (OSI) of 86 orientation selective (OS) neurons in **e** and 55 OS neurons in **k**. Dashed line: median. (**f**, **l**) Scatter plot of subthreshold OSI vs. suprathreshold OSI for 86 OS neurons in **f** and 55 OS neurons in **l**. Dashed line: reference line with slope = 1.

### 2.2 Evaluation of spike detection pipeline using simulated data

We further validated the accuracy of our spike detection pipeline using simulating voltage traces. The simulated traces contain supra- and subthreshold -ΔF/F activity similar to the *in vivo* data (at 1.25 ms time step size, **Supplementary Fig. 6**). The simulated suprathreshold spikes had 20% -ΔF/F and 2 ms FWHM. For simulated subthreshold activity, we set the mean peak fluctuation amplitude to be 8% ΔF/F. Poisson noise, corresponding to the photon count per frame, was then added to the ground-truth traces to approximate photon noise (**Supplementary Fig. 6a**).

We then processed these simulated traces with our spike detection pipeline and calculated the true positive rates (the percentage of ground-truth spikes that were detected by our pipeline) and the false positive rates (the number of incorrectly detected spikes divided by the total number of time points of the recording) at different photon counts (**Supplementary Fig. 6b,c**). As expected, higher photon counts are associated with more accurate spike detection. At the average photon counts for neurons in Fig. 2 (316.2 photons per ROI, **Supplementary Table 5**), our method achieved >90% true positive rates and ~0.04% false positive rates (i.e., ~4 false spikes in 10,000 time points or a 12.5-s-long recording).

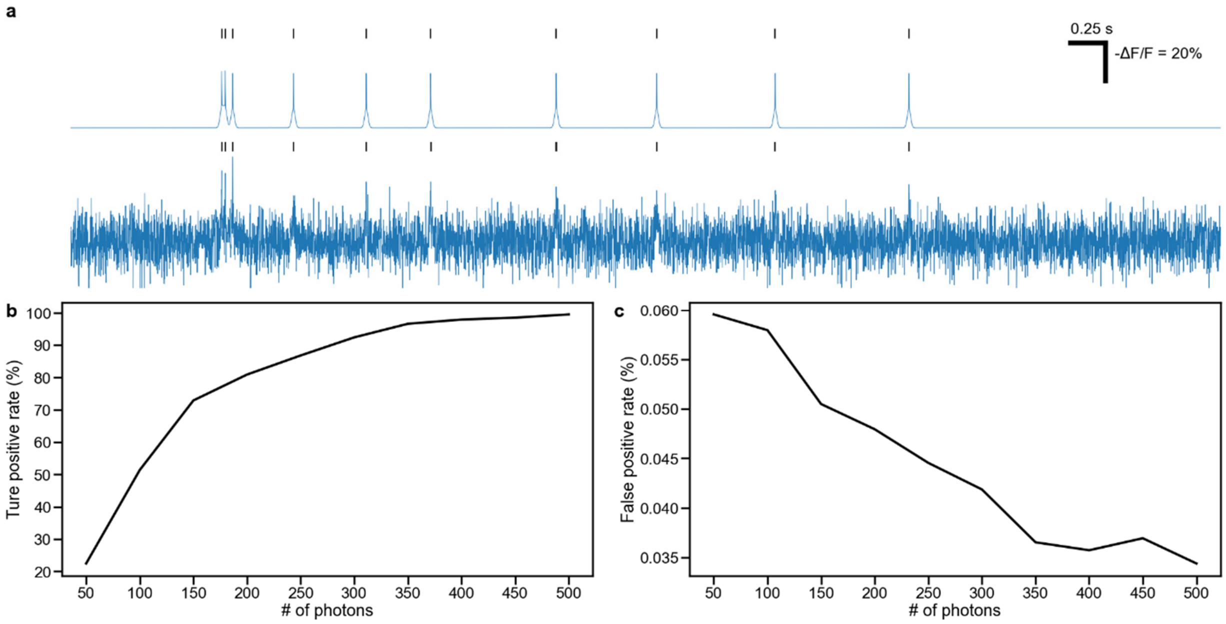

**Supplementary Fig. 6. Validation of the spike detection method on simulated voltage traces.** (**a**) Top trace: an example ground-truth voltage trace. Top black ticks: ground-truth spikes. Bottom trace: corresponding noise-added voltage trace with a SNR from 150 photons/frame. Bottom black ticks: spikes detected from noise-added voltage trace. (**b**) True positive rate of spike detection across different SNR conditions (SNR is quantified by number of photons per frame) (**c**) False positive rate of spike detection across different SNR conditions (SNR is quantified by number of photons per frame). For **b** and **c**, data point for each SNR condition is computed from 1000 simulated noisy traces with the number of spikes randomly selected (1 to 50) and randomly positioned within the trace duration (**Supplementary Codes**).

### 3. Voltage imaging at 800 Hz versus 400 Hz

We carried out additional analysis to determine whether ~400 Hz frame rate is sufficient for revealing most population voltage activity characteristics. We down-sampled the 800 Hz data in **Fig. 2** (by keeping alternate data points in time) to 400 Hz and then carried out the same analyses on correlated activity, timing and magnitude of visually evoked response, input-output function, and population orientation tuning properties (**Extended Data Fig. 14**).

Unsurprisingly, given the much slower variation of subthreshold activity with time, properties related to subthreshold voltage activity alone do not show perceivable differences between the original 800 Hz data and the down-sampled 400 Hz data (lower panels, **Extended Data Fig. 14a,d,g,j**). Measured spike rate is lower in the down-sampled 400-Hz data than in 800-Hz data (upper panels, **Extended Data Fig. 14d,j**), likely because the slower sampling rate misses APs. However, both datasets have similar timing and temporal variation of firing rates (upper panels, **Extended Data Fig. 14a,d,g,j**).

Activity correlation across the population is insensitive to frame rate except for neuron pairs with highest correlated spiking activity (**Extended Data Fig. 14b,h**). Distributions of the exponent of the input-output function also have very similar median values (2.17 vs 2.23), with the main difference being that 800 Hz sampling revealed more highly nonlinear neurons (e.g., those with exponents larger than 3) than the down-sampled 400 Hz data (**Extended Data Fig. 14c,i**). These differences are likely caused by 400 Hz data tending to miss more APs during burst firing.

Although the numbers of neurons that were found to have visually evoked AP firing are the same for the 800 Hz and 400 Hz data (88 out of the total 182 neurons), the number of neurons with suprathreshold orientation selectivity is higher for the 800 Hz data (43 neuros at 800 Hz vs. 30 neurons at 400 Hz). However, OSI distributions are again highly similar between the two datasets (median: 0.66 vs. 0.63; **Extended Data Fig. 14e,f,k,l**).

The above analysis indicates that imaging JEDI-2P-kv-expressing neurons at ~400 Hz can inform on neuronal activity with similar accuracy as imaging at 800 Hz for most application scenarios. 400 Hz data become less accurate only when high-fidelity recording of high-frequency firing is required.

### 4. Discussion on potential contamination due to fluorescence lifetime in activity imaging

In the current FACED implementation, consecutive FACED foci are delayed by 1 ns. For fluorophores with lifetimes substantially longer than 1 ns, fluorophores excited by one FACED focus may emit their fluorescence photons during the data acquisition periods of subsequent pixels. Because most fluorescent proteins have lifetimes around 3 ns[3], this would lead to a practical decrease of the lateral resolution along the FACED linescan direction, as can be seen in structural images of dendritic spines (**Fig. 1d, Extended Data Fig. 4**). However, as detailed below, this lifetime effect minimally affects population activity imaging of neuronal cell bodies (e.g., data in **Fig. 2,3**).

FACED is distinct from other 2PFM techniques that utilize temporal multiplexing for fast scanning. Those techniques utilize much longer delays (for example, 3 ns delay[4], 6.7 ns delay[5], and 8 ns delay[6]), because consecutive laser pulses and thus pixels sample distinct structures tens of microns apart along either lateral or axial direction. For such implementations, it is essential to have large temporal delay in order to avoid crosstalk of signals from distinct structures. In contrast, neighboring pulses in FACED linescans probe structures that are 0.8 μm (for voltage imaging) or 1.4 μm (for calcium imaging) apart along the X direction. Therefore, neighboring pixels mostly sample the cell body of the same neurons and contain similar activity information.

When analyzing somatic voltage and calcium activity, ROIs are selected to include the entire somata. The values of pixels inside each ROI are then averaged to obtain its activity trace, because these pixels contain the same information and pixel averaging gives rise to higher SNR. At our pixel sizes, each cell body is represented by on average 244 pixels for voltage imaging and 100 pixels for calcium imaging. With the FACED foci scanning from left to right, the only pixels that may be affected are those at the left edge of the ROI, with their signal contaminated by pixels to the left of the ROI. There are on average 17 pixels (for voltage imaging) or 9 pixels (for calcium imaging) at the left edge of the ROIs, corresponding to only 6.8% and 8.5% of total pixels for voltage and calcium imaging ROIs, respectively.

In the case of voltage imaging, due to soma-targeting of JEDI-2P-kv, neuropil to the left of the ROIs had much lower brightness than the ROIs themselves. For calcium imaging, expression level in the neuropil was higher, but neuropil activity is still of much smaller magnitude than somatic activity[7]. Together, these factors further minimize contamination in the activity traces due to fluorescence lifetime.

That crosstalk due to fluorescence lifetime minimally contaminates the activity traces of individual neurons is supported by quantitative analysis: If there is substantial crosstalk, then two neurons that are neighbors along the FACED scan direction (X axis) would have more correlated activity (because fluorescence from the neuron on the left would contaminate the signal of the neuron on the right) than two neurons that are neighbors along the Y axis. We calculated the Pearson correlation coefficients (PCC) between the activity traces of the nearest neighbors along X direction ("NNx", blue data points, **Supplementary Fig. 7**) and

---

along Y direction ("NNy", blue data points, **Supplementary Fig. 7**) for both voltage and calcium imaging data.

As shown in **Supplementary Fig. 7**, the PCCs of activity traces of the closest neighbors along X axis and along Y axis exhibited similar distributions. One-sided t-tests on whether the PCC distribution along X is greater than the PCC distribution along Y have p-values of 0.95 (**Supplementary Fig. 7a**), 0.51 (**Supplementary Fig. 7b**), 0.73 (**Supplementary Fig. 7c**), and 0.70 (**Supplementary Fig. 7d**), indicating that there is no statistically significant difference between these distributions. Therefore, activity traces of neighboring neurons along X do not exhibit a higher correlation than those along Y, indicating that activity imaging does not suffer from contamination due to fluorescence lifetime.

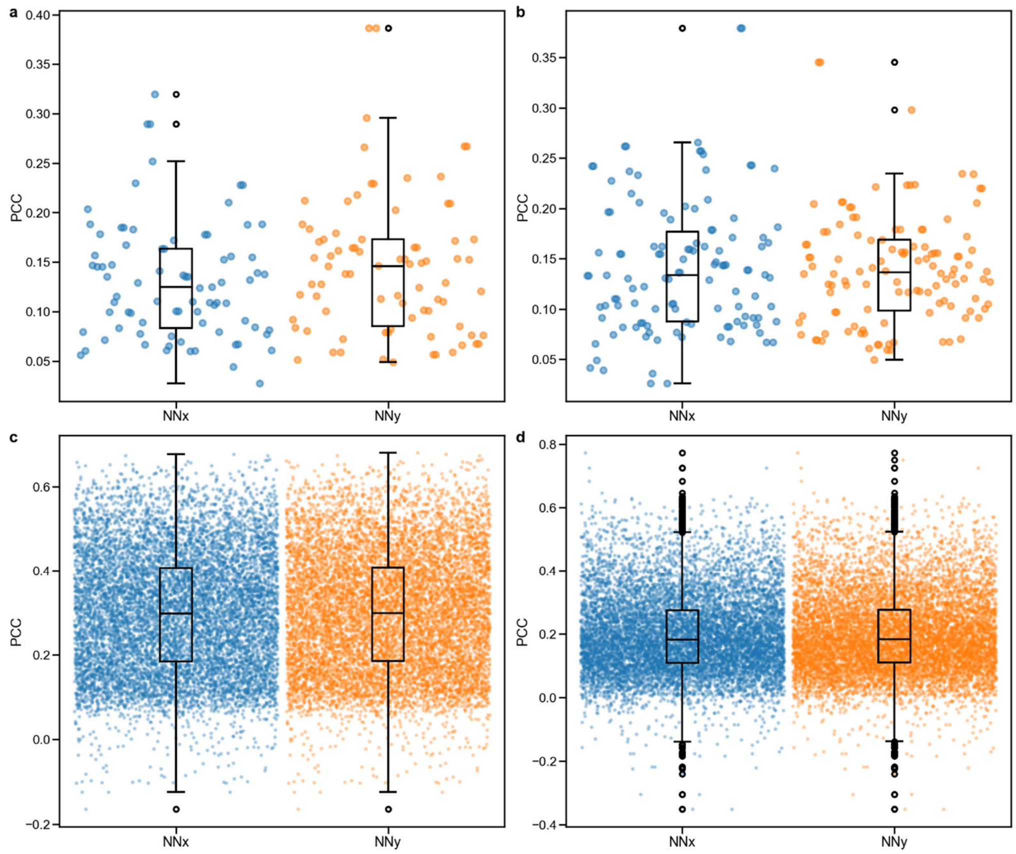

**Supplementary Figure 7. Comparison of Pearson correlation coefficients (PCCs) between activity traces of neurons that are nearest neighbors along X axis (direction with potential contamination due to fluorescence lifetime) and Y axis (direction without the possibility of contamination).** (**a,b**) Scatter and box-and-whisker plots of PCCs of voltage traces from **Fig. 2** and **Extended Data Fig. 11**, respectively. (**c,d**) Scatter and box-and-whisker plots of PCCs of calcium traces from Fig. 3 and Extended Data Fig. 16, respectively. Blue dots and "NNx": PCC between two neurons that are nearest neighbors along X axis; Orange dots and "NNy": PCC between two neurons that are nearest neighbors along Y axis. Box-and-whisker plots: box represents interquartile range (IQR) extending from the first quartile to the third quartile; horizontal line indicates median; whisker extends to the most extreme data points within 1.5×IQR from box edge; black circles: outliers. One-sided t-tests comparing PCC(NNx) and PCC(NNy) yields p-values of 0.92, 0.42, 0.73, and 0.70 for the data in **a**, **b**, **c**, and **d**, respectively. Each scatter plot in **a**, **b**, **c**, and **d** contains data points from 81, 118, 9575, 9638 pairs of neurons, respectively.

### 5. FACED 2.0 versus FACED 1.0

FACED 2.0 substantially enhances the imaging capabilities of FACED 2PFM by offering larger imaging FOVs and longer continuous imaging durations than FACED 1.0, in addition to having volumetric imaging capability. These improvements, achieved via both hardware and software optimization, allow for selecting imaging area/volume/throughput based on the temporal dynamics of biological events, enabling the study of larger neuronal populations over wider spatial domains and longer timescales at higher throughput than FACED 1.0.

In FACED 1.0, the FACED module was configured to generate linescans containing 80 pixels at a 2 ns inter-pixel delay, with its imaging FOV restricted to 50 µm × 250 µm (25× 1.05 NA objective). In FACED 2.0, the FACED module is configured to generate linescans containing 100 pixels at a 1 ns inter-pixel delay, with a single FACED FOVs reaching 80 µm × 585 µm (25× 1.05 NA objective) or 140 µm × 1035 µm (16× 0.8 NA objective).

A complete rewrite of the microscope control software for FACED 2.0 allows us to increase Y galvo scanning to 833 Hz (1.67 kHz frame rate) over 585 µm for 25× 1.05-NA objective and 1,035 µm for 16× 0.8 NA objective, a substantial improvement over from FACED 1.0 with 500 Hz Y galvo scanning over 250 µm for 25× 1.05-NA objective. Adding an additional X galvo that scans along the FACED linescan direction in FACED 2.0 allows tiling FACED FOVs to further enlarge the area that can be scanned at high speed. For example, using the 16× 0.8 NA objective, an area of 1111 µm × 1000 µm can be scanned at 192 Hz.

The control software also synchronizes 2D FACED imaging with continuous axial scanning of the microscope objective, enabling volumetric imaging. The large FOV and high frame rate of FACED 2.0 enable calcium activity of >10,000 neurons to be collected over large volumes at true cellular resolution (e.g., 1111 µm × 1000 µm × 780 µm at 365.7 ms/volume).

FACED 1.0 utilized a PXIe-5160 digitizer (National Instruments), which limits the maximal continuous imaging duration to 6 s. FACED 2.0 uses a high-speed digitizer with onboard FPGA (ADQ7-DC, Teledyne SP Devices). Together with a custom multi-thread data acquisition program that optimizes data transfer and storage, it enables continuous data streaming, limited in its duration only by disk storage size.

**Supplementary Table 1. Comparison of 2PFM techniques**

| Technique | Pixels/voxels per second | Lateral resolution: X × Y | Axial (Z) resolution | Max depth in mouse brain in vivo | 2D imaging performance: FOV @ frame rate (pixel size) | 3D imaging performance: Volume @ volume rate (voxel size) | Notes |
|---|---|---|---|---|---|---|---|
| FACED 2.0 | $1.0 \times 10^8$ [a] | 0.83 µm × 0.53 µm (1.05-NA objective)<br><br>1.27 µm × 0.76 µm (0.8-NA objective) | 2.0 µm (1.05-NA objective)<br><br>4.8 µm (0.8-NA objective) | 757 µm | 80 µm × 585 µm @ 1 kHz (0.8 µm × 0.65 µm)<br>80 µm × 585 µm @ 1.67 kHz (0.8 µm × 1.17 µm)<br><br>140 µm × 1035 µm @ 1 kHz (1.4 µm × 1.15 µm)<br><br>80 µm × 585 µm @ 1.67 kHz (0.8 µm × 2.07 µm)<br><br>160 µm × 400 µm @ 800 Hz (0.8 µm × 0.8 µm)<br><br>320 µm × 400 µm @ 385 Hz (0.8 µm × 0.8 µm)<br><br>1111 µm × 1000 µm @ 192 Hz (1.4 µm × 2 µm) | 1111 µm × 1000 µm × 780 µm @ 2.7 Hz (1.4 µm × 2 µm × 15 µm)<br><br>1111 µm × 1000 µm × 400 µm @ 3.8 Hz (1.4 µm × 2 µm × 10 µm)<br><br>480 µm × 585 µm × 360 µm @ 3.8 Hz (0.8 µm × 0.65 µm × 12 µm)<br><br>555.2 µm × 800 µm × 360 µm @ 10.4 Hz (1.4 µm × 1.6 µm × 15 µm) | Current manuscript |
| FACED 1.0[1] | $0.8 \times 10^8$ [a] | 0.82 µm × 0.35 µm | 3.0 µm | 345 µm | 50 µm × 250 µm @ 1 kHz (0.6 µm × 0.3 µm) | N/A | |

| | | | | | | | |
|---|---|---|---|---|---|---|---|
| Multifoci excitation[2] | $2.0 \times 10^9$ | 1.4 µm × 1.4 µm | 9.6 µm | 300 µm | 95,000 µm² @ 1 kHz (1024 × 196 pixels)<br><br>248,000 µm² @ ≤400 Hz (1024 × 512 pixels) | N/A | Camera detection: limited depth in opaque brains |
| Bessel focus scanning[3,4] | $7.9 \times 10^6$ | 0.3-0.7 µm | 20-160 µm | 650 µm | N/A | 180 µm × 180 µm × 160 µm @ 30 Hz (0.35 µm × 0.35 µm × 160 µm)<br><br>270 µm × 270 µm × 100 µm @ 30 Hz (0.53 µm × 0.53 µm × 100 µm)<br><br>301 µm × 450 µm × 612 µm @ 3.2 Hz (0.4 µm × 0.4 µm × 100 µm) [b]<br><br>3020 µm × 1500 µm × 600 µm @ 1 Hz (2 µm × 2 µm × 100 µm) [b] | Extended-depth excitation: requires sparse labeling |
| Light beads microscopy[5] | $1.41 \times 10^8$ | 1.2 µm × 1.9 µm | 19-30 µm | 600 µm | N/A | 600 µm × 600 µm × 500 µm @ 9.6 Hz (1 µm × 1 µm × 16 µm) [b]<br><br>900 µm × 900 µm × 500 µm @ 38 Hz (3.1 µm × 3.1 µm × 16 µm) [b]<br><br>3000 µm × 5000 µm × 500 µm @ 4.7 Hz (4.2 µm × 5 µm × 16 µm) [b]<br><br>5400 µm × 6000 µm × 500 µm @ 2.2 Hz (4.2 µm × 5 µm × 16 µm) [b] | Resolution values from Extended Data Fig. 4i,j |

| | | | | | | | |
|---|---|---|---|---|---|---|---|
| Light sculpting based on temporal focusing[6] | $2.3 \times 10^6$ | 5 µm × 5 µm | 10 µm | 500 µm | 500 µm × 500 µm @ 158 Hz (4.2 µm × 4.2 µm)<br><br>500 µm × 500 µm @ 10 Hz Dual plane (4.2 µm × 4.2 µm) | 500 µm × 500 µm × 500 µm @ 3 Hz (4.2 µm × 4.2 µm × 11.9 µm)<br><br>500 µm × 500 µm × 200 µm @ 5.7 Hz (4.2 µm × 4.2 µm × 9.5 µm) | |
| Hybrid Multiplexed Sculpted Light (HyMS) Microscopy[7] | $1.2 \times 10^7$ | 5 µm × 5 µm | 15 µm | 700 µm | N/A | 690 µm × 675 µm × 600 µm @ 16.7 Hz (5 µm × 5 µm × 15 µm)<br><br>1000 µm × 1000 µm × 600 µm @ 5.1 Hz (5 µm × 5 µm × 15 µm)<br><br>765 µm × 665 µm × 640 µm @ 4.3 Hz (5 µm × 5 µm × 13.3 µm) | Data reported are for 2PFM modality |
| Scan multiplier unit (SMU)[8,9] | $6.6 \times 10^7$ | 2.2 µm × 1.9 µm | 8.2 µm | 540 µm | 200×200 µm @ 1kHz (0.78 µm × 0.78 µm) | N/A | |
| Optical gearbox[10] | $7.2 \times 10^7$ | 0.49 µm × 0.48 µm | 2.60 µm | 460 µm | 700 µm ×560 µm @ 50 Hz (0.52 µm × 0.52 µm)<br><br>344 µm ×274 µm @ 200 Hz (0.52 µm × 0.51 µm)<br><br>350 µm × 44 µm @ 1 kHz (0.58 µm × 0.59 µm)<br><br>155 µm × 124 µm @ 200 Hz (0.23 µm ×0.23 µm)<br><br>251 µm × 454 µm @ 200 Hz (0.38 µm × 0.43 µm)<br><br>196 µm × 196 µm @ 200 Hz (0.29 µm × 0.38 µm) | 2 × 155 µm ×124 µm × 40 µm @ 10 Hz (0.23 µm × 0.23 µm × 4 µm) | Image FOV and pixel sizes are estimated from figures shared data |

[a] Can be increased by simply increasing laser repetition rate; See Ref. 1.

[b] Applied in a mesoscope system.

[1] Wu, J. et al. Kilohertz two-photon fluorescence microscopy imaging of neural activity in vivo. Nat Methods 17, 287–290 (2020).
[2] Zhang, T. et al. Kilohertz two-photon brain imaging in awake mice. Nat Methods 16, 1119–1122 (2019).
[3] Lu, R. et al. Video-rate volumetric functional imaging of the brain at synaptic resolution. Nat Neurosci 20, 620–628 (2017).
[4] Lu, R. et al. Rapid mesoscale volumetric imaging of neural activity with synaptic resolution. Nat Methods 17, 291–294 (2020).
[5] Demas, J. et al. High-speed, cortex-wide volumetric recording of neuroactivity at cellular resolution using light beads microscopy. Nat Methods 18, 1103–1111 (2021).
[6] Prevedel, R. et al. Fast volumetric calcium imaging across multiple cortical layers using sculpted light. Nat Methods 13, 1021–1028 (2016).
[7] Weisenburger, S. et al. Volumetric Ca2+ Imaging in the Mouse Brain Using Hybrid Multiplexed Sculpted Light Microscopy. Cell 177, 1050-1066.e14 (2019).
[8] Xiao, S., Davison, I. & Mertz, J. Scan multiplier unit for ultrafast laser scanning beyond the inertia limit. Optica 8, 1403 (2021).
[9] Xiao, S., Giblin, J. T., Boas, D. A. & Mertz, J. High-throughput deep tissue two-photon microscopy at kilohertz frame rates. Optica 10, 763 (2023).
[10] Lin, J., Cheng, Z., Yang, G. & Cui, M. Optical gearbox enabled versatile multiscale high-throughput multiphoton functional imaging. Nat Commun 13, (2022).

### Supplementary Table 2: Parameters for 2D imaging with FACED 2.0

| Figure | Sample | FOV dimension (X µm × Y µm) | Pixel size (X µm × Y µm) | Rescaled pixel size for display (µm) | Frame rate (ms/frame ) | Post-objective power @ wavelength | Imaging Depth (µm) | Microscope objective | Filter* | # of frame average for display |
|---|---|---|---|---|---|---|---|---|---|---|
| Fig. 1d | Thy1-GFP line M mouse | 46 x 64 (Cropped from a larger image) | 0.8 × 0.2 | 0.2 | 9.3 | 150 mW @ 1035 nm | 36±2.5 (0.5 µm /step) | 25× 1.05 NA | 525/50 BP | 400 |
| Fig. 1e-g | Wild-type mouse injected with Rhodamine B | 1111 × 1000 | 1.4 × 2.0 | 1.4 | 5.3 | 215 mW @ 1035 nm | 50 | 16× 0.8 NA | 680 SP | 6000 |
| Fig. 2 | Wild-type mouse transfected with JEDI-2P-kv voltage indicator | 160 × 400 | 0.8 × 0.8 | 0.8 | 1.25 | 163 mW @ 1035 nm | 132 | 25× 1.05 NA | 680 SP | 6600 |
| Extended Data Fig. 4 | Thy1-GFP line M mouse | 92 × 128 (Cropped from larger raw image) | 1.4 × 0.4 | 0.4 | 9.3 | 165 mW @ 1035 nm | 29.5±5 (0.5 µm /step) | 16× 0.8 NA | 525/50 BP | 400 |
| Extended Data Fig. 5 | Thy1-GFP line M mouse | 640 × 585 | 0.8 × 0.65 | 0.65 | 9.3 | 140 mW @ 1035 nm | 50-800 | 25× 1.05 NA | 525/50 BP | 4000 |
| Extended Data Fig. 6 | Thy1-GFP line M mouse | 1111 × 1035 | 1.4 × 1.15 | 1.15 | 9.3 | 170 mW @ 1035 nm | 50-800 | 16× 0.8 NA | 525/50 BP | 4000 |
| Extended Data Fig. 7 | Wild-type mouse injected with Rhodamine B | 1111 × 1000 | 1.4 × 2.0 | 1.4 | 5.3 | 208 mW @ 1035 nm | 0-750 | 16× 0.8 NA | 680 SP | 6000 |
| Extended Data Fig. 11 | Wild-type mouse transfected with JEDI-2P-kv voltage indicator | 320 × 400 | 0.8 × 0.8 | 0.8 | 2.6 | 164 mW @ 1035 nm | 140 | 25× 1.05 NA | 680 SP | 3200 |

*525/50 BP: band-pass filter with 525 nm center wavelength and 50 nm bandwidth; 680 SP: short-pass filter with 680 nm cutoff wavelength.

**Supplementary Table 3: Parameters for 3D imaging with FACED 2.0**

| Figure | Sample | Volume dimension (X µm × Y µm × Z µm) | Voxel size (X µm × Y µm × Z µm) | Rescaled Voxel size for display (XY µm × Z µm) | Volume rate (ms/vol) | Post-Objective power @ wavelength | Imaging Depth (µm) | Microscope objective | Filter* | # of volumes average for display |
|---|---|---|---|---|---|---|---|---|---|---|
| Fig. 1b | Wild-type mouse injected with Rhodamine B | 1111 × 1000 × 600 | 1.4 × 2.0 × 5.0 | 1.4 × 5.0 | 624 | 240 mW @ 1035 nm | 25-625 | 16× 0.8 NA | 680 SP | 100 |
| Fig. 3 | a Slc17a7-IRES2-Cre × Ai162D mouse | 1111 × 1000 × 400 | 1.4 × 2.0 × 10.0 | 1.4 × 10.0 | 259.7 | 102 mW @ 920 nm | 0-400 | 16× 0.8 NA | 525/50 BP | 9000 |
| Extended Data Fig. 16 | Wild-type mouse transfected with GCaMp6s | 1111 × 1000 × 780 | 1.4 × 2.0 × 15.0 | 1.4 × 15.0 | 365.7 | 140 mW @ 920 nm | 0-780 | 16× 0.8 NA | 525/50 BP | 8976 |
| Extended Data Fig. 18 | larval Tg[Elavl3:H2B-GCaMP6s] zebrafish | 480 × 585 × 360 | 0.8 × 0.65 × 12.0 | 0.65 × 12.0 | 262.4 | 78 mW @ 920 nm | 0-360 | 25×1.05 NA | 525/50 BP | 230 |
| Extended Data Fig. 19 | larval Tg[Elavl3:H2B-GCaMP6s] zebrafish | 555.6 × 800 × 360 | 1.4 × 1.6 × 15.0 | 1.4 × 15.0 | 96 | 121 mW @ 920 nm | 0-360 | 16× 0.8 NA | 525/50 BP | 500 |
| Supp. Fig. 1 | Fluorescent lens tissue fiber | 1111 × 1000 × 400 | 1.4 × 2.0 × 10.0 | 1.4 × 10.0 | 259.7 | 21 mW @ 1035 nm | 0-400 | 16× 0.8 NA | 680 SP | 100 |
| Supp. Fig. 2 | Fluorescent lens tissue fiber | 1111 × 1000 × 780 | 1.4 × 2.0 × 15.0 | 1.4 × 15.0 | 365.7 | 21 mW @ 1035 nm | 0-780 | 16× 0.8 NA | 680 SP | 100 |

*525/50 BP: band-pass filter with 525 nm center wavelength and 50 nm bandwidth; 680 SP: short-pass filter with 680 nm cutoff wavelength.

### Supplementary Table 4: Brightness comparison for FACED 2.0 figures

| Figure | Sample | Average of pixel value in the top 99% of image frame[1] | Frame/volume rate | Post-objective power @ wavelength | Imaging Depth (µm) | Microscope objective | Filter[2] | # of frame/volume average for figure display |
|---|---|---|---|---|---|---|---|---|
| Fig. 1b depth = 42.5 µm | Wild-type mouse injected with Rhodamine B | 9821 | 624 ms/vol | 240 mW @ 1035 nm | 42.5 | 16× 0.8 NA | 680 SP | 100 |
| Fig. 1b depth = 342.5 µm | Wild-type mouse injected with Rhodamine B | 1078 | 624 ms/vol | 240 mW @ 1035 nm | 342.5 | 16× 0.8 NA | 680 SP | 100 |
| Fig. 1b depth = 592.5 µm | Wild-type mouse injected with Rhodamine B | 138 | 624 ms/vol | 240 mW @ 1035 nm | 592.5 | 16× 0.8 NA | 680 SP | 100 |
| Fig. 1d | Thy1-GFP line M mouse | 1372 | 9.3 ms/frame | 150 mW @ 1035 nm | 36±2.5 (0.5 µm /step) | 25× 1.05 NA | 525/50 BP | 400 |
| Fig. 1e | Wild-type mouse injected with Rhodamine B | 7005 | 5.3 ms/frame | 215 mW @ 1035 nm | 50 | 16× 0.8 NA | 680 SP | 6000 |
| Fig. r9 | Wild-type mouse transfected with JEDI-2P-kv | 3766 | 1.25 ms/frame | 163 mW @ 1035 nm | 132 | 25× 1.05 NA | 680 SP | 6600 |
| Fig. 3b | Slc17a7-IRES2-Cre × Ai162D mouse | 842 | 259.7 ms/vol | 102 mW @ 920 nm | 105 | 16× 0.8 NA | 525/50 BP | 9000 |
| Fig. 3c | Slc17a7-IRES2-Cre × Ai162D mouse | 497 | 259.7 ms/vol | 102 mW @ 920 nm | 185 | 16× 0.8 NA | 525/50 BP | 9000 |
| Extended Data Fig. 4 | Thy1-GFP line M mouse | 314 | 9.3 ms/frame | 165 mW @ 1035 nm | 29.5±5 (0.5 µm /step) | 16× 0.8 NA | 525/50 BP | 400 |
| Extended Data Fig. 5 depth = 50 µm | Thy1-GFP line M mouse | 369 | 9.3 ms/frame | 140 mW @ 1035 nm | 50 | 25× 1.05 NA | 525/50 BP | 4000 |
| Extended Data Fig. 5 depth = 400 µm | Thy1-GFP line M mouse | 63 | 9.3 ms/frame | 140 mW @ 1035 nm | 400 | 25× 1.05 NA | 525/50 BP | 4000 |
| Extended Data Fig. 5 depth = 750 µm | Thy1-GFP line M mouse | 38 | 9.3 ms/frame | 140 mW @ 1035 nm | 750 | 25× 1.05 NA | 525/50 BP | 4000 |
| Extended Data Fig. 6 depth = 50 µm | Thy1-GFP line M mouse | 225 | 9.3 ms/frame | 170 mW @ 1035 nm | 50 | 16× 0.8 NA | 525/50 BP | 4000 |
| Extended Data Fig. 6 depth = 400 µm | Thy1-GFP line M mouse | 89 | 9.3 ms/frame | 170 mW @ 1035 nm | 400 | 16× 0.8 NA | 525/50 BP | 4000 |

| | | | | | | | | |
|---|---|---|---|---|---|---|---|---|
| Extended Data Fig. 6 depth = 750 µm | Thy1-GFP line M mouse | 29 | 9.3 ms/frame | 170 mW @ 1035 nm | 750 | 16× 0.8 NA | 525/50 BP | 4000 |
| Extended Data Fig. 7 depth = 50 µm | Wild-type mouse injected with Rhodamine B | 2641 | 5.3 ms/frame | 208 mW @ 1035 nm | 50 | 16× 0.8 NA | 680 SP | 6000 |
| Extended Data Fig. 7 depth = 400 µm | Wild-type mouse injected with Rhodamine B | 321 | 5.3 ms/frame | 208 mW @ 1035 nm | 400 | 16× 0.8 NA | 680 SP | 6000 |
| Extended Data Fig. 7 depth = 750 µm | Wild-type mouse injected with Rhodamine B | 43 | 5.3 ms/frame | 208 mW @ 1035 nm | 750 | 16× 0.8 NA | 680 SP | 6000 |
| Extended Data Fig. 11 | Wild-type mouse transfected with JEDI-2P-kv | 3303 | 2.6 ms/frame | 164 mW @ 1035 nm | 140 | 25× 1.05 NA | 680 SP | 3200 |
| Extended Data Fig. 16b | Wild-type mouse transfected with GCaMP6s | 1355 | 365.7 ms/vol | 140 mW @ 920 nm | 97.5 | 16× 0.8 NA | 525/50 BP | 8976 |
| Extended Data Fig. 16c | Wild-type mouse transfected with GCaMP6s | 1277 | 365.7 ms/vol | 140 mW @ 920 nm | 142.5 | 16× 0.8 NA | 525/50 BP | 8976 |
| Extended Data Fig. 16d | Wild-type mouse transfected with GCaMP6s | 474 | 365.7 ms/vol | 140 mW @ 920 nm | 412.5 | 16× 0.8 NA | 525/50 BP | 8976 |
| Extended Data Fig. 16e | Wild-type mouse transfected with GCaMP6s | 195 | 365.7 ms/vol | 140 mW @ 920 nm | 517.5 | 16× 0.8 NA | 525/50 BP | 8976 |
| Extended Data Fig. 16f | Wild-type mouse transfected with GCaMP6s | 34 | 365.7 ms/vol | 140 mW @ 920 nm | 697.5 | 16× 0.8 NA | 525/50 BP | 8976 |
| Extended Data Fig. 16i | Wild-type mouse transfected with GCaMP6s | 29 | 365.7 ms/vol | 140 mW @ 920 nm | 742.5 | 16× 0.8 NA | 525/50 BP | 8976 |
| Extended Data Fig. 16k | Wild-type mouse transfected with GCaMP6s | 27 | 365.7 ms/vol | 140 mW @ 920 nm | 757.5 | 16× 0.8 NA | 525/50 BP | 8976 |
| Extended Data Fig. 18a Z = 330 µm | larval Tg[Elavl3:H2B-GCaMP6s] zebrafish | 306 | 262.4 ms/vol | 78 mW @ 920 nm | 30 | 25×1.05 NA | 525/50 BP | 230 |

| | | | | | | | | |
|---|---|---|---|---|---|---|---|---|
| Extended Data Fig. 18a Z = 210 µm | larval Tg[Elavl3:H2B-GCaMP6s] zebrafish | 250 | 262.4 ms/vol | 78 mW @ 920 nm | 150 | 25×1.05 NA | 525/50 BP | 230 |
| Extended Data Fig. 18a Z = 90 µm | larval Tg[Elavl3:H2B-GCaMP6s] zebrafish | 104 | 262.4 ms/vol | 78 mW @ 920 nm | 270 | 25×1.05 NA | 525/50 BP | 230 |
| Extended Data Fig. 19 Z = 322.5 µm | larval Tg[Elavl3:H2B-GCaMP6s] zebrafish | 132 | 96 ms/vol | 121 mW @ 920 nm | 37.5 | 16× 0.8 NA | 525/50 BP | 500 |
| Extended Data Fig. 19 Z = 172.5 µm | larval Tg[Elavl3:H2B-GCaMP6s] zebrafish | 182 | 96 ms/vol | 121 mW @ 920 nm | 187.5 | 16× 0.8 NA | 525/50 BP | 500 |
| Extended Data Fig. 19 Z = 22.5 µm | larval Tg[Elavl3:H2B-GCaMP6s] zebrafish | 56 | 96 ms/vol | 121 mW @ 920 nm | 337.5 | 16× 0.8 NA | 525/50 BP | 500 |
| Supplementary Fig. 1c | Fluorescent lens tissue fiber | 20961 | 259.7 ms/vol | 21 mW @ 1035 nm | 75 | 16× 0.8 NA | 680 SP | 100 |
| Supplementary Fig. 1d | Fluorescent lens tissue fiber | 20942 | 259.7 ms/vol | 21 mW @ 1035 nm | 155 | 16× 0.8 NA | 680 SP | 100 |
| Supplementary Fig. 1g | Fluorescent lens tissue fiber | 634 | 259.7 ms/vol | 21 mW @ 1035 nm | 395 | 16× 0.8 NA | 680 SP | 100 |
| Supplementary Fig. 2b | Fluorescent lens tissue fiber | 22198 | 365.7 ms/vol | 21 mW @ 1035 nm | 7.5 | 16× 0.8 NA | 680 SP | 100 |
| Supplementary Fig. 2d | Fluorescent lens tissue fiber | 1005 | 365.7 ms/vol | 21 mW @ 1035 nm | 307.5 | 16× 0.8 NA | 680 SP | 100 |
| Supplementary Fig. 2g | Fluorescent lens tissue fiber | 481 | 365.7 ms/vol | 21 mW @ 1035 nm | 772.5 | 16× 0.8 NA | 680 SP | 100 |

[1]On average, 1 photon is equivalent to 1300 pixel value.

[2]525/50 BP: band-pass filter with 525 nm center wavelength and 50 nm bandwidth; 680 SP: short-pass filter with 680 nm cutoff wavelength.

**Supplementary Table 5: Estimated numbers of photons in activity measurement datasets**

| Figure number | Sample | 75% percentile of photon counts in a ROI per frame | 50% percentile of photon counts in a ROI per frame | 25% percentile of photon counts in a ROI per frame | Average # of photon counts per ROI per frame | Standard deviation of photon counts per ROI per frame | Average # of photon counts per pixel within each ROI per frame |
|---|---|---|---|---|---|---|---|
| Fig. 2 | Wild-type mouse transfected with JEDI-2P-kv | 414.1 | 298.7 | 203.0 | 316.2 | 138.6 | 1.4 |
| Extended Data Fig. 11 | | 401.2 | 305.3 | 224.6 | 329.6 | 144.7 | 1.4 |
| Fig. 3 | Slc17a7-IRES2-Cre-D mouse | 38.9 | 24.6 | 17.0 | 31.4 | 21.8 | 0.3 |
| Extended Data Fig. 16 | Wild-type mouse transfected with GCaMp6s | 48.8 | 29.1 | 19.0 | 38.8 | 29.2 | 0.3 |

Note: Number of photons were calculated from the baseline ($F_0$) trace of each ROI, using the conversion factor of a photon on average corresponding to 1300 in pixel value. The conversion factor was measured assuming Poisson statistics.

### Supplementary Table 6: Estimated baseline noise level for activity measurement datasets

| Figure number | Sample | Noise level for ROIs at 75% percentile of photon counts per frame | Noise level for ROIs at 50% percentile of photon counts per frame | Noise level for ROIs at 25% percentile of photon counts per frame | Noise level for average photon counts per frame |
|---|---|---|---|---|---|
| Fig. 2 | Wild-type mouse transfected with JEDI-2P-kv voltage indicator | 4.9% | 5.8% | 7.1% | 5.6% |
| Extended Data Fig. 11 | | 3.8% | 4.6% | 6.7% | 5.5% |
| Fig. 3 | Slc17a7-IRES2-Cre-D mouse | 16.1% | 20.2% | 24.3% | 17.8% |
| Extended Data Fig. 16 | Wild-type mouse transfected with GCaMp6s | 14.4% | 18.6% | 23.0% | 16.1% |

Note: The baseline noise level is estimated from $\sqrt{N}$, where N represents the photon count from **Supplementary Table 5**. This noise level estimation is then normalized to baseline photon count N to represent the noise level in $\Delta F/F$ traces.

### Supplementary Videos

**Supplementary Video 1**. Imaging cerebral blood flow over 1,111 µm × 1,000 µm at 5.3 ms/frame in the mouse cortex *in vivo* using the 16× 0.8 NA objective. Pixel size: 1.4 µm × 2 µm. Same data as in Fig. 1e. First half: 4-frame-binned image sequence; second half: raw image sequence. Video was saved at 15 frames per second (fps).

**Supplementary Video 2**. Volumetric calcium imaging over 1,111 µm × 1,000 µm × 400 µm at 259.7 ms/vol in the visual cortex of a Slc17a7-IRES2-Cre × Ai162D mouse using the 16× 0.8 NA objective. Voxel size: 1.4 µm × 1.6 µm × 15 µm. Same data as shown in Fig. 3. The image sequence at each depth was averaged across 36 trials with rolling 3-frame averaging and saved at 9 fps. First half: all imaging depths acquired; second half: imaging depths highlighted in yellow in the first half.

**Supplementary Video 3**. Volumetric calcium imaging over 1,111 µm × 1,000 µm × 780 µm at 365.7 ms/vol in the visual cortex of a wild-type mouse transfected with GCaMP6s using the 16× 0.8 NA objective. Voxel size: 1.4 µm × 2 µm × 15 µm. Same data as in Supplementary Figure 12. The image sequence at each depth was averaged across 51 trials with rolling 3-frame averaging and saved at 9 fps. First half: all imaging depths acquired; second half: imaging depths highlighted in yellow in the first half.

**Supplementary Video 4**. Volumetric calcium imaging over 480 µm × 585 µm × 360 µm at 262.4 ms/vol of a larval GCaMP6s transgenic zebrafish (Tg[Elavl3:H2B-GcaMP6s]) brain using the 25× 1.05 NA objective. Voxel size: 0.8 µm × 0.65 µm × 12 µm. Same data as in Supplementary Fig. 14. The image sequence at each depth was binned every 4 frames and saved at 15 fps. The video shows the image sequences at different depths sequentially.

**Supplementary Video 5**. Volumetric calcium imaging over 555.6 µm × 800 µm × 360 µm at 96 ms/vol of a larval GcaMP6s transgenic zebrafish (Tg[Elavl3:H2B-GCaMP6s]) brain using the 16× 0.8 NA objective. Voxel size: 1.4 µm × 1.6 µm × 15 µm. Same data as shown in Supplementary Fig. 15. The image sequence at each depth was denoised with DeepCADRT and saved at 120 fps. The video shows the image sequences at different depths sequentially.
